# High-throughput micro-patterning platform reveals Nodal-dependent dissection of peri-gastrulation-associated versus pre-neurulation associated fate patterning

**DOI:** 10.1101/465039

**Authors:** Mukul Tewary, Dominika Dziedzicka, Joel Ostblom, Laura Prochazka, Nika Shakiba, Curtis Woodford, Elia Piccinini, Alice Vickers, Blaise Louis, Nafees Rahman, Davide Danovi, Mieke Geens, Fiona M. Watt, Peter W. Zandstra

## Abstract

*In vitro* models of post-implantation human development are valuable to the fields of regenerative medicine and developmental biology. Here, we report characterization of a robust *in vitro* platform that enabled high-content screening of multiple human pluripotent stem cell (hPSC) lines for their ability to undergo peri-gastrulation-like fate patterning upon BMP4 treatment of geometrically-confined colonies and observed significant heterogeneity in their differentiation propensities along a gastrulation associable and neuralization associable axis. This cell line associated heterogeneity was found to be attributable to endogenous nodal expression, with upregulation of Nodal correlated with expression of a gastrulation-associated gene profile, and Nodal downregulation correlated with a neurulation-associated gene profile expression. We harness this knowledge to establish a platform of pre-neurulation-like fate patterning in geometrically confined hPSC colonies that arises due to a stepwise activation of reaction-diffusion and positional-information. Our work identifies a Nodal signalling dependent switch in peri-gastrulation versus pre-neurulation-associated fate patterning in hPSC cells, provides a technology to robustly assay hPSC differentiation outcomes, and suggests conserved mechanisms of self-organized fate specification in differentiating epiblast and ectodermal tissues.

## Introduction

Following implantation, embryos undergo a dramatic transformation mediated by tissue growth, cell movements, morphogenesis, and fate specifications resulting in the self-organized formation of the future body plan[1]. Post-implantation development to the neurula stage embryo is orchestrated by two vital developmentally conserved events called gastrulation and neurulation. Gastrulation is the developmental stage that segregates the pluripotent epiblast into the three multipotent germ layers, namely – the ectoderm, the mesoderm, and the endoderm[2–4]. Closely following gastrulation, the ectoderm undergoes further fate specification resulting in the patterned neural plate, neural plate border, and non-neural ectoderm regions thereby setting the stage for the onset of neurulation[5–9]. As neurulation proceeds, morphogenetic changes in these tissues result in the formation of the neural tube, the neural crest, and the epithelium respectively[10]. Initiation of the morphogenetic restructuring of the epiblast and the ectoderm occurs due to self-organized gradients of signalling molecules called *morphogens*, and morphogens belonging to the transforming growth factor beta (TGFβ) superfamily, such as bone morphogenetic proteins (BMPs) and Nodal, play vital roles in these developmental stages.

Two biochemical models, Reaction-Diffusion (RD) and Positional-Information (PI), have strongly influenced our mechanistic understanding of self-organized fate specification during embryogenesis. The RD model describes how a homogenously distributed morphogen can self-organize into a signalling gradient in a developing tissue due to the presence of an interaction network between the morphogen and its inhibitor, both of which are hypothesized to be diffusible molecules albeit with differential characteristic diffusivities[11–13]. Recent interpretations of RD have proposed that higher order (>2 molecules) network topologies can also underlie this self-organization[14]. The PI model describes how fate patterning can occur in a developing tissue due to an asymmetric morphogen distribution. The classical version of this paradigm hypothesized that the cells in the developing tissue sense the morphogen concentration in their immediate vicinity and acquire fates according to a threshold model[15,16]. Recent studies have updated this interpretation of the PI model and suggest that fates are acquired as a function of both the morphogen concentration and time of induction[17,18]. Although both RD and PI have been incredibly valuable in facilitating our comprehension of how developmental fates arise in a self-organized manner, these models are typically studied in individual signalling pathways. How multiple signalling pathways may work in concert to execute the rules specified by either RD or PI are not well understood.

Studying post-implantation developmental events, like gastrulation and neurulation, directly in human embryos would unequivocally provide the most reliable interpretations of human development. While valuable progress has been made of late in culturing human blastocysts *in vitro*[19,20], ethical concerns preclude their maintenance beyond 14 days – prior to the onset of gastrulation. On the other hand, recent studies on *in vivo* human development have provided some incredible insight into human development well into the fetal stages[21]. However, these studies are performed on specimens acquired from terminations or abortions that are typically accessible after the stages of gastrulation and neurulation have already transpired. Consequently, investigation of the mechanisms underpinning early post-implantation human embryonic development directly in human embryos is currently not possible. Nevertheless, the ability of stem cells to self-organize into structures *in vitro* that mimic aspects of post-implantation human development when provided appropriate biophysical and biochemical cues is well established[22–27]. We and others have used human pluripotent stem cells (hPSCs) to demonstrate that BMP4 treatment of geometrically confined hPSC colonies recapitulate some aspects of human peri-gastrulation-like self-organized fate patterning[24–26]. Although stem cell derived *in vitro* constructs are indisputably an incomplete representation of embryos, they can serve to provide insights into some cell organizational events that occur during the critically important post-implantation developmental stages.

In addition to providing insight into cell organizational aspects of early gastrulation-like behaviour, *in vitro* models of human development are also of great value to the field of regenerative medicine. This is because the ability of these models to specify early developmental cell fates highlights their suitability for characterization of differentiation propensities of hPSC lines. It is well known that hPSC lines – whether they are derived from embryos or from reprogrammed somatic cells – have an inherent bias to differentiate toward specific lineages[28–30]. Consequently, assays that can characterize different hPSC lines to identify these biases are of crucial importance to the field of regenerative medicine. Given the significance of developing approaches for achieve this goal, multiple assays have been established to address this need. The most prominent of these are assays like the *teratoma* assay[31], the *scorecard* assay[32], and the *pluritest[33].* Although each of these approaches have their benefits, they are either lengthy, tedious and expensive *(teratoma* and *scorecard*), or do not directly measure differentiation of the hPSC lines *(pluritest).* In contrast, the peri-gastrulation-like assay provides a quantitative measure of the generation of lineage-specific fates, is rapid and dramatically inexpensive in comparison to approaches like the teratoma assay. In addition, it is readily amenable to high-content screening. However, to capitalize on the capabilities of this assay, we require a robust micropatterning platform that readily enables high-content studies. Conventional approaches to establish micropatterning platforms have employed techniques like micro-contact printing μCP)[30,34–36], or other soft-lithography approaches[37–39]. In fact, we have previously reported a high-throughput μCP platform that enables geometric confinement in 96-well microtiter plates[30]. However, such approaches require a manual step of stamping the extracellular matrix (ECM) proteins to transfer the adhesive ‘islands’ onto the substrate of choice (glass, tissue culture polystyrene, etc.), which can result in variability in the patterning efficiency and fidelity between experiments, and between users. Additionally, they also employ soft-lithography based protocols that require costly equipment and access to clean rooms, which is detrimental to their broad utility. In contrast, techniques that employ the use of Deep UV (<200nm) light to photo-oxidize Polyethylene Glycol (PEG) coated substrates offer an attractive alternative to establish robust platforms that can enable high-content screening of hPSC lines [40,41].

Here, we report characterization of a high-content platform to produce microtiter plates that allow robust geometric confinement of a variety of adherent cell types at single cell resolution. Employing this platform, we tested the response of a panel of hPSC lines to a previously reported peri-gastrulation-like assay and observed significant variability in the induction of the Brachyury (BRA) expressing region between the lines. To probe the emergent differentiation trajectories of hPSC lines, we assessed their differentiation-associated gene expression profiles and found a switch-like response in the upregulation of gene profiles associated with either gastrulation or neurulation. This switch in gene expression showed a strong association with Nodal signalling; hPSC lines that exhibited higher levels of a gastrulation-associated gene expression profile also upregulated Nodal signalling, and those that exhibited a higher neurulation-associated gene expression profile downregulated Nodal signalling. We further validated this observation by inhibiting Nodal signalling in an hPSC line that induces gastrulation-like responses and validated the Nodal dependent switch of gastrulation versus neurulation associated gene expression switch. In addition, we report that geometrically-confined hPSC colonies induced to differentiate in the presence of BMP4 and a Nodal inhibitor undergo an RD-mediated self-organization of pSMAD1 activity and PI-mediated fate patterning into compartments that express markers like TFAP2A, SIX1, OTX2, and GATA3 indicative of differentiation toward ectodermal progenitors. We further demonstrate the ability of these progenitor regions to induce marker expression of the definitive fates of the respective compartments. Our findings provide insight into how hPSCs process information from morphogen inputs like BMP and Nodal to generate early developmental fates as output.

## Results

### A high-throughput platform for screening studies of geometrically confined cell colonies

Photo-oxidation of organic polymers like Polyethylene Glycol (PEG) – a widely reported bio-inert polymer[42], by Deep UV (DUV) light has been shown to upregulate carboxyl groups[40,41] which can be readily biofunctionalized with ECM proteins[43]. We employed this knowledge to develop a protocol to generate micropatterned, carboxyl-rich regions (**Fig. 1A**)[24]. We confirmed that incubation with Poly-L-Lysine-*grafted*-Polyethylene Glycol (PLL-g-PEG) resulted in a PEGylated surface on plasma treated borosilicate glass coverslips by probing the carbon 1s (C1s) spectra profile using X-Ray Photoelectron Spectroscopy (XPS). Consistent with previous reports[40], a peak indicating the presence of the C-O-C functional group present in PEG was detected at 286.6eV in the C1s spectrum on the PLL-g-PEG incubated glass coverslip, in addition to the peak at 285eV that was observed in the blank glass coverslip control (**Fig. 1B**). DUV treatment of the PEGylated coverslips progressively reduced the peak at 286.6eV (**Fig. 1C**) suggesting photo-oxidation mediated ablation of the PEG layer. However, we were unable to detect any carboxyl presence, which has been reported to occur at 289eV[40]. We hypothesized that the photo-oxidation of the PEG during DUV treatment reduced the polymer thickness below the detection limit of the XPS equipment employed in this study, causing the absence of the carboxyl peak in the emission spectra. Given that biochemical assays circumvent the need of minimum polymer thickness to detect the presence of functional groups of interest, we opted to employ a previously reported assay based on the preferential affinity of Toluidine blue-O (TBO) to carboxyl functional groups (see Materials and Methods for assay description)[44] and asked if DUV treatment changed the amount of TBO adsorbed onto PEGylated coverslips. Indeed, DUV treatment resulted in an increase in the amount of TBO adsorption on PEGylated coverslips, with relative levels increasing with exposure times up to 12 minutes after which the relative levels detected decreased (**Fig. 1D**). These findings indicate that, consistent with previous reports[40,45], PLL-g-PEG incubation results in PEGylation of coverslips, and that the optimal exposure time to maximize the presence of carboxyl functional groups on the PEGylated coverslips in our experimental setup was 12 minutes. To produce 96-well microtiter plates for patterned cell-culture surfaces, PEGylated large coverslips (110mmx74mm) were photopatterned by DUV exposure through Quartz photo-masks for 12 minutes and assembled to bottomless 96-well plates (**Fig. 1E**). Carboxyl groups were activated using carbodiimide chemistry[43] (**Fig. 1F**) to enable covalent attachment to primary amines on ECM molecules. This “PEG plates” platform enabled robust geometrical-confinement of a variety of cell types in colonies of a variety of shapes and sizes (**Fig. 1F, Sup. Fig. 1A-F**).

**Figure 1:**
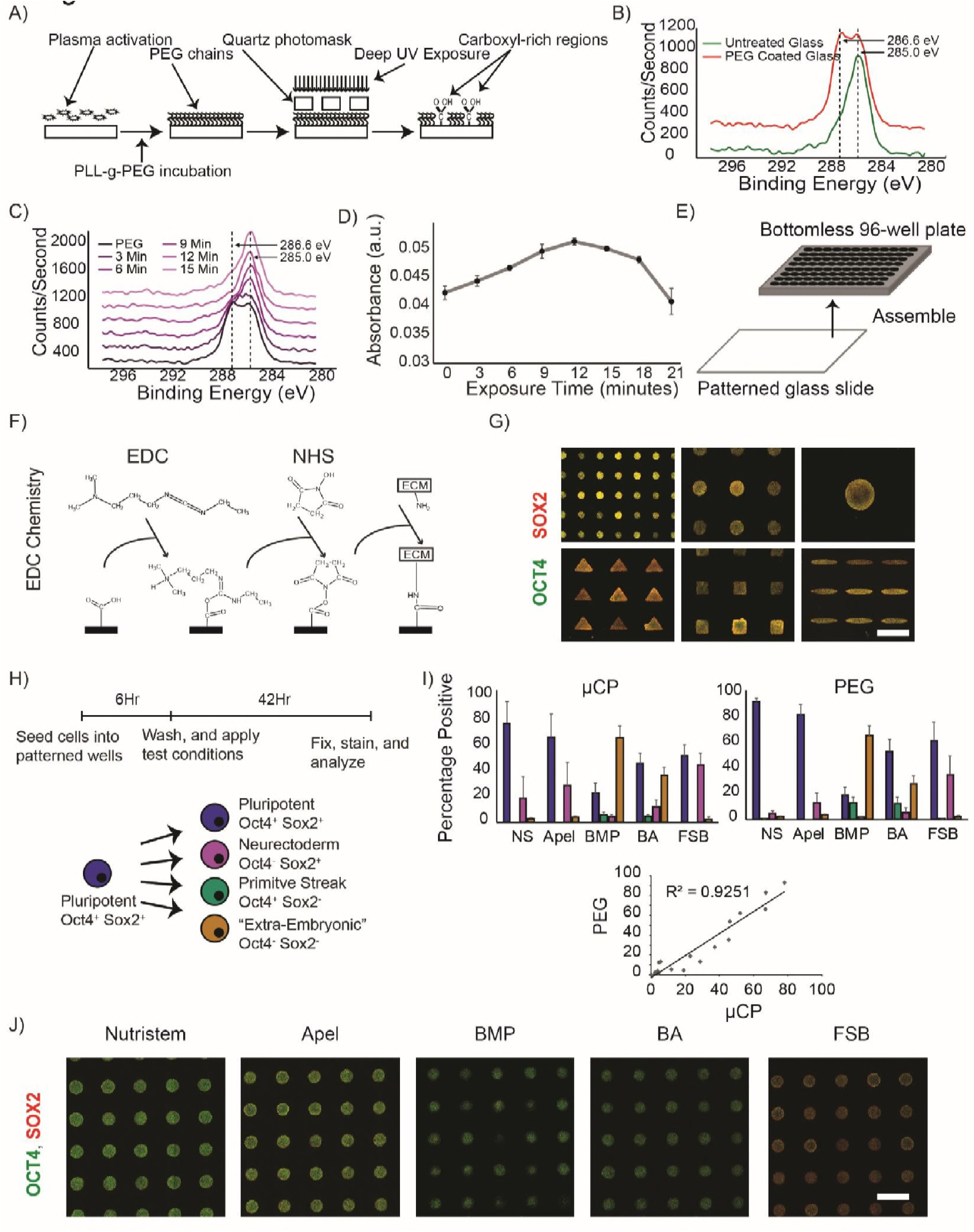
Development of Poly(ethylene glycol) based micro-patterning platform. A) Scheme of protocol for transferring carboxyl-rich micro-patterns onto glass coverslips. B-C) Carbon 1s (C1s) spectra acquired using X-Ray Photoelectron Spectroscopy. B) C1s spectra of glass coverslip incubated with PLL-g-PEG compared to blank glass coverslip. C) C1s spectra of PLL-g-PEG coated glass coverslips photoexposed to Deep-UV light for different times of exposure. Dotted lines signify binding energies associated with untreated glass (285.0eV), or presence of PEG (286.6eV). D) Line plot representation of detected absorbance at 580nm wavelength of coverslips photo-oxidized for different times of exposure indicating the relative amounts of adsorbed Toluidine Blue-O (assay details in Materials and Methods). Data represented as mean (±s.d) for three technical replicates. The assay was performed once to identify optimal exposure times for our experimental setup. E) Overview of assembly procedure to produce 96-well micro-titer plates with micro-patterned culture surface. F) Overview of carbodiimide based ECM protein immobilization scheme. G) Representative immunofluorescent images of micropatterned hPSCs colonies stained for OCT4, and SOX2. H) Overview of a previously described micro-patterning based hPSC differentiation assay[30] using OCT4 and SOX2 expression levels as indicators of early fate choices to compare PEG and μCP plates. I) Quantified compartments of early fate choices as defined in H), in both PEG, and μCP plates. The media conditions tested were ‘NS’ – Nutristem, Apel (vehicle for the following), ‘BMP’ (BMP4), ‘BA’ (BMP4+ActivinA), ‘FSB’ (bFGF+SB431542) (See Materials and Methods for concentration details). Data represented as mean (+s.d) of four independent replicates. The fate choice responses of hPSCs on both the plates were highly correlated (R^2^ >0.9). J) Representative immunofluorescent images of hPSC colonies stained for OCT4, and SOX2 in the different media conditions tested. Scale bars indicate 500μm.

Given the vital role that interactions between cells and the surrounding ECM play on cellular responses[46], we next asked whether the approach of covalent attachment of ECM molecules interfered with fate decisions of hPSCs micropatterned on the PEG plates. We opted to employ a recently reported two-day assay using OCT4, and SOX2 expression as readouts to assess fate decisions in geometrically confined hPSC colonies[30] (**Fig. 1H**), and directly compared fate acquisition of hPSCs on the PEG plates with μCP plates, a micro-patterning technique that does not require any chemical immobilization of ECM molecules. We observed a highly correlated (R^2^ > 0.9) differentiation response between μCP and PEG plates (**Fig. 1I-J**). Furthermore, the PEG plates responded in a more reproducible manner than the μCP plates both in terms of the number of colonies achieved per well of a 96-well plate, and the number of cells attached per colony (**Sup. Fig. 1G-H**). Taken together, these data demonstrate that the PEG plates enable robust geometric-confinement of cell colonies, and the differentiation response of hPSC colonies micro-patterned using the PEG plates differentiate in a highly correlated manner to those micro-patterned on μCP plates; making them a valuable platform for high-throughput screening studies for the bioengineering community.

### hPSC line screen for peri-gastrulation-like patterning response yields variable responses

Recent studies have reported that BMP4 treatment of geometrically confined hPSC colonies results in self-organized fate patterning of gastrulation-associated markers[24–26]. Notably, these studies demonstrated that the differentiating geometrically-confined hPSC colonies gave rise to a Brachyury (BRA) expressing compartment, representing a primitive-streak-like identity. Given that lineage-specific differentiation potential between hPSC lines is known to vary widely[28–30], we hypothesized that different hPSC lines would induce the primitive-streak-like compartment at different efficiencies. We employed our platfrom to evaluate the response of BMP4-treatment of geometrically-confined hPSC colonies (1mm in diameter) in a screen of the following five hPSC lines: H9-1, H9-2, HES2, MEL1, and HES3-1. The induction medium employed for this screen, and all subsequent experiments (unless otherwise stated) was a Knockout Serum Replacement based medium supplemented with BMP4 and bFGF (see Materials and Methods for composition). Although all hPSC lines tested expressed high levels of pluripotency markers at the start of the differentiation culture (**Fig. S2**), induction of BRA expression levels varied markedly between hPSC lines at 48h after BMP4 treatment (**Fig. 2A,C**). Notably, although the MEL1, and HES3-1 lines were unable to induce the expression of BRA, they did differentiate as indicated by the reduction of SOX2 expression relative to the starting population (**Fig. 2B-C, Fig. S2**). These data indicate that although all hPSC lines tested under these experimental conditions differentiated upon BMP4 treatment, induction of the primitive-streak-like compartment, as indicated by BRA expression, varied considerably.

**Figure 2:**
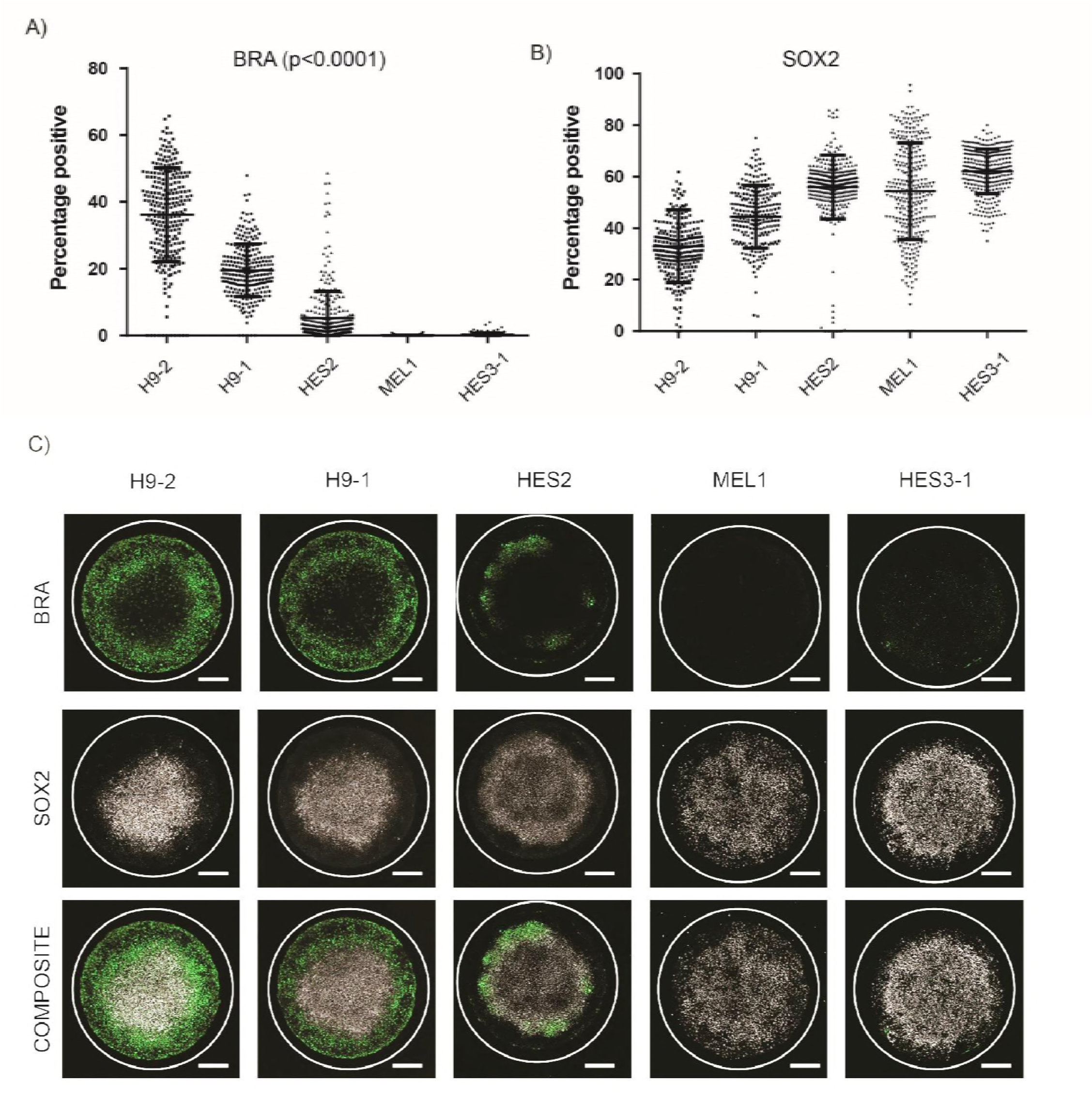
Variability in peri-gastrulation-like induction observed between test hPSC lines. A-B) Quantified expression of BRA (A), and SOX2 (B) observed within the assayed hPSC lines tested. Number of colonies were 252, 245, 327, 288, and 304 for GKH9, 7TGP, HES2, PDX1-GFP, and MIXL1-GFP respectively. Each data point represents individual colonies identified. Data pooled from two experiments. C) Representative immunofluorescent images for BRA, SOX2, and CDX2 for the test hPSC lines. Scale bar represents 200μm.

### hPSC differentiation propensities are correlated with endogenous Nodal signalling

We hypothesized that differences in regulation of key signalling pathways controlling mesendodermal induction between the tested hPSC lines underlay the variation in BRA expression observed in the peri-gastrulation-like patterning. To test this hypothesis, we employed a recently reported approach that addressed a similar question in mouse epiblast stem cell (mEpiSC) lines[47]. In their study, Kojima *et al* made embryoid bodies (EBs) out of various mEpiSC lines, allowed them to spontaneously differentiate in culture conditions unsupportive of pluripotency, and assayed for the expression of differentiation associated genes to compare the transcriptional and functional profiles between the lines[47]. Employing a similar approach, we generated EBs from nine hPSC lines – H9-3, H1, H7, HES3-2 in addition to the previous panel (complete list of lines and their respective culture conditions shown in **Table S1**) – and cultured them in conditions unsupportive of pluripotency for three days and analyzed differentiation marker gene expression levels daily (henceforth – ‘EB assay’) (**Fig. 3A**). We observed strong variation in expression profiles of differentiation associated genes between the test hPSC lines (**Fig. 3Bi**). To simplify data interpretation, we used unsupervised K-means clustering to segregate the hPSC lines into ‘Strong’, ‘Intermediate’, and ‘Weak’ expressers for each gene tested. This analysis revealed distinct sets of responses in the test lines where some lines upregulated expression of genes associated with gastrulation while others upregulated expression of neurulation-associated genes (**Fig. 3Bii**). In a recent study, Funa *et al* showed that Wnt signalling mediated differentiation of hPSCs results in fate acquisition that is dependent on Nodal signalling[48]. Specifically, the authors demonstrated that presence of Nodal signalling during Wnt mediated differentiation of hPSCs resulted in the acquisition of a primitive streak fate, whereas the absence of Nodal signalling during Wnt mediated differentiation resulted in the induction of the neural crest fate[48]. Given that the primitive streak is a gastrulation-associated fate, neural crest arises during neurulation, and the fact that our data demonstrated a gastrulation versus neurulation switch in differentiating hPSC lines, we hypothesized that difference in Nodal signalling could be responsible for these gene expression profiles in our EB assay. Consistent with this hypothesis, the expression of Nodal and GDF3 (a Nodal target) in the differentiating hPSC lines showed a strong trend indicative of their upregulation linked with the induction of gastrulation-associated genes and their absence linked with the induction of neurulation-associated genes (**Fig. 3Ci**). Furthermore, clustering the hPSC lines with reference to the expression profiles of Nodal and GDF3 by either unsupervised K-means clustering (**Fig. 3Cii, S3**), or by hierarchical clustering based on Euclidean distance (**Fig. S4**), indicated that upregulation of Nodal and GDF3 coincided with gastrulation-associated gene expression, whereas their downregulation corresponded with neurulation-associated gene expression.

**Figure 3:**
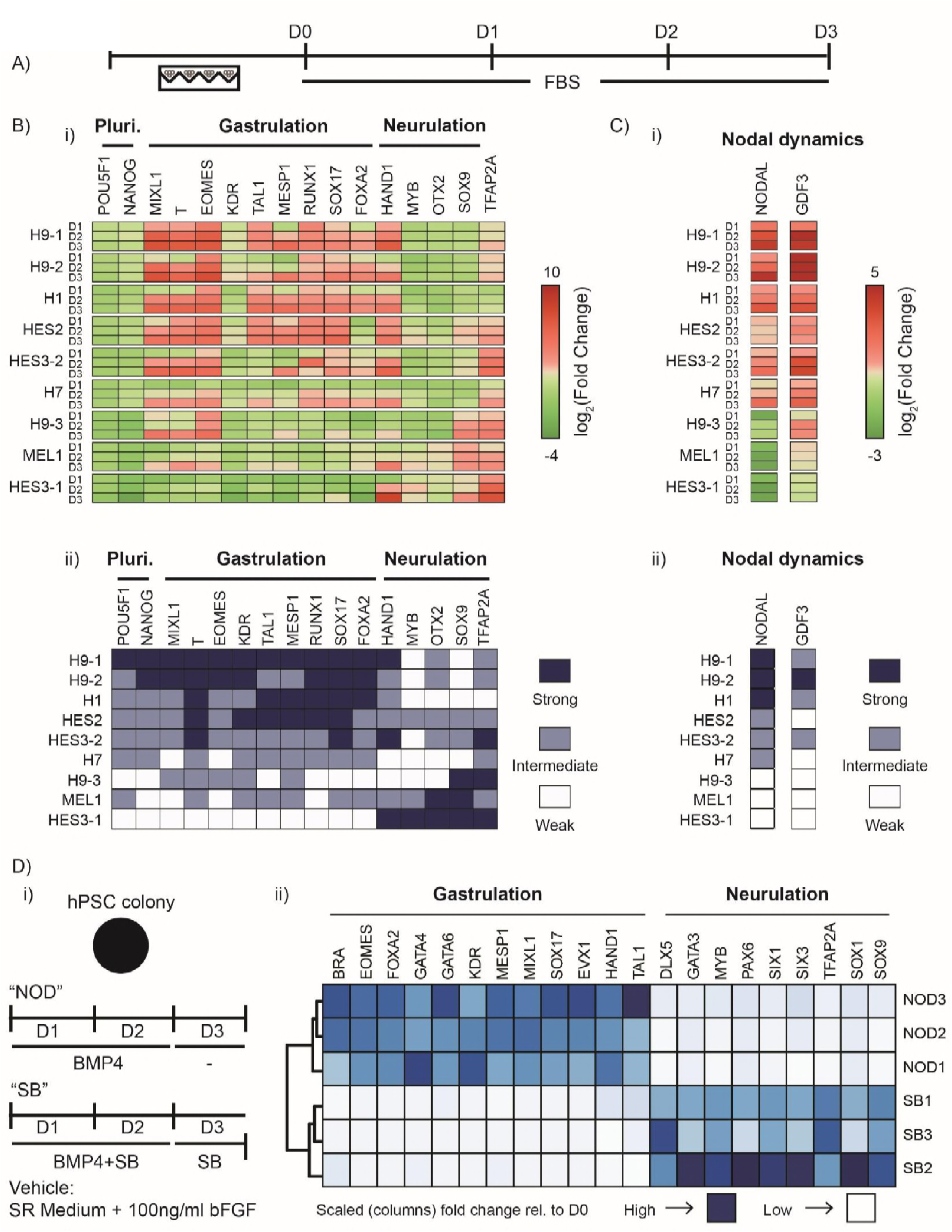
Nodal dissects gastrulation and neurulation associated gene expression profiles. A) Overview of experimental setup for Embryoid Body (EB) assay. EBs were made from each test hPSC line and allowed to spontaneously differentiate in presence of Fetal Bovine Serum (FBS) for three days. B) Observed gene expression dynamics of test cell lines when differentiated as EBs in FBS. i) Observed gene expression for a panel of differentiation associated genes (shown under ‘Gastrulation’ and ‘Neurulation’ groups) along with POU5F1 (OCT4) and NANOG. Data shown as heatmap of mean expression of each day from three biological replicates (s.d. not shown), represented as log_2_(Fold Change) relative to the D0 sample of respective hPSC line. ‘Pluri’ indicates the pluripotency associated genes. ii) Heatmap representation of (B(i)) with the panel of hPSC lines clustered into three groups of ‘Strong’, ‘Intermediate’, and ‘Weak’ responders for each gene using unsupervised K-means clustering. ‘Pluri’ indicates the pluripotency associated genes. C) Nodal dynamics during EB assay. i) Observed gene expression of Nodal and a Nodal signalling target (GDF3). Data shown as heatmap of mean expression of each day from three biological replicates (expression levels for individual replicates shown in Fig. S3B), represented as log_2_(Fold Change) relative to the D0 sample of the respective hPSC line. ii) Heatmap representation of (C(i)) with the panel of hPSC lines clustered into three groups of ‘Strong’, ‘Intermediate’, and ‘Weak’ responders for Nodal, and GDF3 using unsupervised K-Means clustering. D) Effect of modulation of Nodal during previously reported peri-gastrulation-like assay using geometrically-confined colonies of the ‘CA1’ hPSC line[24]. i) Overview of experimental setup. Geometrically-confined colonies of CA1s were induced to differentiate for three days, with either a two-day pulse of BMP4 and Nodal, and just Nodal for the third day; or a two-day pulse of BMP4 and an inhibitor of Nodal signalling (SB431542 – ‘SB’), and just SB for the third day. The vehicle employed in this experiment was SR medium (see Materials and Methods for composition). ii) Heatmap representation of a panel of differentiation genes associated with either gastrulation, or neurulation. Dark blue represents higher levels of expression, whereas light blue represents lower levels of expression. Data shown as mean of three biological replicates. Expression levels of individual replicates shown in **Fig. S7**.

### Validation of gene expression responses in EB assay

We next sought to validate the gene expression differences observed in the differentiating hPSC lines in the EB assay by asking if the variation translated to cell fate acquisition during directed differentiation. Given the key role that MIXL1 plays in the induction of definitive endoderm[49], Kojima *et al* investigated the expression profiles of Mixl1 in their EB assay with mEpiSCs, and demonstrated that Mixl1 expression was predictive of endodermal differentiation bias of the mEpiSC lines[47]. Importantly, much like MIXL1, EOMES is also known to play an important role in the endoderm specification[50,51]. Consistent with this idea, the ‘Strong’, ‘Intermediate’, and ‘Weak’ responders of the panel of hPSCs for both MIXL1, and EOMES contained the identical hPSC line cohorts (**Fig. 3Bii, S5A**), suggesting the likelihood of parallel functions of both these genes in differentiating hPSCs. To validate the gene expression profiles observed in our EB assay, we asked if the observed expression differences of these genes that critically regulate endoderm specification were able to predict the propensity of the hPSC lines to differentiate toward the definitive endodermal fates. Consequently, we differentiated the panel of hPSCs toward definitive endoderm using an established protocol (**Fig. S5B**), and consistent with the findings of Kojima *et al*[47], the expression profiles of endoderm specifiers (MIXL1 and EOMES in our case) in our EB assay closely matched the propensity of the hPSC lines to induce SOX17 expression upon directed differentiation toward the definitive endoderm fate (**Fig. S5C,D**). The differential expression profiles of MIXL1 and EOMES in the EB assay were also able to predict the induction efficiency of mature endodermal fates. Specifically, lines from the MIXL1/EOMES-Strong cluster outperformed candidate lines from the MIXL1/EOMES-Weak cluster in the induction of pancreatic progenitors as marked by the co-expression of PDX1 and NKX6.1 (**Fig. S5E**). These data provide protein level phenotypic validation of the variable gene expression observed in our EB assay.

Given that geometrically confined hPSC colonies are able to induce organized fate patterning[24–26], we asked if subjecting geometrically-confined hPSC colonies to defined endodermal differentiation conditions could be used as an assay to predict the differentiation propensity of hPSC lines. We selected three hPSC lines – H9-1, HES3-2, and HES3-1 – to represent each MIXL1/EOMES induction compartment defined in the EB assay (**Fig. 3Bii, S5A**), and differentiated them as geometrically-confined colonies in defined endodermal induction conditions (**Fig. S6A**). Interestingly, we found that the relative efficiency of SOX17 and FOXA2 double-positive expression under these experimental conditions closely matched the endodermal lineage-bias of the lines as predicted by the EB assay (**Fig. S5A-D, Fig. S6B,C**). Taken together, these data validate the differential gene expression observed between the panel of hPSC lines by demonstrating congruence between MIXL1 and EOMES temporal dynamics and endoderm lineage bias of hPSC lines and provide proof-of-concept data that the defined differentiation protocols in geometrically-confined hPSC colonies can be used as quick assays to assess lineage bias of hPSC lines.

### Nodal dissects gastrulation versus neurulation associated hPSC differentiation

Thus far, our data showed that in conditions that do not support pluripotency, differentiating embryoid bodies made from hPSC lines assume a transcriptional state associated with either gastrulation or neurulation, and endogenous Nodal dynamics correlated with this switch. However, whether the differential Nodal dynamics caused the switch in the acquired transcriptional state or if the association was merely correlative remained unclear. Given that BMP4 treatment has been previously reported to induce gastrulation-associated fate patterning[24–26], and that we have previously demonstrated the robust response of peri-gastrulation-like fate acquisition in the CA1 hPSC line[24], we revisited the peri-gastrulation-like model in hPSC colonies to test if Nodal signalling had a direct effect in regulating this switch. We asked if inducing geometrically confined colonies of the CA1 line to differentiate in response to BMP4 either in the presence or absence of a small molecule inhibitor of Alk4/5/7 receptors (SB431542, hereafter ‘SB’) which antagonizes Nodal signalling (**Fig. 3Di**) recapitulated the observed switch in emergent gene expression. After a three-day induction, we observed that colonies grown in the presence of SB upregulated genes associated with neurulation whereas those grown in the absence of SB upregulated genes associated with gastrulation (**Fig. 3Dii, Fig. S7**). These results are consistent with our hypothesis that Nodal signalling distinguishes gastrulation and neurulation-associated gene expression profiles in differentiating hPSCs. Given that differentiating hPSCs in the absence of Nodal signalling upregulated neurulation associated genes, we next set to investigate if BMP4 treatment of geometrically confined hPSC colonies in presence of SB gave rise to early neurulation-associated spatially patterned fate allocation.

### An RD network in BMP signalling can self-organize pSMAD1 activity independent of Nodal

In a recent study, we demonstrated that the peri-gastrulation-like fate patterning in geometrically confined hPSC colonies occurs via a stepwise process of RD and PI where a BMP4-Noggin RD network self-organizes a phosphorylated SMAD1 (pSMAD1) signalling gradient within the colonies, resulting in the peri-gastrulation-like fates being patterned in a manner consistent with the PI paradigm[24]. We set out to investigate if a conserved mechanism would give rise to neurulation-associated fate patterning. As a first step, we asked if a BMP4-Noggin RD network governed pSMAD1 self-organization within the geometrically-confined hPSC colonies treated with BMP4 and SB. Consistent with the presence of a BMP4-Noggin RD network[24], we observed an upregulation of both BMP4 and Noggin upon BMP4 treatment of hPSCs in the presence of SB (**Fig. 4A**). We next asked if BMP4 treatment of geometrically-confined hPSC colonies in the presence of SB would result in the self-organized gradient of nuclear localized pSMAD1. Indeed, pSMAD1 activity within the colonies rapidly self-organized into a radial gradient under these experimental conditions (**Fig. 4B-D**). We next queried the importance of Noggin in the formation of the pSMAD1 gradient by generating two homozygous knock-outs of Noggin (‘C1’, and ‘C7’) using Crispr/CAS9 (characterization of lines shown in **Fig. S8**) and tested whether the absence of Noggin compromised the self-organization of pSMAD1. Consistent with our hypothesis that a BMP4-Noggin RD network was underlying the pSMAD1 self-organization, the formation of the pSMAD1 signalling gradient was significantly compromised in Noggin knockout lines C1 and C7 compared to the wildtype control (**Fig. 4E-F**), indicating an integral involvement of BMP inhibitors like Noggin in the self-organization of the pSMAD1 signalling gradient. In our previous study using the peri-gastrulation-like model, we showed that a BMP4-Noggin RD computational model predicts the experimentally observed responses of a pSMAD1 self-organized gradient at the periphery and the center of the colonies to perturbations to the BMP4 dose in the induction medium and size of the geometrically-confined hPSC colony[24]. Specifically, we showed that reducing the BMP4 dose while maintaining the colony size reduces the levels of pSMAD1 at the periphery, and reducing the colony size while maintaining a constant BMP4 dose in the induction medium results in an increase of pSMAD1 levels at the center of the colonies[24]. We reasoned that a conserved mechanism underlying the pSMAD1 self-organization would result in identical responses to these perturbations. Consistent with our anticipated results, reducing the BMP4 dose in the induction medium while maintaining the colony size resulted in a reduction of the detected immunofluorescent levels of nuclear localized pSMAD1 at the colony periphery (**Fig. S9A-C**). Furthermore, reducing the colony size while maintaining the BMP4 dose in the induction medium increased the detected immunofluorescent levels of nuclear localization of pSMAD1 at the colony centers (**Fig. S9D-E**). Taken together, these data demonstrate that in absence of Nodal signalling, pSMAD1 activity in the geometrically-confined hPSC colonies self-organizes into a signalling gradient and suggest that a BMP4-Noggin RD system governs this observation (**Fig. 4G**) – consistent with our previous study[24].

**Figure 4:**
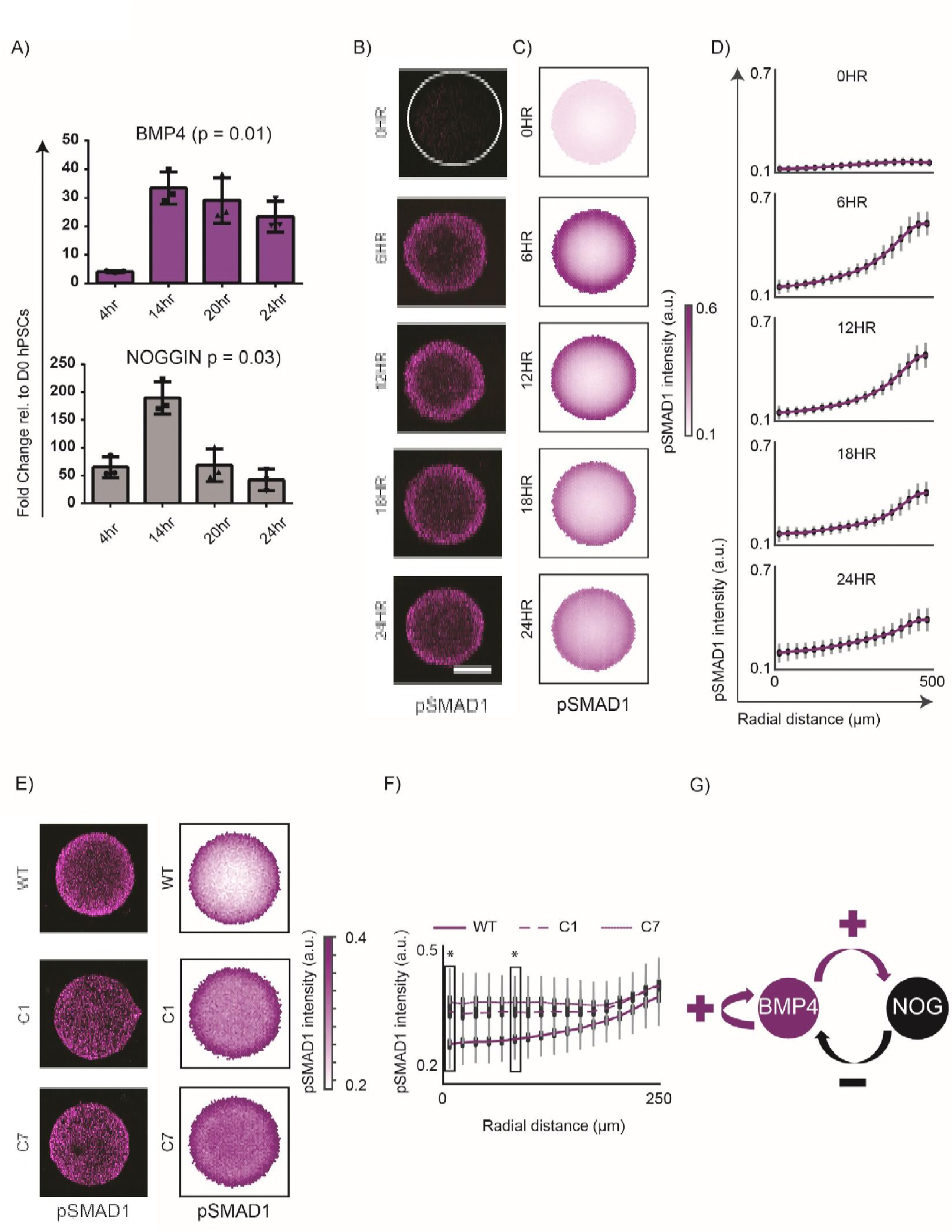
Interaction network between BMP4-Noggin underlies self-organization of pSMAD1 gradient. A) Temporal gene expression for BMP4 and Noggin at 4h, 14h, 20h, and 24h after BMP4 treatment. Data shown as mean ± s.d. of three independent experiments. The p-values shown were calculated using Kruskal-Wallis test. B) Representative immunofluorescent images of geometrically confined hPSC colonies of 500μm in diameter stained for pSMAD1 after different times (0h, 6h, 12h, 18h, and 24h) of BMP4 exposure. Scale bar represents 200μm. C) Average pSMAD1 intensity represented as overlays of 231, 241, 222, 238, and 228 colonies for respective induction times. Data pooled from two experiments. D) The average radial trends of pSMAD1 at each duration shown as line plots. Standard deviations shown in grey, and 95% confidence intervals shown in black. E-G) Response of pSMAD1 self-organization in homozygous knockout lines of Noggin. E) Representative immunofluorescent images of geometrically confined hPSC colonies of wild-type (WT), Noggin^-/-^ clones C1, and C7 (characterization shown in Fig. S8) stained for pSMAD1 after 24h of BMP4 exposure. F) The average radial trends of pSMAD1 shown for the WT, C1, and C7 clones. Data pooled from two experiments and include 151, 150, 150 colonies for each line respectively. G) The average radial trends of pSMAD1 at each duration shown as line plots. Standard deviations shown in grey, and 95% confidence intervals shown in black. The p-values were calculated using Mann-Whitney U-test. ^∗^ indicates p<0.0001 for each clone relative to the WT control. H) Model of reaction-diffusion mediated self-organization of pSMAD1.

### Nodal signalling contributes to the shape of the self-organized pSMAD1 gradient

Our data indicate that the pSMAD1 signalling gradient self-organizes via an RD network present in the BMP signalling pathway where Noggin functions as an important inhibitor (**Fig. 4**). Given that Nodal signalling targets include multiple BMP antagonists such as Cer1, GDF3, Follistatin (FST), etc.[52], we asked if Nodal signalling contributed to the formation of the pSMAD1 signalling gradient in BMP4-treated geometrically confined hPSC colonies. To probe the role of Nodal in the observed pSMAD1 self-organization, we compared the formation of the pSMAD1 gradient in geometrically confined hPSC colonies of 500μm diameter treated with BMP4 for 24h where the induction media either contained Nodal or SB (**Fig. 5A**). The pSMAD1 signalling gradients formed in the presence and absence of Nodal signalling were significantly different from each other when treated with either 25ng/ml of 50ng/ml of BMP4 in the induction medium demonstrating the involvement of Nodal signalling in the formation of the self-organized signalling gradient (**Fig. 5B-D, S10A-C**). The results from these studies provided a few notable observations. First, in 500μm diameter colonies treated for 24h with BMP4 in presence of Nodal ligands, we observed two prominent peaks of pSMAD1 expression – one peak was observed at the periphery as expected from previous reports[24–26], and another on at the colony center that has not been previously reported in colonies of this size (**Fig. 5B-D**). This observation provides further support to the proposition that the self-organization of pSMAD1 arises via an RD mechanism which can result in spatial oscillations of morphogen activity[24], while providing evidence of crosstalk between BMP and Nodal signalling pathways in establishing the pSMAD1 signalling gradient. Another notable observation was that the pSMAD1 expression that declined from the periphery of the colonies dropped rapidly in the presence of Nodal ligands, but the decline was far more gradual in the presence of SB (**Fig. S10A-C**). Cells expressing discernible levels of pSMAD1 were observed much farther into the colony from the periphery than observed in the condition when Nodal ligands were present in the induction medium. Finally, the level of pSMAD1-associated immunofluorescence detected at the colony periphery in the presence of SB was significantly higher than the levels detected in the presence of Nodal (**Fig. 5B-D, S10A-C**). Taken together these data provides further justification for the hypothesis that the pSMAD1 signalling gradient arises via an RD mechanism, and demonstrate that Nodal signalling contributes to the shape of the self-organized pSMAD1 signalling gradient.

**Figure 5:**
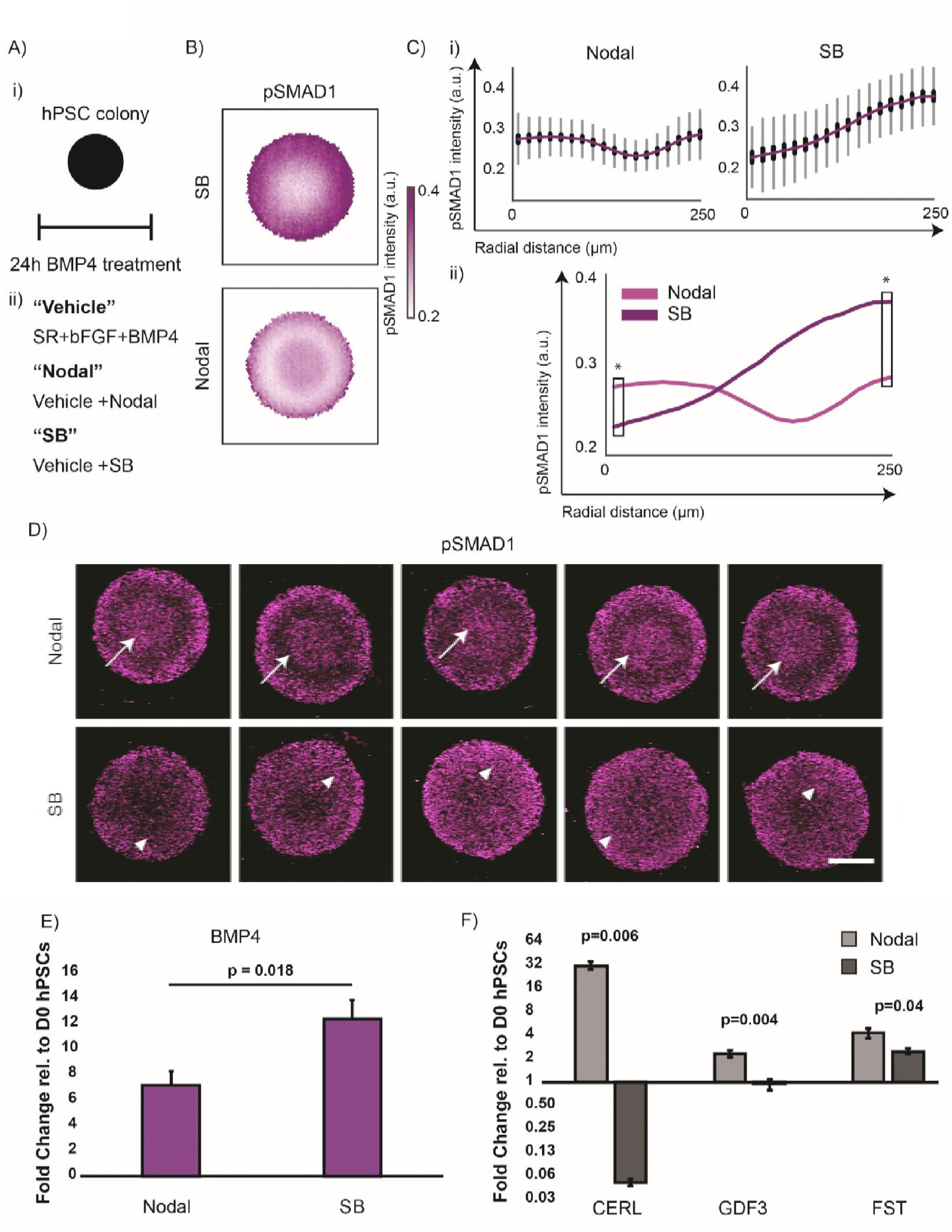
Nodal signalling contributes to the formation of the pSMAD1 gradient. A) (i-ii) Overview of experimental setup. (i) Geometrically confined hPSC colonies were treated with BMP4 for 24h. (ii) Media tested. ‘**Vehicle**’ indicated SR medium (see Materials and Methods for composition) supplemented with BMP4 and bFGF. ‘**Nodal**’ and ‘**SB**’ indicated vehicle supplemented with either Nodal (100ng/ml) or μM SB431542. B-D) Perturbing Nodal signalling results in a significant change in the pSMAD1 self-organized gradient formation. B) Average pSMAD1 intensity represented as overlays of 188, and 163 colonies for Nodal and SB conditions respectively. Data pooled from two experiments. C) The average radial trends of pSMAD1 shown as line plots. (i) Line plots shown individually for SB and NODAL conditions. Standard deviations shown in grey, and 95% confidence intervals shown in black. (ii) Line plots represented in the same graph. The p-values were calculated using Mann-Whitney U-test. ^∗^ indicates p<0.0001. D) Representative immunofluorescence images of 500μm diameter hPSC colonies stained for pSMAD1 after 24h of BMP4 treatment in ‘Vehicle’ and ‘SB’ conditions (average response shown in B). Scalebar represents 200μm. White arrows indicate regions where second peak of pSMAD1 appears. White triangles indicate regions of discernable pSMAD1 levels that appear to be lower than the levels at the colony periphery. E) Gene expression for BMP4 after 24h of treatment with either ‘Vehicle’ of ‘SB’ media (described in A(ii)) of hPSCs. Data shown as mean + s.d. (n=3, technical replicates, independent wells). The p-value was calculated using two-sided t-test. F) Gene expression of Activin-Nodal pathway associated targets that are known antagonists of BMP signalling (CERL, GDF3, Follistatin – ‘FST’). The data represented as mean ± s.d. of hPSCs from three technical replicates, independent wells. The experiment was performed once. The p-values were calculated using two-sided Student’s t-test.

Since an RD network results in the morphogen gradient as a consequence of the expression of both activators and inhibitors of the morphogen[11,12], we hypothesized that the likely reasons for this observation could be due to either a change in the amount of activator (change in BMP4 levels) or the amount of inhibitor (change in the level of BMP antagonists) in the system. When we tested gene expression of activators and inhibitors after 24h of Vehicle versus Nodal or SB treatment on hPSCs which either allowed Nodal expression or dramatically downregulated it (**Fig. S10D**), SB treatment provoked an increased positive feedback as indicated by increased detected levels of BMP4 transcripts (**Fig. 5E**); and a reduced negative feedback as indicated by significantly reduced transcript levels of BMP antagonists like CERL, GDF3, and FST (**Fig. 5F**). Taken together, these data suggest that Nodal signalling can contribute to the RD-mediated self-organization of the pSMAD1 signalling gradient; and that this contribution might occur due to a change in the levels of activators and antagonists of BMP signalling. Having established that pSMAD1 activity in the geometrically confined hPSC colonies treated with BMP4 and SB self-organizes into a signalling gradient, we next focused on investigating if this gradient induced the expression of fates associated with the differentiating ectoderm.

### Pre-neurulation-like fate patterning arises in a manner consistent with PI

In the presence of Nodal signalling, BMP4 treatment of geometrically-confined hPSC colonies results in self-organized pSMAD1 gradient and a spatially patterned acquisition of gastrulation-associated fates[24]. Although we observe the formation of the pSMAD1 gradient when geometrically-confined hPSC colonies are treated with BMP4 and SB (**Fig. 4-5**), we did not observe expression of key gastrulation associated markers like BRA, EOMES, SOX17, and GATA6 (**Fig. S11A-B**). These observations are consistent with the need of Nodal in inducing gastrulation-associated fates[24]. Since we observed that differentiating hPSCs in the absence of Nodal signalling upregulate a neurulation-associated gene profile (**Fig. 3D**), we asked if BMP4 and SB treatment of geometrically confined hPSC colonies resulted in the fate patterning associated with the differentiating ectoderm. After the germ layers segregate from the epiblast, a BMP signalling gradient along the medial-lateral axis in the developing ectoderm patterns the early pre-neural (PN) tissue at the medial end, and non-neural (NN) tissue at the lateral end appropriately arranging the tissue for the onset of neurulation[8]. The PN tissue gives rise to the neural plate (NP), which later folds to form the neural tube[5,7], and the NN tissue gives rise to the non-neural ectoderm (NNE) and the neural plate border (NPB)[8]. The NNE subsequently specifies to generate the epidermis and the NPB is a multipotent tissue that produces the neural crest (NC) and the craniofacial placodes in the anterior ectoderm[8,9,53–55]. The early PN region maintains the expression of SOX2 which is present in the epiblast, and the early NN regions induce expression of markers like GATA3[8]. Furthermore, markers like transcription factor AP2-alpha (TFAP2A) mark the NN, NNE, and maturing NPB region that marks the NC fate; and SIX1 are expressed in the maturing NPB region which marks panplacodal competent tissues[8]. Consistent with our observation that BMP4 treatment in the absence of Nodal signalling upregulated genes associated with neurulation, we observed spatially segregated expression of SOX2 (PN), and GATA3 (NN) with concomitant expression of TFAP2A (NN, NPB), and SIX1 (panplacodal competent NPB) (**Fig. S11C**). We define this fate patterning as ‘pre-neurulation-like’ and using SOX2, and GATA3 as the markers of the PN, and NN tissues, we set out of test if the fate patterning arose in a manner consistent with the positional information (PI) paradigm.

Given that the PI paradigm posits that developmental fates arise due to thresholds of morphogen levels, and we asked if the perturbations of pSMAD1 levels at the colony periphery (**Fig. S9A-C**) and the colony center (**Fig. S9D-E**) resulted in pSMAD1 threshold mediated changes in expression of GATA3 (NN), and SOX2 (PN) fates respectively. Consistent with the idea of a pSMAD1 threshold dependent patterning of the PN and the NN tissues marked by GATA3, and SOX2, we find that reducing the pSMAD1 levels at the colony periphery (**Fig. S12A**) significantly reduced the GATA3 expression at the colony periphery (**Fig. S12B-C**) and increasing the pSMAD1 levels at the colony center (**Fig. S12D**) dramatically reduced the SOX2 expression (**Fig. S12E-F**). These data indicate that thresholds of pSMAD1 regulated the patterning of the SOX2 and GATA3 within the geometrically confined hPSC colonies. However, the formalization of the PI paradigm has been updated to include time as a critical parameter that patterns the developmental cell fates. Specifically, fate patterning mediated by PI is known to arise as a function of the morphogen concentration and time of induction[17,18,24,56]. Consequently, we tested four different doses of BMP4 (3.125ng/ml, 6.25ng/ml, 12.5ng/ml, and 25ng/ml) in the induction medium for four different induction times (12h, 24h, 36h, and 48h) and measured the levels of SOX2 and GATA3 detected. We observed that the fate patterning of GATA3 arose as a function of both the concentration of BMP4 in the induction medium and the time of induction (**Fig. 6A-B, C(i), Fig. S13**) indicating that the patterning within the geometrically confined colonies arises in a manner consistent with PI (**Fig. 6C(ii)**).

**Figure 6:**
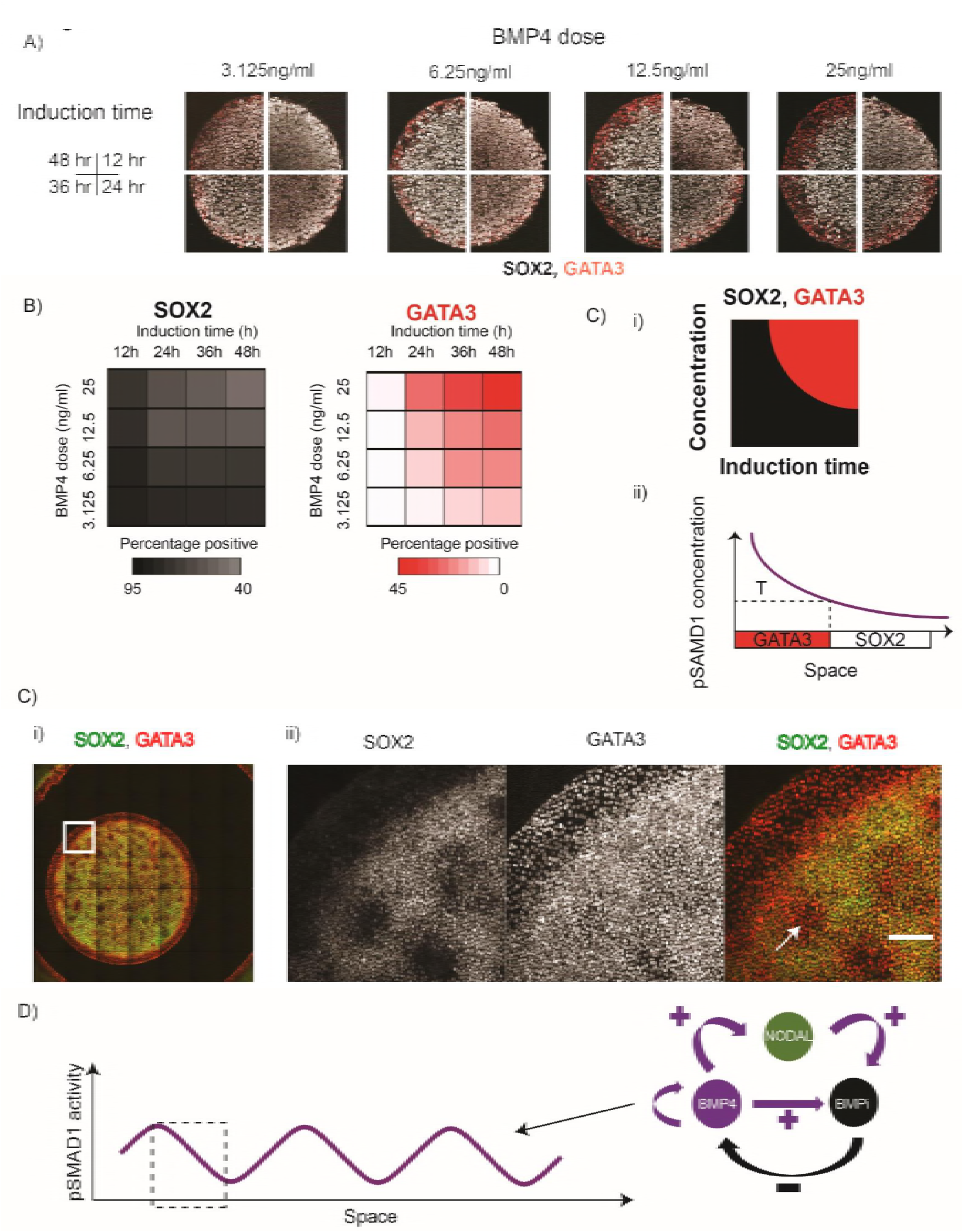
Pre-neurulation-like fates arise in a manner consistent with positional-information. A) Representative immunofluorescence images of 500μm diameter colonies stained for SOX2, and GATA3 after different doses (6.25ng/ml, 12.5ng/ml, 25ng/ml, and 50ng/ml) and times of BMP4 treatment. Scale bar represents 200μm. B) Mean expression levels of SOX2 and GATA 3 represented as heat maps. Darker shades represent higher expression levels and lighter shades represent lower levels of expression (for detailed data see Fig. S12). C) (i-ii) Model of GATA3 patterning. (i) Overview of (B) where GATA3 is expressed as a function of BMP4 dose and induction time. (ii) Fate patterning of GATA3 consistent with positional information. T indicates the presumptive threshold of fate switch to GATA3. D) Treatment of geometrically confined-hPSC colonies of 3mm diameter with 200ng/ml of BMP4 and SB for 48h results in multiple peaks of GATA3 expressing regions consistent with RD hypothesis. i) Representative stitched images of 3mm diameter hPSC colonies differentiated with 200ng/ml of BMP4 for 48h. Scale bar represents 1mm. ii) Zoomed section outlined by the white square in (i). White arrows indicate regions of high GATA3 and low SOX2 expression indicative of PI mediated fate patterning due to presumptive localized pSMAD1 expression. The experiment was repeated three times. Additional images shown in **Fig. S22**.

### A stepwise model of RD and PI governs pre-neurulation-like fate patterning

Thus far, our data indicate that the pSMAD1 gradient was enforced outside-in within the geometrically confined hPSC colonies via a BMP4-Noggin RD network, and the pre-neurulation-like fates arose in a manner consistent with PI. In agreement with this idea, perturbing the shapes of the geometrically confined hPSC colonies did not result in fate patterning that deviated from the expected results (**Fig. S14**). However, a strong test of this overall model is asking if large colonies are able to generate stereotypical RD-like periodic signalling and fate profile. We previously reported that treatment of large geometrically confined hPSC colonies (3mm) with high doses of BMP4 would give rise to multiple foci of BMP activity as indicated by pSMAD1 staining, and patterned gastrulation-associated fates[24] – consistent with the expected spatial oscillations in accordance with the RD paradigm. Surprisingly, when we tested the response of pSMAD1 spatial signalling dynamics in 3mm diameter colonies after BMP4 and SB treatment for 24h, we did not observe any obvious additional foci at either 50ng/ml (**Fig. S15**) or 200ng/ml (**Fig. S16**) BMP4 dose. Of note, the medium used for differentiating these geometrically confined hPSC colonies contained Knockout Serum Replacement (SR). An ingredient of SR called AlbumaxII is known to contain lipid associated proteins that have been shown to have an effect on hPSC biology – mechanisms of which are currently unclear[57,58]. We asked if using medium devoid of SR would rescue the expected appearance of multiple foci of pSMAD1 activity and GATA3 expression consistent with the predictions of the RD paradigm[24]. Indeed, when we tested N2B27 medium which does not contain any AlbumaxII or SR (see Materials and Methods for composition), a 24h BMP4 and SB treatment of hPSC colonies of 3mm diameter resulted in rudimentary peaks of PSMAD1 activity at a BMP4 dose of 50ng/ml (**Fig. S17**) and prominent peaks of pSMAD1 activity at a dose of 200ng/ml (**Figs. S18-S19**). In addition, after 48h of BMP4 and SB treatment, although we did not note any additional foci of GATA3 in SR medium at BMP4 doses of either 50ng/ml (data not shown) or 200ng/ml (**Fig. S20**), in an N2B27 basal medium supplemented with SB, rudimentary peaks were observed at 50ng/ml of BMP4 (**Fig. S21**), and robust peaks were noted at 200ng/ml of BMP4 (**Figs. 6D, S22**). These observations are consistent with the proposition that an RD network self-organizes BMP signalling activity (**Fig. 6E**) and the patterned fates arise in a manner consistent with PI (**Fig. 6C**), although we note that undefined components present in the induction medium contribute to deviations from the expected results.

### PN and NN regions give rise to definitive ectodermal fates

As a final validation that BMP4 and SB treatment induced pre-neurulation-associated fates in the differentiating geometrically confined hPSC colonies, we asked if the patterned fates of the early PN and NN tissues were capable of inducing marker expression of definitive ectodermal fates like the NP, the NC, and the NNE. During embryogenesis, the NNE specifies toward the lateral end of the medial-lateral axis due to sustained levels of high BMP signalling in the ectoderm; the NC fate is specified at regions of intermediate BMP levels that activate wnt signalling; and the NP is specified at the medial end where the tissue is subject to low/no BMP signalling. To test the competence of the pre-neurulation-like patterned colonies to give rise to these fates, we treated the colonies with BMP4 and SB for 24h, then tested three different treatments. Specifically, we either treated the colonies for a further 48h with BMP and SB and stained for keratins using a pan-keratin antibody and DLX5 (markers of NNE); or CHIR99021 (‘CHIR’ – a wnt agonist) and SB and stained for SOX10 (a marker of the NC fate); or for a period of 72h with Noggin and SB and stained for PAX6 (**Fig. 7A**). Consistent with our expected results, we observed that sustained BMP4 and SB treatment resulted in robust expression of DLX5 and showed clear staining of a pan-keratin antibody, indicating acquisition of an NNE identity (**Fig. 7B**). Furthermore, robust SOX10 staining was observed in colonies treated with CHIR and SB (**Fig. 7C**); and the colonies treated with Noggin and SB expressed PAX6 – a bona fide marker of the NP (**Fig. 7D**).

**Figure 7:**
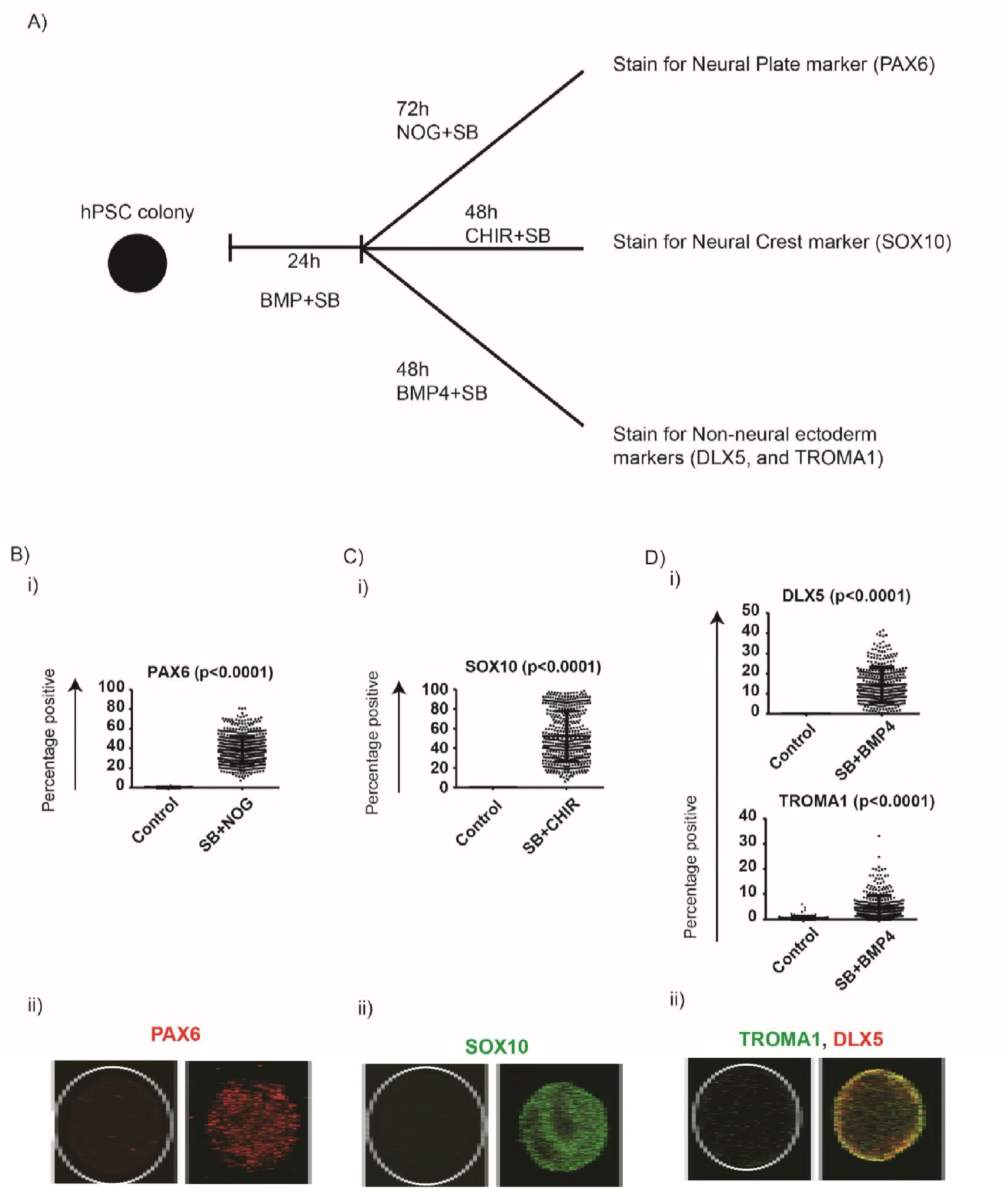
Pre-neurulation-like platform can give rise to definitive fates associated with the differentiating ectoderm. AA) Overview of the experimental setup. Geometrically confined hPSC colonies were treated with BMP4 for 24h, and then treated with one of the following conditions: SB and Noggin for 72h and subsequently stained for PAX6; SB and CHIR99021 (CHIR) for 48h and stained for SOX10; SB and BMP4 for 48h and stained for DLX5 and TROMA1. B) Expression of NP marker (PAX6) in colonies differentiated with BMP4 for 24h and SB+Noggin for 72h. (i) Quantified expression observed for PAX6 observed in the treated and control conditions. The number of colonies were 76 for control, and 606 for treated. (ii) Immunofluorescent images of representative colonies stained for PAX6. C) Expression of NC marker (SOX10) in colonies differentiated with BMP4 for 24h and SB+CHIR for 48h. (i) Quantified expression observed for SOX10 observed in the treated and control conditions. The number of colonies were 286 for control, and 493 for treated. (ii) Immunofluorescent images of representative colonies stained for SOX10. D) Expression of NNE markers (DLX5 and TROMA1) in colonies differentiated with BMP4 for 24h and SB+BMP4 for 48h. (i) Quantified expression observed for DLX5 and TROMA1 observed in the treated and control conditions. The number of colonies were 163 for control, and 376 for treated. (ii) Immunofluorescent images of representative colonies stained for DLX5 and TROMA1. For B(i), C(i), and D(i), each data point represents an identified colony, and bars represent mean ± s.d. The data were pooled from two experiments, and the p-values were measured using Mann-Whitney U test.

Taken together, our data are consistent with our hypothesis that a RD network in BMP signalling self-organizes the pSMAD1 gradient in geometrically confined hPSC colonies and Nodal signalling dissects peri-gastrulation-associated and pre-neurulation-associated fates (**Fig. 8**) that arise within these colonies in a manner consistent with PI.

**Figure 8:**
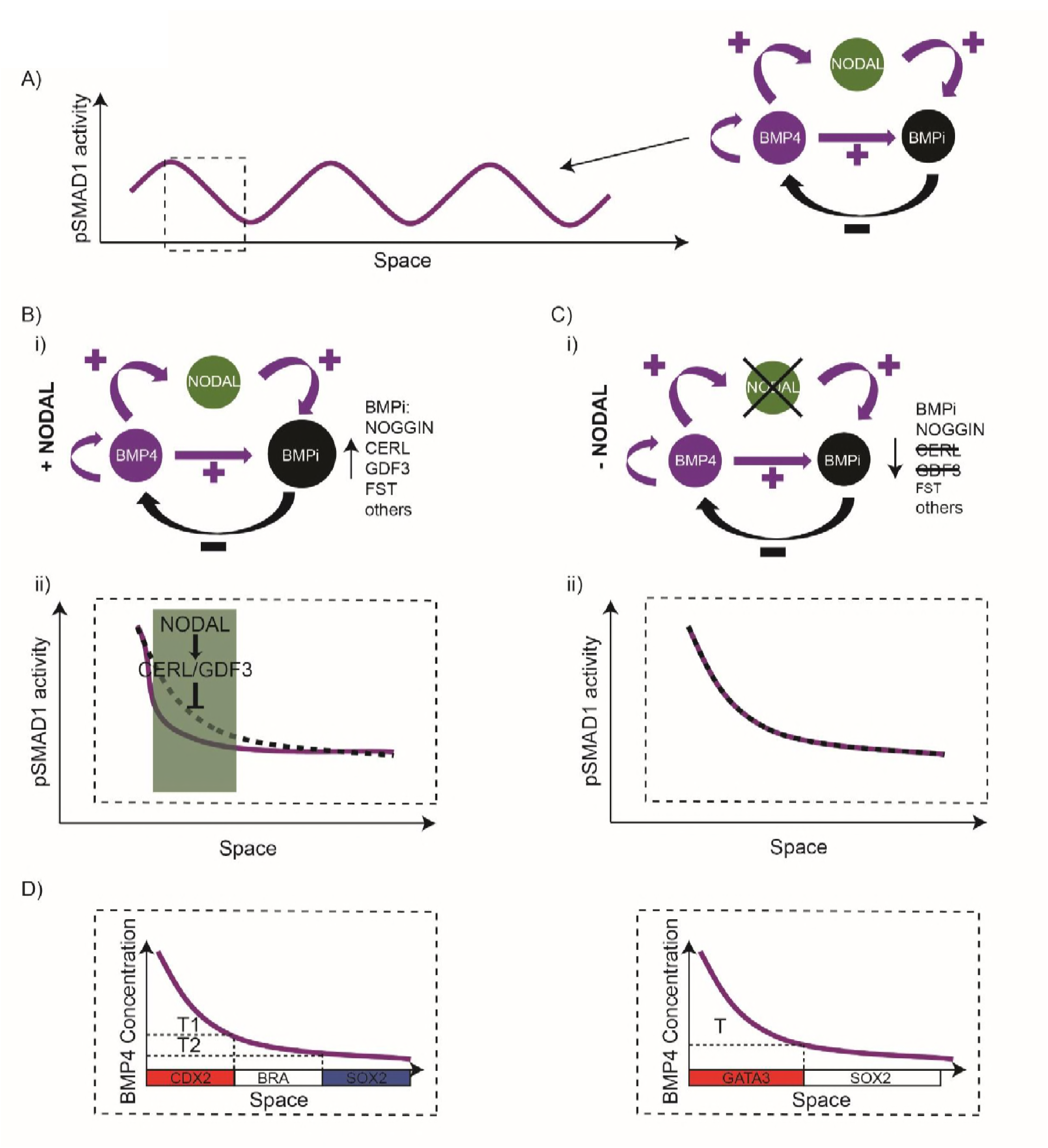
Mechanism of Nodal dependent fate patterning in the geometrically confined hPSC colonies. A) Model overview for self-organization of pSMAD1: An RD network in BMP signalling which comprises BMP ligands, NODAL, and BMP antagonists self-organizes the pSMAD1 gradient within the geometrically confined hPSC colonies. B) (i-ii) pSMAD1 self-organization in the presence of Nodal. (i) BMP antagonists downstream of Nodal signalling (like CERL, GDF3, FST) can contribute to the self-organization of the pSMAD1 gradient. (ii) The presence of BMP antagonists in the Nodal pathway downregulates pSMAD1 levels enforcing a sharp gradient from the colony periphery. Green region signifies region of Nodal mediated downregulation of BMP signalling. Dotted black line represents gradient that would arise in the absence of BMP antagonists in the Nodal pathway. Purple line represents gradient established due to pronounced inhibition of BMP signalling. C) (i-ii) pSMAD1 self-organization in the absence of Nodal. (i) In the absence of Nodal signalling the overall level of BMP inhibitors is reduced due to removal of CERL, GDF3, and reduction in FST levels. (ii) The established gradient is more gradual relative to when Nodal signalling is active. Dotted black line represents gradient that would arise in the absence of BMP antagonists in the Nodal pathway. Purple line represents gradient established due to pronounced inhibition of BMP signalling. D) In the presence of Nodal signalling, the fate patterning recapitulates the peri-gastrulation-like stage of human development. In the absence of Nodal signalling, the fate patterning recapitulates pre-neurulation-like stage of human development. In both instances, the fate patterning arises in a manner consistent with positional information.

## Discussion

### Screening platform for organoid-like structures and hPSC ‘finger-printing’ assay

Stem cells have a remarkable ability to self-organize into complex, higher-order tissues. Numerous studies have exploited this capacity of stem cells to generate structures that resemble organs and developmentally-relevant tissues[23,36,59–64]. These so-called ‘organoids’[22,65] offer exciting possibilities as they can be employed as an experimental model for screening studies of organs/tissues of interest because they are derived from human cells while maintaining aspects of the structure and organization of the native tissues. Although the field has taken impressive strides towards making organoids for a variety of different organs, achieving a reproducible response between each organoid remains problematic. Furthermore, quantitative image analysis of immunofluorescent data from high-content organoid-based screening studies is currently challenging. An alternative approach for harnessing the potential of employing appropriately organized tissues in screening studies is to start with 2-dimensional cultures of the specific stem/progenitor cells with controlled geometries and allow them to self-organize into ‘organoid-like’ tissue surrogates. Of late, numerous studies have employed this approach to derive developmentally-relevant tissue organization [24–26,66,67]. Importantly, the response between individual organoid-like structures – for specific cell lines and medium conditions – is far more reproducible than what can currently be achieved in 3D organoids. Furthermore, given that these organoid-like structures are secured in position for the assay-duration, they are far more amenable to high-content image analysis than their 3D counterparts. The high-throughput platform we report enables robust geometric confinement of a variety of cell types and can be employed for numerous applications. Indeed, we have employed it to micro-pattern human PSCs, mouse PSCs, Retinal Pigmented Epithelial cells, human hemogenic, and pancreatic progenitors, human cardiomyocytes, human keratinocytes, mouse embryonic fibroblasts (MEFs), among others. This platform is poised to be employed for high-throughput drug screens of organoid-like surrogates of a variety of tissues.

As proof-of-principle for one such application, we employed the peri-gastrulation-like patterning to assess the differentiation propensity of hPSC lines. To capitalize on the well-established promise of hPSCs[68], many groups and initiatives have banked large numbers of human induced (hi)PSCs for future use in regenerative medicine applications[28]. Importantly, it is widely recognized that different hiPSC lines – even if derived from identical genetic and tissue backgrounds – significantly vary in their ability to induce certain cell fates[28–30]. Consequently, the field needs an assay that enables rapid, high-content quantification of lineage bias of a starting pool of hPSCs from which an ideal line would be chosen to produce cells of a target fate. Although some assays currently attempt to provide a solution to this need[31–33], they are either qualitative, or prohibitively expensive and time consuming. In addition to the variation in BRA expression observed in the peri-gastrulation-like assay, we demonstrate that Wnt3a and ActivinA treatment of geometrically confined hPSC colonies of different hPSC lines results in variable endoderm induction efficiencies that mirror the predicted propensity both from directed differentiation toward definitive endoderm, and as indicated by the temporal dynamics of MIXL1 and EOMES in the EB assay. Notably, this observation parallels the observation reported by Kojima et al who showed that the temporal dynamics of MIXL1 during undirected differentiation of mEpiSC lines predicted endodermal differentiation propensity[47]. Consequently, we both corroborate the approach taken by Kojima *et al* in the human system and provide proof-of-concept data that indicates that morphogen treatment of geometrically confined hPSC colonies in defined conditions might represent a rapid, quantitative, and an inexpensive solution for finger-printing hPSCs.

### Involvement of Nodal in RD and PI associated with the BMP signalling

Although both RD and PI have been studied in a variety of different model systems and have provided much insight into how developmental fate patterning occurs during embryogenesis, the question of how multiple different signalling pathways can work in concert to execute the rules associated with each paradigm remains unclear. In this study we demonstrate that Nodal signalling works in concert with BMP signalling to not only orchestrate the self-organization of pSMAD1 activity into a signalling gradient in a manner consistent with RD, but also coordinates the interpretation of the gradient into either peri-gastrulation-like, or pre-neurulation-like fate patterns in accordance with PI.

#### Reaction diffusion mechanisms

In a recent study, we proposed that the pSMAD1 signalling gradient self-organizes under the regulation of a BMP4-Noggin RD system[24]. Our data in this study demonstrates that the pSMAD1 gradient formed in the presence and absence of Nodal signalling is significantly different (**Fig. 5, S10**); indicating the necessity of updating that proposed RD network topology to include Nodal signalling which is activated downstream of BMP signalling during the onset of mammalian gastrulation. Our data also suggest that the role played by Nodal signalling may, at least in part, be enacted by BMP antagonists that are targets of Nodal signalling, which is consistent with the fact that RD-mediated self-organization of morphogen signalling relies on the function of the morphogen inhibitors[11,12]. For instance, Nodal signalling gradient during zebrafish embrogenesis forms due to a Nodal-Lefty RD system[69], and Rodgers *et al* have demonstrated that removing the Lefty in an increased amount of Nodal signalling and specification of mesendoderm in the embryos during gastrulation[70]. Under experimental conditions where Nodal signalling is inhibited due to SB supplementation, Noggin, Follistatin (**Fig. 5F**), Gremlin family proteins[26], and possibly others can act as the inhibitors that enforce the pSMAD1 signalling gradient; and in conditions permissive of Nodal signalling, CERL and GDF3 and possibly others can further antagonize BMP signalling. In support of this notion, a previous study has reported that siRNA mediated inhibition of CERL and Lefty in experimental conditions permissive of Nodal signalling dramatically compromises the formation of peri-gastrulation-associated patterns (Fig. 6 in ref. *25*). Taken together, the topology of the RD network in BMP signalling needs to incorporate the role played by Nodal and potentially multiple BMP4 antagonists in addition to Noggin (like CERL GDF3, FST among others). A deeper and more comprehensive understanding of the RD network in the BMP pathway requires further careful studies and computational platforms that enable studying multiple nodes in RD networks will be very valuable[14].

In Nodal-permissive experimental conditions, BMP4 treatment of geometrically confined hPSC colonies results in a gradient that downregulates sharply (**Fig. S10B-C**)[24–26]. This has led to a proposition that in BMP4 treated geometrically confined hPSC colonies, BMP signalling is active exclusively at the colony periphery and inactive everywhere else – a spatial profile that can be modelled as a step-function along the colony radius[71]. These authors claim that this apparent step-function-like response in the pSMAD1 activity occurs due negative feedback enforced on BMP signalling at two different levels. First, by a BMP signalling mediated upregulation of BMP inhibitors[24,26], and second, due to a cell density mediated re-localization of BMP receptors from being present apically to becoming localized at basolateral regions, rendering them inaccessible for ligand mediated activation, everywhere in the colony except for the periphery [26]. Consistent with their proposed model, we also identify a valuable role of BMP inhibitors in orchestrating the pSMAD1 signalling gradient. However, our data indicate that the underlying mechanism regulating the pSMAD1 self-organization is inconsistent with a wide-spread dampening of BMP signalling owing to inaccessible BMP receptors. In this study, in addition to the unambiguous RD-like spatial expression patterns of pSMAD1 and differentiation markers like GATA3, in large colonies (3mm diameter), we identify experimental conditions that result in prominent expression of pSMAD1 at the periphery and the center of hPSC colonies of 500μm diameter when treated with 25ng/ml of BMP4 for 24h – observations consistent with RD-like behaviour (**Fig. 5D**). Neither of these observed expression profiles would arise in conditions where BMP receptors were inaccessible everywhere except for the colony periphery. Additional insight has been provided by Xue et al [27] where it was demonstrated that when treated with BMP4 for an extended period of time (4 days), all cells in the hPSC colony expressed nuclear-localized pSMAD1 (Fig 5A-B in ref. 27). We propose an alternative hypothesis for both the observed density dependent dampening, and the apparent step-function-like activity of BMP signalling observed in this system that invokes the involvement of Nodal signalling. Nodal is known to have a community effect whereby endogenous Nodal levels become more pronounced at higher cell densities[72,73], which would upregulate BMP antagonists downstream of Nodal signalling[52]. We argue that at increasing cell densities in the differentiating hPSC colonies, along with an increase in levels of BMP antagonists like Noggin and Gremlin family proteins owing to more cells secreting these inhibitors, there would likely be a dramatic increase in the levels of BMP antagonists like CERL, GDF3, FST which would upregulate due to pronounced levels of a community-effect mediated increase in endogenous Nodal. Abrogation of the fate patterning in this system in response to siRNA mediated inhibition of CERL and LEFTY (Fig 6 in ref 24) – an experimental condition that in principle should neither interfere with Noggin expression or change colony density, also supports our interpretation. Finally, our study provides direct evidence for the involvement of Nodal signalling in the apparent step-function-like response in pSMAD1 expression along the colony radius. When colonies of 500μm diameter were treated with 25ng/ml of BMP4 and SB, the signalling gradient formed showed no step-like response in the spatial pSMAD1 profile. Instead the signalling gradient gradually decreased in strength (as indicated by pSMAD1 fluorescence levels) from the colony periphery to the colony center (**Fig 5A-D**). However, in the presence of Nodal signalling, the pSMAD1 signalling gradient, indeed, downregulated sharply – as expected from a step-function-like response. This is consistent with the proposition that in regions within the differentiating geometrically-confined hPSC colonies that have active Nodal signalling, which have been shown to be immediately interior to the peripheral cells[26], there would be heightened levels of BMP antagonists like CERL causing a dramatic reduction of BMP signalling.

Notably, in our study, we report different aspects that can result in variability in experimental results when studying the stereotypic RD-like periodic response in BMP signalling in the hPSC context. One such source of variability is the level of endogenous Nodal signalling between different hPSC lines (**Fig. 3C**). Given the role that Nodal signalling plays in the formation of the pSMAD1 gradient (**Fig. 5, S10**), and the critical role it plays in ensuring the peri-gastrulation-associated fate patterning[24], the variability in endogenous levels of Nodal signalling can cause inconsistent responses between different cell lines and culture conditions. Importantly, a recent study has also shown drastically different responses in endogenous Nodal activation within the same hPSC line when cultured under different conditions for routine maintenance[74] – highlighting that even culture conditions for routine hPSC maintenance can have an effect in the response in the peri-gastrulation-like assay. Secondly, we observed that when geometrically confined hPSC colonies of 3mm diameter were treated with a high dose of BMP4 in SR medium, the stereotypical RD-like spatial periodicity of either pSMAD1 or GATA3 were not readily observed. However, changing the medium to an N2B27 based medium rescued these periodic responses. A key component of SR medium is Knockout Serum Replacement (KSR) which is known to contain lipid associated proteins like lysophosphatidic acid (LPA), and although the mechanism of action remains unclear, molecules like LPA have been shown to have an inhibitory effect on hPSC differentiation[57,58]. Given the above caveats associated with *in vitro* experiments, studies aimed at investigating the details of the RD network in BMP signalling in the hPSC context – especially those directed toward investigating the specifics of the spatial periodicity of morphogen activity and fate patterning, would benefit from removing these sources of variability between hPSC lines. Employing basal medium like N2B27 which is devoid of components like AlbmaxII and LPA, avoiding undefined media like those conditioned on MEFs, and removing Nodal signalling from their system by SB supplementation represent experimental conditions better suited for these studies.

#### Positional information mechanisms

In this study, we report that differentiating hPSCs (either culturing EBs in an FBS containing medium, or treatment of geometrically confined colonies with BMP4) upregulate a gastrulation-associated expression profile when endogenous Nodal signalling is active. However, in the case where Nodal signalling is downregulated, the same differentiation pulse upregulates a neurulation-associated gene expression profile. In addition, we demonstrated that perturbing Nodal signalling during BMP4 treatment of geometrically-confined hPSC colonies can result in a PI mediated interpretation of the emergent pSMAD1 signalling gradient into peri-gastrulation-associated or pre-neurulation-associated fate patterning. Notably, inducing the pre-neurulation-like fates did not require an initial differentiation toward the ectodermal lineage prior to inducing a pSMAD1 signalling gradient within the colonies. This observation, although apparently contradictory from a developmental point of view, is consistent with previous *in vitro* studies. For instance, the neuromesodermal precursors – a developmental population that arises during late gastrulation and resides in the node-streak border, caudal lateral epiblast, and the chordoneural hinge sections in the posterior end of the elongating embryo[75–77], can be derived from hPSCs through a transient 2.5 day pulse of wnt and bFGF[76]. In addition, similar to our observations, Funa *et al* employed activation of wnt signalling in hPSCs and identified a dissection of the primitive streak and the neural crest fates in the same assay durations[48]. These results suggest that genomic accessibility in hPSCs likely does not dictate the ability of inducing these early developmental fates.

Funa *et al* employed a chromatin-immunoprecipitation sequencing (CHIP-seq) study and identified that β-catenin is able to directly regulate both the primitive streak, and the neural crest genes[48]. However, expression of the primitive streak genes requires β-catenin to form a physical complex with SMAD2/3 – the effectors of Nodal signalling. Furthermore, upon the formation of the complex, the expression of genes associated with the neural crest fate were inhibited. The mechanism by which SMAD2/3 can prevent β-catenin mediated activation of the neural crest genes remains unclear. We hypothesize that a similar mechanism with BMP signalling could explain much of our data. Much like β-catenin, SMAD1 may activate a peri-gastrulation-associated gene profile in the presence of SMAD2/3 whereas in the absence of SMAD2/3, SMAD1 may activate pre-neurulation-associated genes. Future studies that employ a similar approach to Funa *et al* by performing CHIP-seq studies to identify the binding dynamics of SMAD1 in the presence and absence of Nodal signalling can provide valuable insights toward a molecular understanding of how SMAD1 regulates the expression of the gastrulation versus neurulation associated gene profiles.

### Biochemical versus biomechanical regulation of pSMAD1 self-organization

In a recent elegant study, Xue et al demonstrated that a pSMAD1 gradient can also arise within the geometrically confined hPSC colonies during extended culture (9 days) in medium that was supplemented with low levels of BMP inhibitors and SB[27]. Notably, these culture conditions in their study ensured that the differentiating hPSCs were subjected to minimal amounts of BMP ligands during differentiation. Under these conditions, the authors demonstrated that biomechanical characteristics of the cells in the colony like size and contractile forces exerted could autonomously activate BMP signalling at the cells in the colony periphery[10,27]. In addition, the authors demonstrated that if provided a transient, 24h long, pulse of wnt activation, the colonies can give rise to regionalized fates associated with the neural plate, and the neural plate border[27]. Furthermore, they show that siRNA mediated Noggin inhibition did not significantly alter the patterning of the neuroectodermal fates. Although their observations provide an additional mechanism by which a signalling gradient could self-organize during development – a biomechanical paradigm, they do not contradict our findings. This is because the mechanism proposed by Xue et al functions under conditions where the hPSCs are subjected to low/minimal levels of BMP ligands whereas our model considers conditions where the medium is supplemented with BMP4. In fact, when the authors added BMP ligands in their induction media over the course of their assay, they observed widespread activation of pSMAD1 activity and abrogation of the expression of the neural plate marker. This highlights the presence of a signalling hierarchy in platforms that begin with 2D cell populations, where biochemical signals may overwhelm biomechanical systems. However, in 3D starting populations and in vivo, the relationship between the biochemical and biomechanical cues is likely far more balanced and it would be interesting to speculate whether these two paradigms could perform redundant functions to ensure developmental robustness at varying levels of presence of BMP ligands. Finally, the fact that Xue et al observe the regionalization of the neural plate and neural plate border identities after extended culture durations provides further evidence to the proposition that the fate patterning occurs in a manner consistent with PI as at reduced levels of BMP signalling, the fate patterning would be predicted to arise after longer durations. In addition, under these conditions where BMP signalling has not activated at sufficiently high levels, consistent with the PI model, the authors did not observe fates associated with the non-neural ectodermal identity.

## Conclusions

In conclusion, we report characterization of a high-throughput microtiter plate that enables robust geometric confinement of a variety of cell types. We employ this platform to screen hPSC lines for their ability to induce gastrulation-associated fate patterning and observe a Nodal-dependent response in the efficiency of BRA (a gastrulation-associated fate) induction, thereby providing a proof of principle of the ability of this platform to be employed for high-throughput screening experiments. In addition, we identify that differentiating hPSCs upregulate either gastrulation, or neurulation associated gene profiles in a Nodal signalling dependent manner. Further, we demonstrate that in BMP4 treated geometrically-confined hPSC colonies, Nodal signalling can affect the RD mediated self-organization of pSMAD1 – the downstream effector of BMP signalling; and that it also regulates the switch between peri-gastrulation-like and pre-neurulation-like identities in PI mediated fate patterning occurring within the differentiating colonies. Finally, consistent with a previous study that investigated peri-gastrulation-like fate patterning in BMP4 treated hPSC colonies, we demonstrate that the pre-neurulation-like fate patterning follows a stepwise model of RD and PI, hinting at possible conservation of the underlying mechanism that regulates differentiation of the epiblast and ectoderm in human development.

## Materials and Methods

### Human Pluripotent Stem Cell Culture

CA1 human embryonic stem cell line was provided by Dr. Andras Nag7y (Samuel Lunenfeld Research Institute). H9-1 was provided by Dr. Sean Palecek (University of Wisconsin – Madison). H9-2, HES2, and MEL1 (PDX1-GFP) were provided by Dr. Gordon Keller (McEwen Centre for Regenerative Medicine/University Health Network). HES3-1, and HES3-2 were provided by Dr. Andrew Elefanty (Monash University). H1, H7, H9-3 were acquired from WiCell Research Institute. For routine maintenance, CA1, H9-1, and H9-2 were cultured on Geltrex (Life Technologies, diluted 1:50) coated 6-well tissue culture plates using mTeSR1 medium (StemCell Technologies) as per manufacturer’s instructions. The cells were passaged at a ratio of 1:12 using ReleSR (StemCell Technologies) per manufacturer’s instructions. For the first 24h after passage, the cells were cultured in ROCK inhibitor Y-27632 to increase cell viability. The medium was changed every day and passaged every 4-5 days or when the cells reached 75-80% confluence. For routine maintenance, H1, H7, H9-3, HES3-1, HES3-2, MEL1, HES2 were cultured on feeder layers of irradiated MEFs in Dulbecco’s Modified Eagle’s Medium (DMEM) (Invitrogen), 1% Penicillin/Streptomycin, 1% non-essential amino acids, 0.1mM β-mercaptoethanol, 1% Glutamax, 2% B27 minus retinoic acid, 20% KnockOut serum replacement **(referred to as ‘SR’ medium)** and supplemented with 20 ng ml–1 FGF-2 (PeproTech). H1, H7, and H9-3 cells were passaged 1:6 every 4–5 days and were disassociated into small clumps using 0.1% collagenase IV (Invitrogen). HES3-1, HES3-2 were passaged 1:24 every 4–5 days and dissociated using TryplE Express (Invitrogen). All cell lines were confirmed negative for mycoplasma contamination.

### Preparation of PEG plates

Platform set up, and XPS studies were performed using 22mmx22mm borosilicate coverslips (Fisher Scientific), and the 96-well plate platform was developed using custom sized (110mmx74mm) Nexterion-D Borosilicate thin glass coverslips (SCHOTT). The glass coverslips were activated in a plasma cleaner (Herrick Plasma) for 3 minutes at 700 mTorr and incubated with 1 ml of Poly-L-Lysine-grafted-Polyethylene Glycol (PLL-g-PEG(5KD), SUSOS,) at a concentration of 1 mg/ml at 37°C overnight. The glass slides were then rinsed with ddH2O and dried. The desired patterns were transferred to the surface of the PEG-coated side of the coverslip by photo-oxidizing select regions of the substrate using Deep UV exposure for 10 minutes through a Quartz photomask in a UV-Ozone cleaner (Jelight). Bottomless 96-well plates were plasma treated for 3 minutes at 700 mTorr and the patterned slides were glued to the bottomless plates to produce micro-titer plates with patterned cell culture surfaces. Adhesives validated for biocompatibility standards ISO10993, and USP Class VI were utilized for the assembly of the plates. Prior to seeding cells onto the plates, the wells were activated with N-(3-Dimethylaminopropyl)-N’-ethylcarbodiimide hydrochloride (Sigma) and N-Hydroxysuccinimide (Sigma) for 20 minutes. The plates were thoroughly washed three times with ddH2O, and incubated with Geltrex (diluted 1:150) for 4h at room temperature on an orbital shaker. After incubation, the plate was washed with Phosphate Buffered Saline (PBS) at least three times to get rid of any passively adsorbed extracellular matrix (ECM) and seeded with cells to develop micro-patterned hPSC colonies.

### Comparison between PEG plates with μCP plates

PEG plates (as described above) and μCP plates (as reported previously[30]) were generated with patterned islands of 200μm in diameter with 500μm separation between adjacent colonies. A single cell suspension of CA1s was generated by incubating in 1ml of TryplE (Invitrogen) per well for 3 minutes at 37°C. The TryplE was blocked using in equal volume SR medium (see ‘Human pluripotent stem cell culture’ section above for composition) and the cells were dissociated by pipetting to generate a single cell suspension. The cells were centrifuged into a pellet and the supernatant aspirated to remove any residual TryplE. A single cell suspension was then generated in SR medium supplemented with 10μl of ROCKi and 20ng/ml of bFGF at a cell density of 500,000 cells/ml and 100μl of the suspension was plated onto the PEG and μCP plates for a period of 2-3h till robust cell attachment was observed. The cells were then left to make a confluent colony overnight (~12h). Once confluent colonies were observed, the differentiation was performed using 100μl per well of the following inductive conditions in Apel (Stem Cell Technologies) basal media for 48h – bFGF (40ng/ml) +SB431542 (10μM) to induce differentiation into ectodermal fates, BMP4 (10ng/ml) +ActivinA (100ng/ml) to induce mesendodermal differentiation, BMP4 (40ng/ml) to induce extra-embryonic/’other’ fates, and as controls, Nutristem, and basal Apel media were used. After 48h, the colonies were fixed, and stained for OCT4 and SOX2. The relative percentages of the colonies that were positive for the two markers were used to identify the early fates induced within the colonies. A detailed description of the assay has been previously reported[30].

### Peri-gastrulation-like and pre-neurulation-like fate patterning induction

Except for the experiments where we demonstrated the spatial oscillations of pSMAD1 and the pre-neurulation-like fates in 3mm diameter colonies, all fate patterning studies were performed in SR medium (see ‘Human pluripotent stem cell culture’ section above for composition) supplemented with 100ng/ml of bFGF. The studies where we demonstrate the spatial oscillations of the morphogen activity and fate patterning were performed in N2B27 medium. N2B27 medium was composed of 93% Dulbecco’s Modified Eagle’s Medium (DMEM) (Invitrogen), 1% Penicillin/Streptomycin, 1% non-essential amino acids, 0.1mM β-mercaptoethanol, 1% Glutamax, 1% N2 Supplement, 2% B27 Supplement minus retinoic acid.

The hPSC lines that were cultured in feeder-dependent techniques for routine maintenance were first feeder depleted by passaging the cells at 1:3 on geltrex and cultured on Nutristem. To seed cells onto ECM-immobilized PEG-UV 96-well plates, a single cell suspension of the hPSC lines was generated as described above. The cells were centrifuged and re-suspended at a concentration of 1 × 10^6^ cells/ml in SR medium supplemented with 20ng/ml bFGF (R&D) and 10μM ROCK inhibitor Y-27632. Wells were seeded in the PEG-patterned 96 well plates at a density of 60,000 cells/well for plates with colonies of 500μm diameter, 80,000 cells/well for colonies of 1mm diameter, and at 120,000 cells/well for plates with colonies of 3mm diameter and incubated for 2-3h at 37°C. After 2-3h, the medium was changed to SR without ROCKi. When confluent colonies were observed (12-18h after seeding), the peri-gastrulation-like induction or pre-neurulation-like induction was initiated as follows. A) Peri-gastrulation-like induction (**Fig. 2**) was performed in SR medium supplemented with 100ng/ml of bFGF (R&D) and 50ng/ml of BMP4. B) Unless otherwise stated, pre-neurulation-like induction with 500μm colonies was performed with SR medium (see ‘Human pluripotent stem cell culture’ section above for composition) supplemented with 100ng/ml of bFGF with 25ng/ml of BMP4, and μM SB431542 (‘SB’). C) Endoderm fingerprinting assay (Fig. S6) was performed with N2B27 medium supplemented with 25ng/ml of Wnt3A and 50ng/ml of ActivinA. D) RD-like periodic pattern induction of pSMAD1 activity, and pre-neurulation-like fates was tested in both SR, and N2B27 mediums. In the case of SR, the medium was supplemented with 10μM SB, 100ng/ml of bFGF, and either 50ng/ml or 200ng/ml of BMP4. In the case of N2B27, the medium was supplemented with 10μM SB, 10ng/ml of bFGF, and either 50ng/ml of 200ng/ml of BMP4.

### Embryoid body differentiation assay

The differentiation media for the EB assay contained 76% DMEM, 20% Fetal Bovine Serum (FBS), 1% Penicillin/Streptomycin, 1% non-essential amino acids, 0.1mM β-mercaptoethanol, 1% Glutamax, (all Invitrogen). A large volume of the medium was prepared with a single batch of FBS and frozen at −80C and was used to differentiate the EBs made from all the hPSC lines tested. EB formation from the hPSC lines was achieved by generating a single cell suspension (as described in the section above) directly in the differentiation media supplemented with ROCKi for the first day. The cell suspension was then plated on 24-well microwell plates (Aggrewell – 400μm, Stem Cell Technologies). The seeding density was chosen to allow generation of size-controlled EBs (~500cells/EB) for all hPSC lines. The media was carefully replaced with differentiation media without ROCKi 24hours after seeding to ensure that the EBs were not disturbed. EBs were harvested from the Aggrewell plates each day by adding 1 ml of DMEM into the wells and pipetting till the EBs lifted off from the microwells, and frozen as a pellet at −80C till gene expression was assessed using qPCR.

### CA1 Nog^-/-^ cell line generation

CA1 Noggin knock out lines were generated using a CRISPR/Cas9 mediated donor-free dual knock-out using a previously described strategy[78]. The sgRNA design was performed with CRISPRko Azimuth 2.0 (Broad Institute) using human Noggin (NCBI ID9241) as entry data and SpCas9 for the nuclease. The software ranks sgRNAs with high on-target activities and low off-target activities in a combined rank[79]. We chose sgRNA1 (5’-CTGTACGCGTGGAACGACCT-3’) and sgRNA2 (5’– CAAAGGGCTAGAGTTCTCCG-3’) with a combined rank of 4 and 1, respectively. sgRNA1 & 2 can be used individually or applied together to produce Noggin knock-out. Latter leads to a deletion of a DNA fragment of 112 bp and to a predetermined stop codon (**Fig. S8A**).

Transfection and evaluation of cutting efficiency: We first evaluated the cutting efficiency of SpCas9 for each individual gRNA on a population level. For this, we seeded CAI hESC into 24-well plates such that they are 50-60 % confluent at the day of transfection (approx. 24h after seeding). CmgRNA were generated by mixing μM AltR CRISPR crRNA (IDT, custom oligo entry) with μM AltR CRISPR tacrRNA (IDT, Cat. 1073189), annealed at 95°C for 5 min and cooled down at room temperature. GeneArtTMPlatinumTM Cas9 Nuclease (Invitrogen, B25641) was diluted to μM using Opti-MEM (Thermo Fisher Scientific, 31985062). Cas9 and cmgRNA were mixed at a concentration of 0μM each in 25μl of OptiMEM. After incubation at room temperature for 5 minutes, 1 μl of EditProTM Stem (MTI Globalstem) diluted in 25μl of Opti-MEM was added to the Cas9/cmgRNA complex and incubated for 15 minutes at room temperature. Before adding the reagent-Cas9/cmgRNA mix to the cells, medium was replaced with 500μl / 24 well of fresh mTeSR. Medium was replaced 24h after transfection. 48h after transfection, cells were harvested by incubation in Gentle Cell Dissociation Reagent (STEMCELL Technologies, 07174) for 7 min. Dissociation reagent was removed and cells were resuspended in cultivation medium, pipetted to single cells and spin down for 5min at 200g. Cells were resuspended in 25μl of Cell Lysis Buffer mixed with μl Protein Degrader, both from the GeneArtTM Genomic Cleavage Detection Kit (Invitrogen, A24372). Cells were lysed at 68°C for 15min, 95°C for 10min and kept on ice. PCR was performed using Phusion High Fidelity DNA Polymerase (NEB, M0530) according to manufactures protocol using 2μl of the cell lysate. Primer for the PCR were the following: (fwd) 5’CTACGACCCAGGCTTCATGGC’3, (rev) 5’GACGGCTTGCACACCATGC3’. PCR product of un-transfected and transfected samples were analyzed on 2.5% MetaPhore Agarose Gel (Lonza) PCR products were analyzed using GeneArt Genomic Cleavage Detection Kit (Invitrogen, A24372) according to manufacturer’s protocol. The cleaved and un-cleaved samples were loaded on 2.5% MetaPhore Agarose Gel (Lonza) and the bands were analyzed using ImageJ. Percentage of gene modification was calculated as described in a previous report (**Fig. S8B**)[80]. Additionally, PCR products were send for Sanger Sequencing. Chromatograms were analyzed using TIDE[81].

Cell line generation: The cell line was generated using gRNA1 & 2 mixed with Cas9 at 0.3μM each. The transfection was proceeded with exact same protocol as described above using 6 × 24 wells. After three days, cells reached confluency and were seeded to 6 well plates at sufficiently low densities to achieve clonal growth from single cells. Approximately 7 days after seeding, single clones were picked and transferred to 96 well plates. 24 clones were expanded for 2 passages and PCR was performed on cell lysates as described above (**Fig. S8C**). PCR products were send for Sanger Sequencing and aligned to (NCBI ID9241) and to untransfected wildtype sequence (**Fig. S8C**). Clones with clear loss of function mutations in both alleles (C1 and C7) were further characterized for their pluripotency marker expression (**Fig. S8D**).

### Quantitative PCR analysis

RNA extraction for all gene expression analysis studies was performed using Qiagen RNAeasy miniprep columns according to the manufacturer’s protocol, and the cDNA was generated using Superscript III reverse transcriptase (Invitrogen) as per the manufacturer’s instructions. The generated cDNA was mixed with primers for the genes of interest and SYBR green mix (Roche, Sigma) and the samples were run on an Applied Biosystems QuantStudio 6 flex real-time PCR machine. The relative expression of genes of interest was determined by the delta–delta cycle threshold (ΔΔCt) method with the expression of GAPDH as an internal reference. Primer sequences used are provided in Supplementary Information (**Table S2**).

### Immunofluorescent staining, and image analysis

After the peri-gastrulation-like or the pre-neurulation-like induction was completed, the plates were fixed with 3.7% paraformaldehyde for 20 min, rinsed three times with PBS and then permeabilized with 100% methanol for 3 min. After permeabilization, the patterned colonies were blocked using 10% fetal bovine serum (Invitrogen) in PBS overnight at 4°C. Primary antibodies were incubated at 4°C overnight (antibody sources and concentrations are shown in **Table S3**). The following day, the primary antibodies were removed, and the plates were washed three times with PBS followed by incubation with the secondary antibodies and DAPI nuclear antibody at room temperature for 1 h. Single-cell data were acquired by scanning the plates using the Cellomics Arrayscan VTI platform using the ‘TargetActivation.V4’ bioassay algorithm. This algorithm utilizes the expression intensity in the DAPI channel to identify individual nuclei in all fields imaged and acquires the associated intensity of proteins of interest localized within the identified region. As previously described[24], single-cell data extracted from fluorescent images were exported into our custom built software, ContextExplorer (Ostblom et al, unpublished), which classifies cells into colonies via the DBSCAN algorithm. Cartesian coordinates relative to the colony centroid are computed for every cell within a colony. Hexagonal binning is used to group cells from multiple colonies according to their relative location within a colony. Average protein expression of cells within a bin is represented by the color map, which is normalized to the lowest and highest expressing hexagonal bins. In the line plots of spatial expression trends, cells are grouped in annular bins according to the Euclidean distance between a cell and the colony centroid. For each colony, the mean expression of all cells within an annular bin is computed. The average of all the colony means is displayed in the line plot together with the standard deviation and the 95% confidence interval (CI).

## Figure Legends

**Figure S1:**
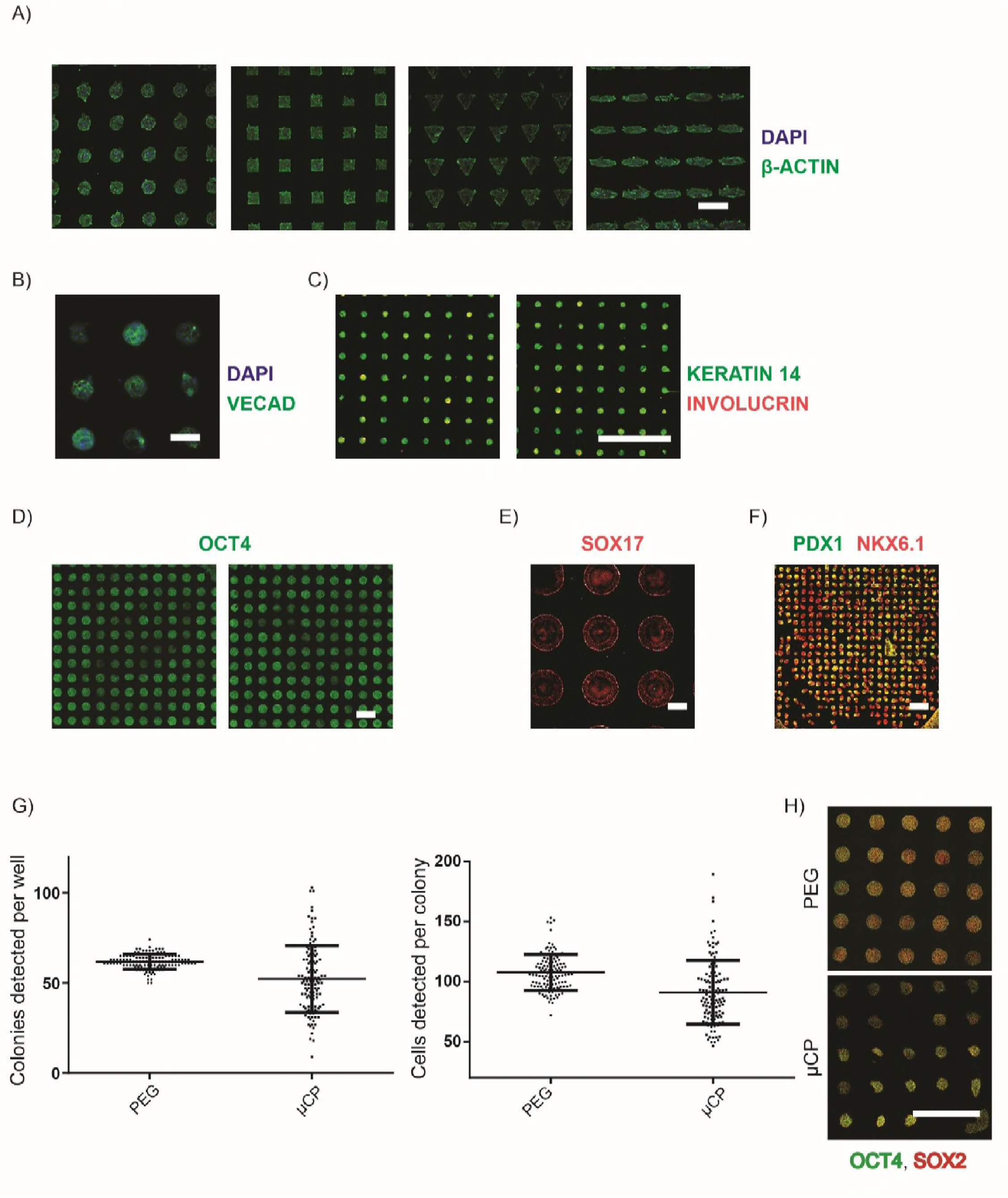
Characterization of PEG plates. A-F) Representative images acquired on PEG plates for multiple cell types. A) Mouse embryonic fibroblasts (MEFs) stained for β-actin in green, and DAPI in blue. B) Hemogenic endothelial cells stained for VECAD in green and DAPI in blue. C) Primary human keratinocytes stained for Keratin 14 in green and Involucrin in red. D) Mouse embryonic stem cells stained for OCT4 in green. E) BMP4 treated human induced pluripotent stem cells stained for SOX17 in red. F) Human endodermal progenitor cells allowed to generate outgrowths stained for NKX6.1 in red and PDX1 in green. G-H) Comparison of patterning response on PEG plates vs μCP plates. G) Number of colonies identified per well between PEG and μCP plates. Each dot represents the number of colonies identified per well for 120 randomly chosen wells between the four replicates of PEG vs μCP plates. Number of cells identified per colony between PEG and μCP plates. Each dot represents the average number of cells per colony for 120 randomly chosen wells between the four replicates of PEG vs μCP plates. H) Representative images of hPSCs micropatterned in 96-well plates using PEG-based technique vs μCP.

**Figure S2:**
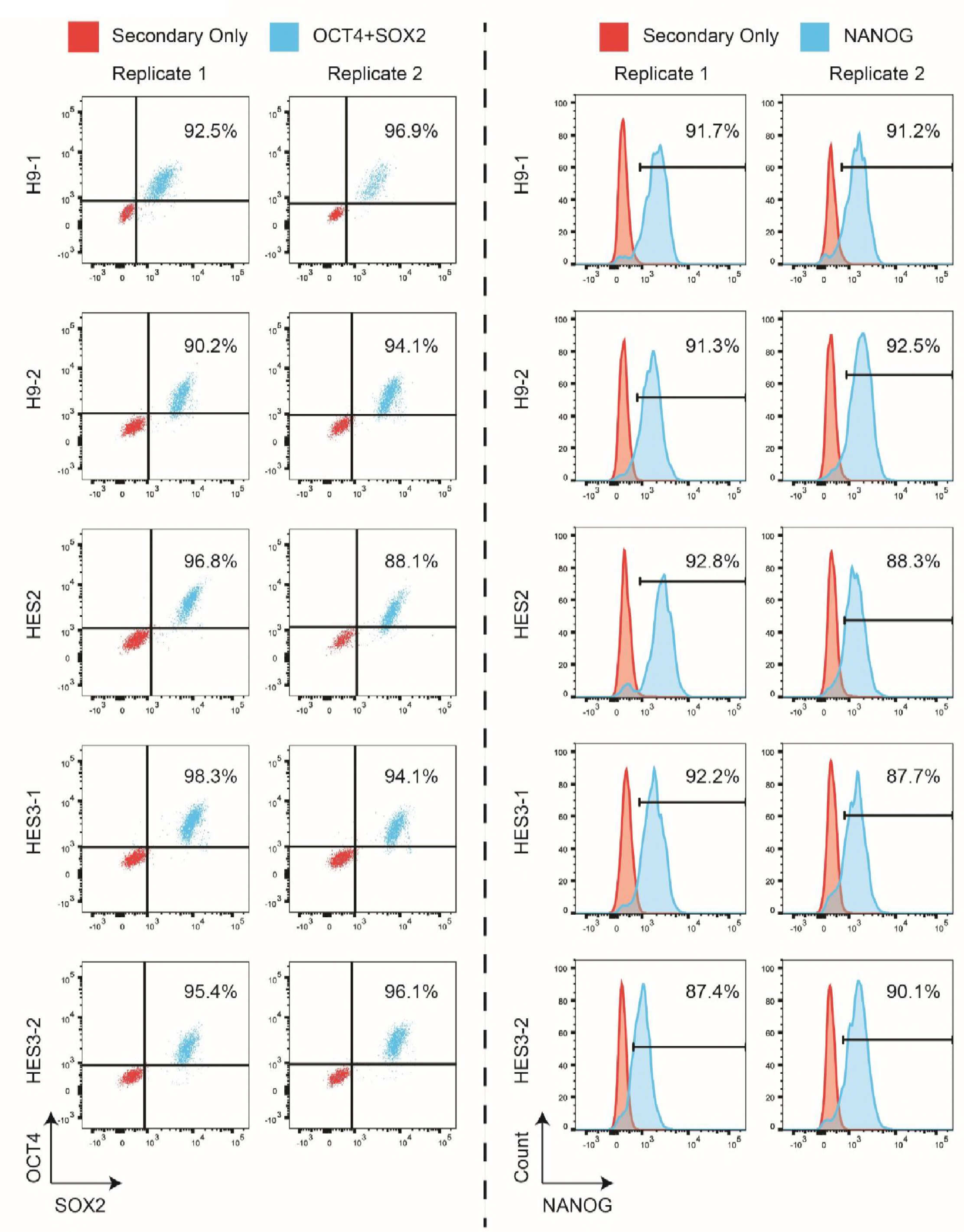
Starting populations of test hPSC lines show high expression of pluripotency associated proteins. FACS plots of OCT4, SOX2, and NANOG of starting populations of H9-1, H9-2, MEL1, and HES3-1.

**Figure S3:**
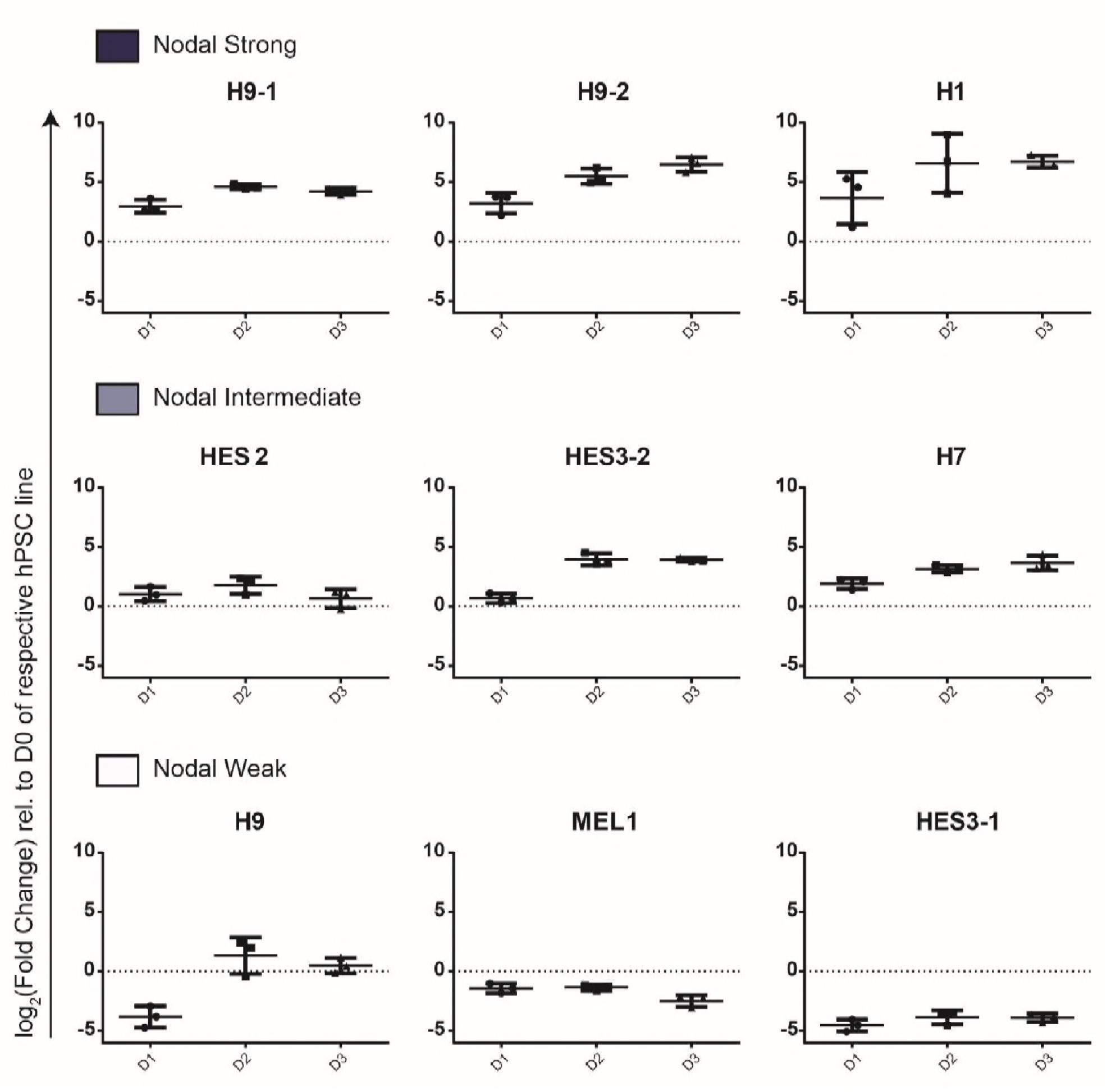
Nodal expression dynamics in FBS mediated non-specific differentiation of hPSC embryoid bodies. Temporal dynamics of Nodal for the test hPSC lines shown for the three clusters of Nodal-Strong, Nodal-Intermediate, and Nodal-weak (**Fig. 3Bii**). Each dot represents the detected expression level for a biological replicate. Bar plots represent mean ± s.d.

**Figure S4:**
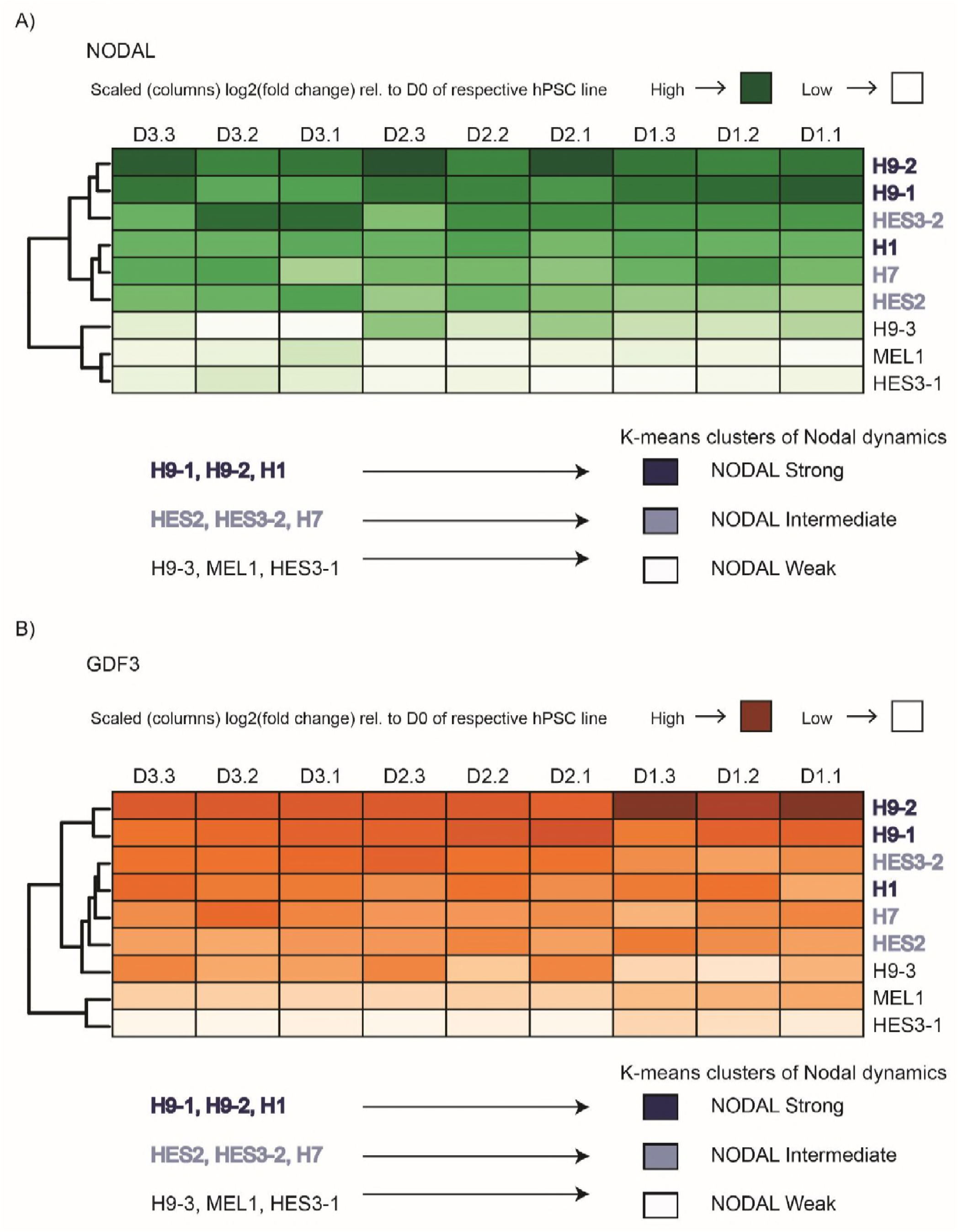
Hierarchical clustering of Nodal and GDF3 is consistent with unsupervised K-means clustering. A) Hierarchical clustering of the Nodal expression in the test hPSC lines based on Euclidian distance reveals similar clusters as the ones from unsupervised K-means clustering. B) Hierarchical clustering of the Nodal target (GDF3) expression in the test hPSC lines based on Euclidian distance reveals similar clusters as the ones from unsupervised K-means clustering of Nodal signalling.

**Figure S5:**
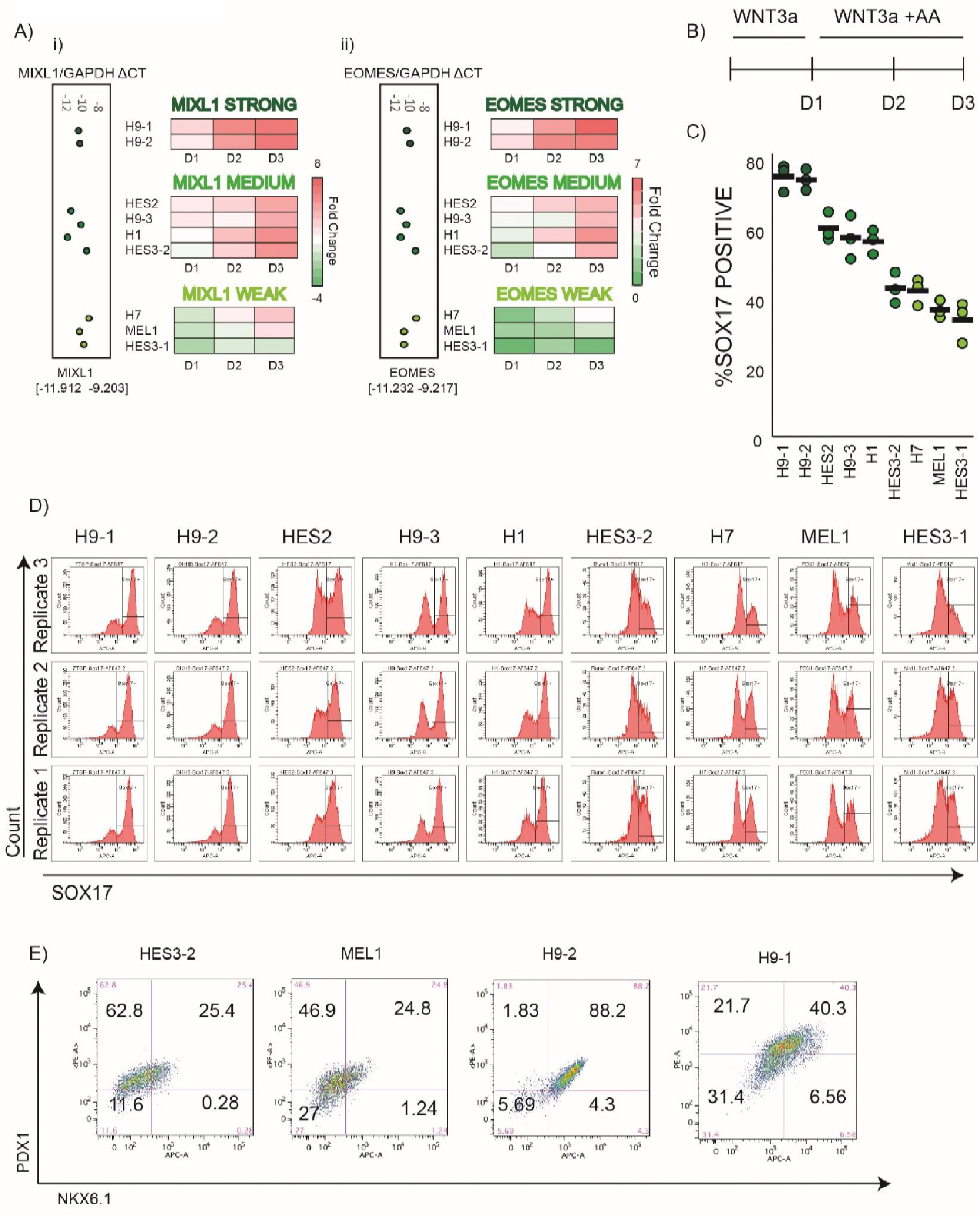
MIXL1 and EOMES dynamics during EB assay predict endoderm differentiation propensity of hPSC lines. A) Panel of hPSC lines clustered into three groups of ‘Strong’, ‘Medium’, and ‘Weak’ responders for (i) MIXL1, and (ii) EOMES from Fig 3Bii. The expression levels of MIXL1 and EOMES in the pluripotent state (Day 0) shown in the boxes adjacent to the heatmaps. B) Overview of the protocol for directed differentiation toward definitive endoderm. The cells were treated with Wnt3a from 0h-24h, and Wnt3a+ActivinA from 24h-72h. C-D) Efficiency of SOX17 induction in the test hPSCs using the protocol in B). C) Black dash denotes the mean of three independent replicates represented by the dots. D) FACS plots for individual replicates from C). E) FACS plots showing the efficiency of induction of pancreatic progenitors as indicated by the expression of PDX1, and NKX6.1 for hPSC lines in ‘Strong’ and ‘Weak’ clusters from A). The differentiation was performed using a previously described protocol[82]. The data are from one biological replicate.

**Figure S6:**
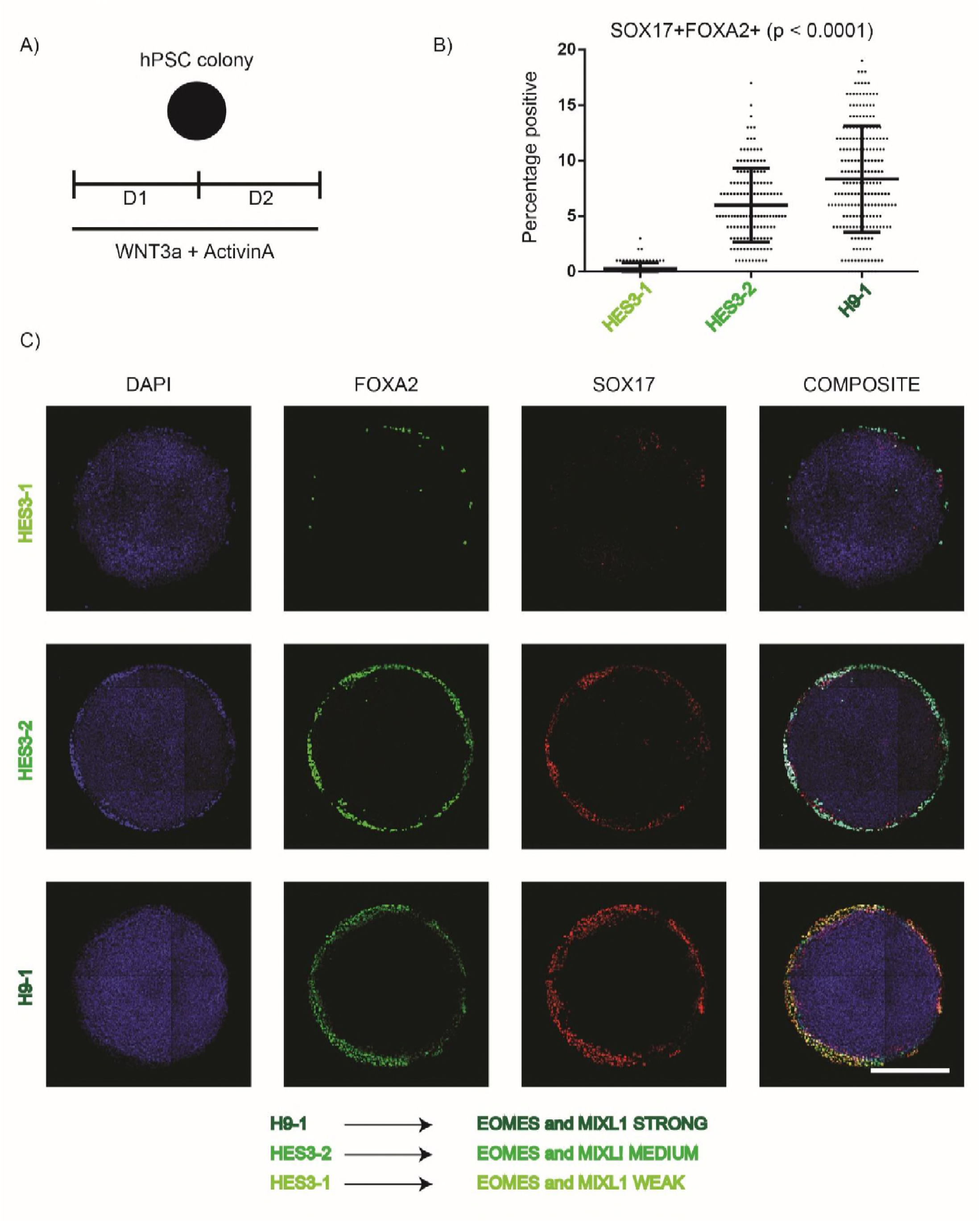
Peri-gastrulation-like assay predicts endoderm differentiation bias of hPSC lines. A) Overview of assay for predicting endodermal differentiation bias. Geometrically-confined hPSC lines were treated with Wnt+ActivinA for 48hours prior to fixation and staining. B) Quantified fraction of endodermal cells, defined as double positive for SOX17, and FOXA2, detected within the geometrically-confined hPSC colonies. The hPSC lines chosen were one each from the ‘Strong’, ‘Medium’, and ‘Weak’ clusters of MIXL1, and EOMES from **Fig. S5A**. Each data point represents an individual identified colony. Data pooled from two different experiments and represented as mean ± s.d.; p-values calculated using one-way ANOVA (Kruskal-Wallis test). C) Representative immunofluorescent images of colonies from B) stained for DAPI, SOX17, and FOXA2. Scale bar represents 500μm.

**Figure S7:**
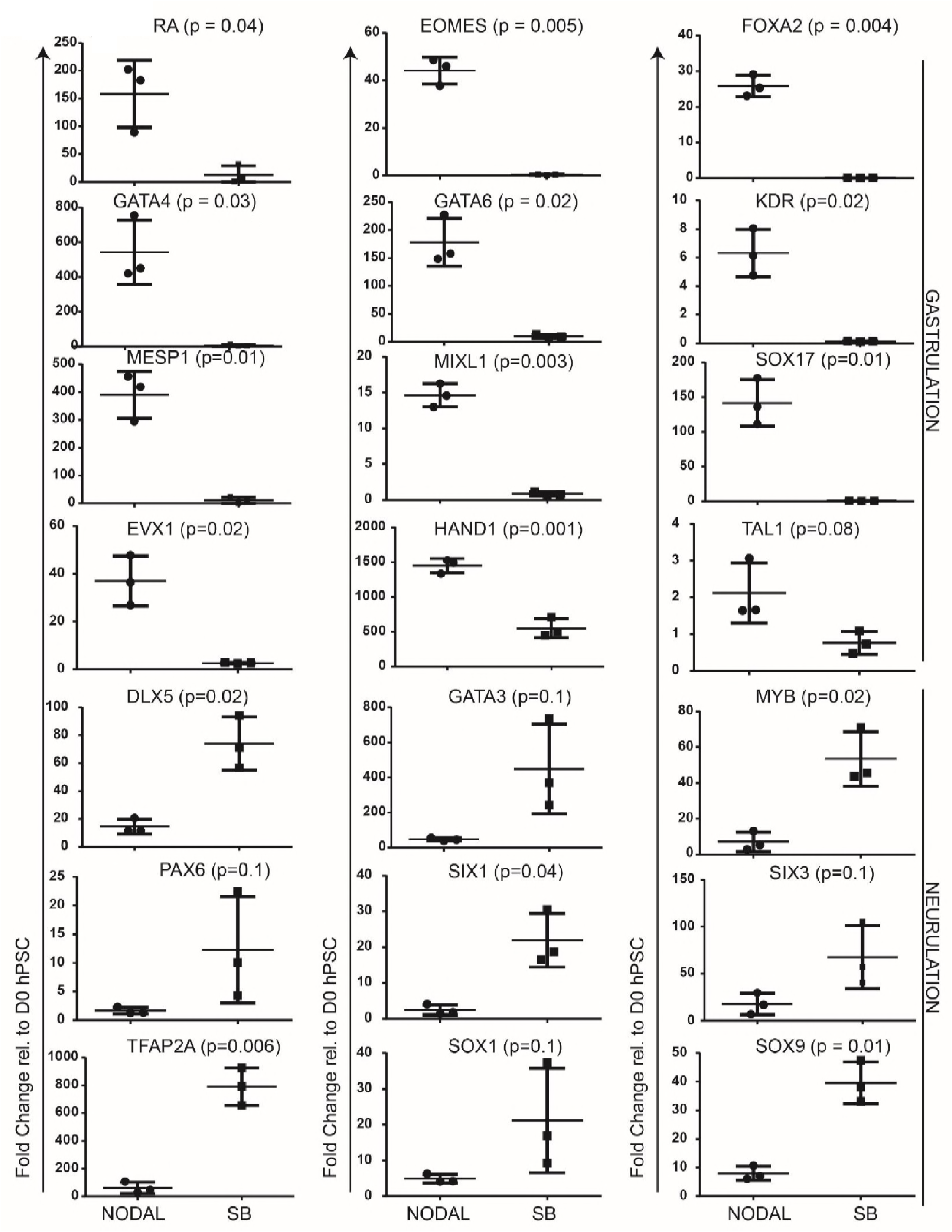
Modulation of Nodal signalling during BMP4 treatment of geometrically-confined hPSC colonies reveals a switch in expression of gastrulation vs neurulation associated genes. Response of modulation of Nodal signalling in expression of gastrulation versus neurulation associated genes in BMP4 treated geometrically-confined CA1 colonies (assay details in Fig 3Di). Individual data points represent biological replicates. Data shown as mean ± s.d., and p-values calculated using Mann-Whitney U test.

**Figure S8.**
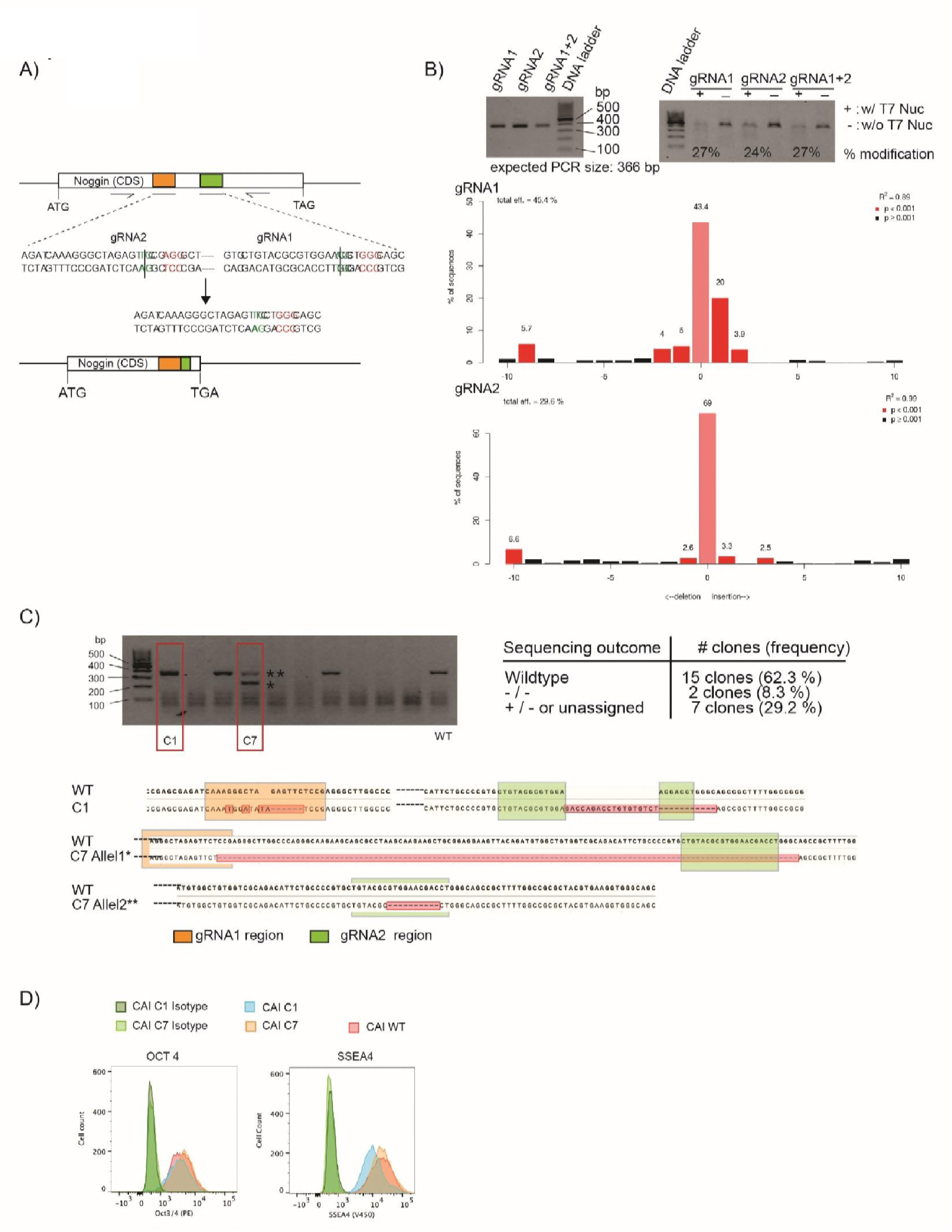
Generation of CA1 Noggin^-/-^ cell line using CRISPR/Cas9: A) Generation of CA1 Nog^-/-^ cell line using CRISPR/Cas9: a) Schematic of dual knock-strategy for Noggin (NCBI ID9241). Nucleotide sequence of gRNA binding regions (gRNA 1; green and gRNA2; orange) with expected cutting sites (green nucleotides) and resulting repaired DNA strand. The gRNAs have been selected such that repaired DNA will lead to a predetermined stop codon (TGA). B) Analysis of gRNA cutting efficiency on transfected hESC CAI population 48h post-transfection. Top left: PCR from transfected cell lysate. Top right: PCR fragments processed with and without T7 Endonuclease (Genomic Cleavage Assay). Bottom: Sanger sequencing of PCR product and subsequent decomposition and analysis using TIDE online analyzing tool. PCR from wildtype CA1 was used as control. Bar chart shows frequency, type, and position of mutations. C) A total of 24 clones were analyzed by PCR (top left) and by Sanger Sequencing (top right). Sequencing revealed 2 clones (C1 and C7) with homozygous knock-out sequences (bottom). The two bands (two alleles) of C7 were separately purified from Agarose gel before sequencing. D) Pluripotency marker staining of C1 and C7 lines of CA1s.

**Figure S9:**
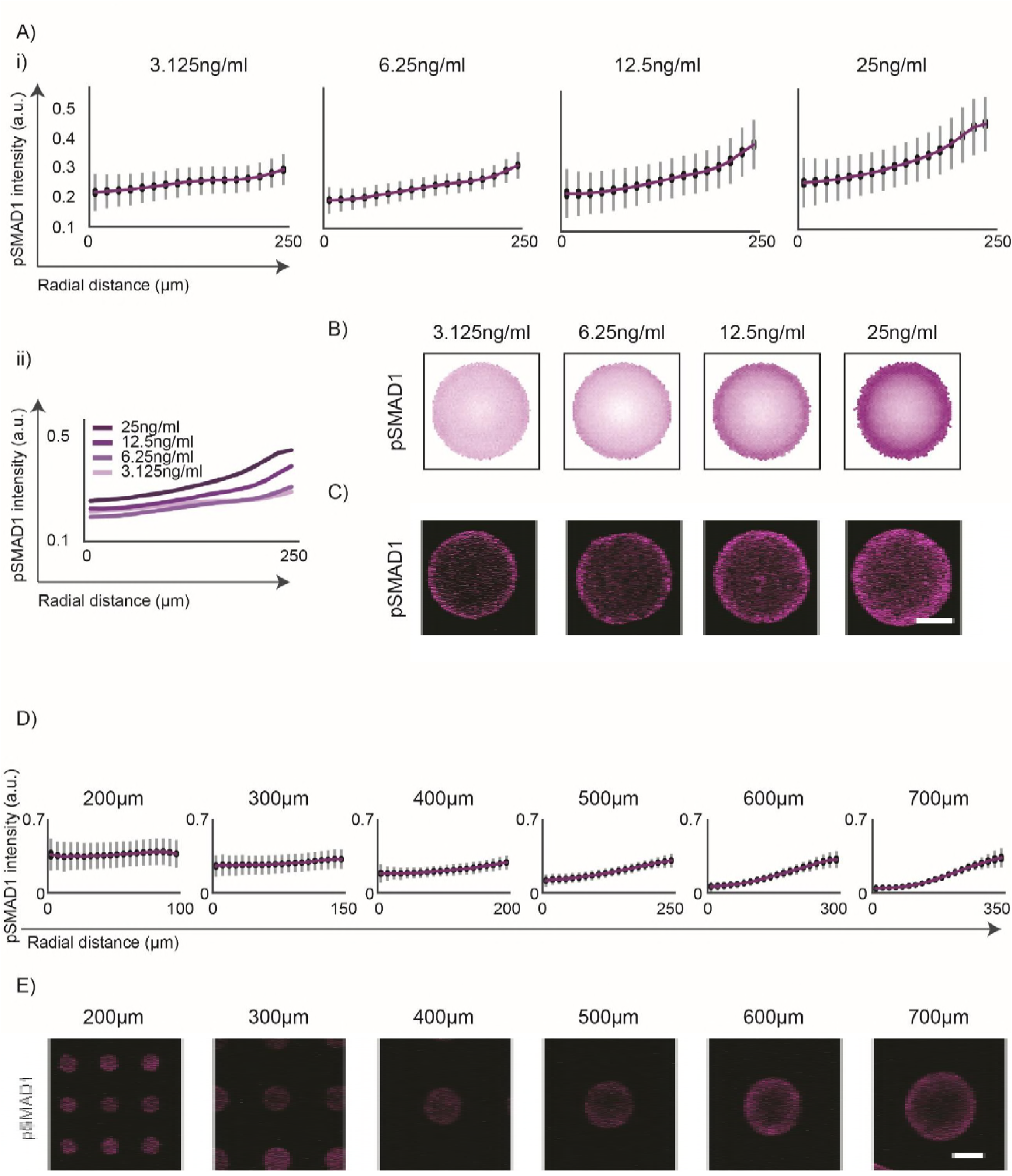
pSMAD1 gradient formation is consistent with a BMP4-Noggin RD network mediated self-organization. A) (i-ii) Radial gradient formed in colonies of 500μm diameter treated with varying doses of BMP4 in induction medium (3.125ng/ml, 6.25ng/ml, 12.5ng/ml, and 25ng/ml) represented as line plots. (i) The gradients shown individually. Data pooled from two experiments, and represent 299, 293, 302, and 343 colonies for the respective doses. Standard deviations shown in grey and 95% confidence intervals shown in black. (ii) Line plots shown in one graph for comparison of pSMAD1 levels at colony periphery. B) Average pSMAD1 expression levels shown as overlay of the detected colonies (numbers mentioned in A). C) Representative immunofluorescent images of pSMAD1 for respective conditions. Scale bars represent 200μm. D) Average pSMAD1 expression of 987, 528, 280, 182, 107, and 89 colonies for varying colony sizes (200μm, 300μm 400μm, 500μm, 600μm, and 700μm) treated with 25ng/ml of BMP4 in induction medium. Data pooled from two experiments. Standard deviations shown in grey and 95% confidence intervals shown in black. Scale bars represent 200μm.

**Figure S10:**
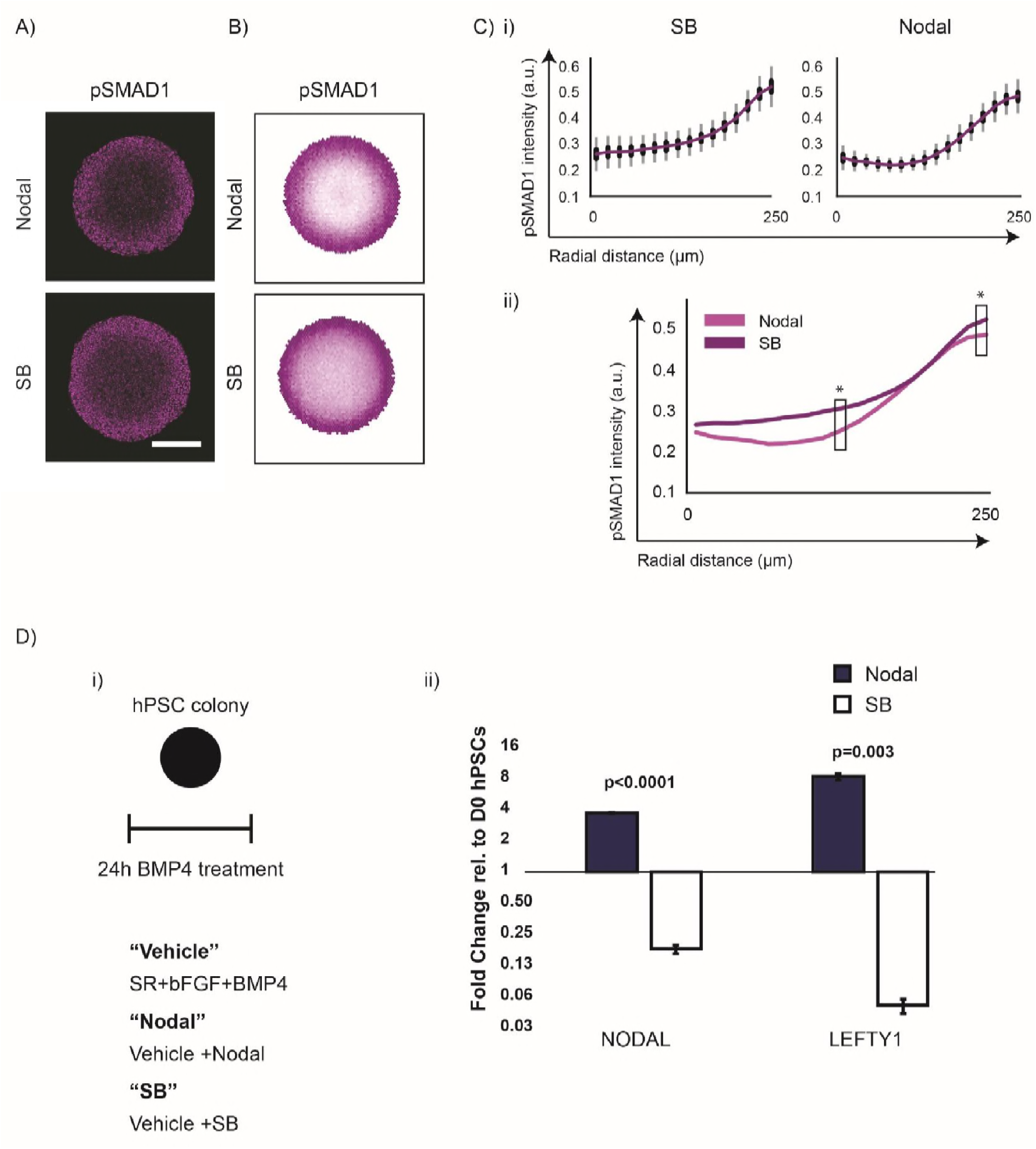
Nodal signalling contributes to the formation of the pSMAD1 gradient. A) Representative immunofluorescence images of 500μm diameter hPSC colonies stained for pSMAD1 after 24h of BMP4 treatment. Scalebar represents 200μm. B) Average pSMAD1 intensity represented as overlays of 368 and 411 colonies for Nodal and SB conditions respectively. Data pooled from two experiments. C) (i-ii) The average radial trends of pSMAD1 shown as line plots. (i) Line plots shown individually for SB and NODAL conditions. Standard deviations shown in grey, and 95% confidence intervals shown in black. (ii) Line plots represented in the same graph. The p-values were calculated using Mann-Whitney U-test. ^∗^ indicates p<0.0001. D) (i-ii) SB supplementation in the induction medium robustly inhibits Nodal signalling. (i) Geometrically confined hPSC colonies were treated with BMP4 for 24h. Media tested were: ‘**Vehicle**’ indicated SR medium (see Materials and Methods for composition) supplemented with BMP4 and bFGF. ‘**SB**’ indicated vehicle supplemented with μM SB431542. (ii) Gene expression Nodal and Lefty-A (a Nodal target) after 24h of treatment with either ‘Vehicle’ of ‘SB’ media. Data shown as mean ± s.d. (n=3, technical replicates, independent wells). The p-value was calculated using two-sided Student’s t-test.

**Figure S11:**
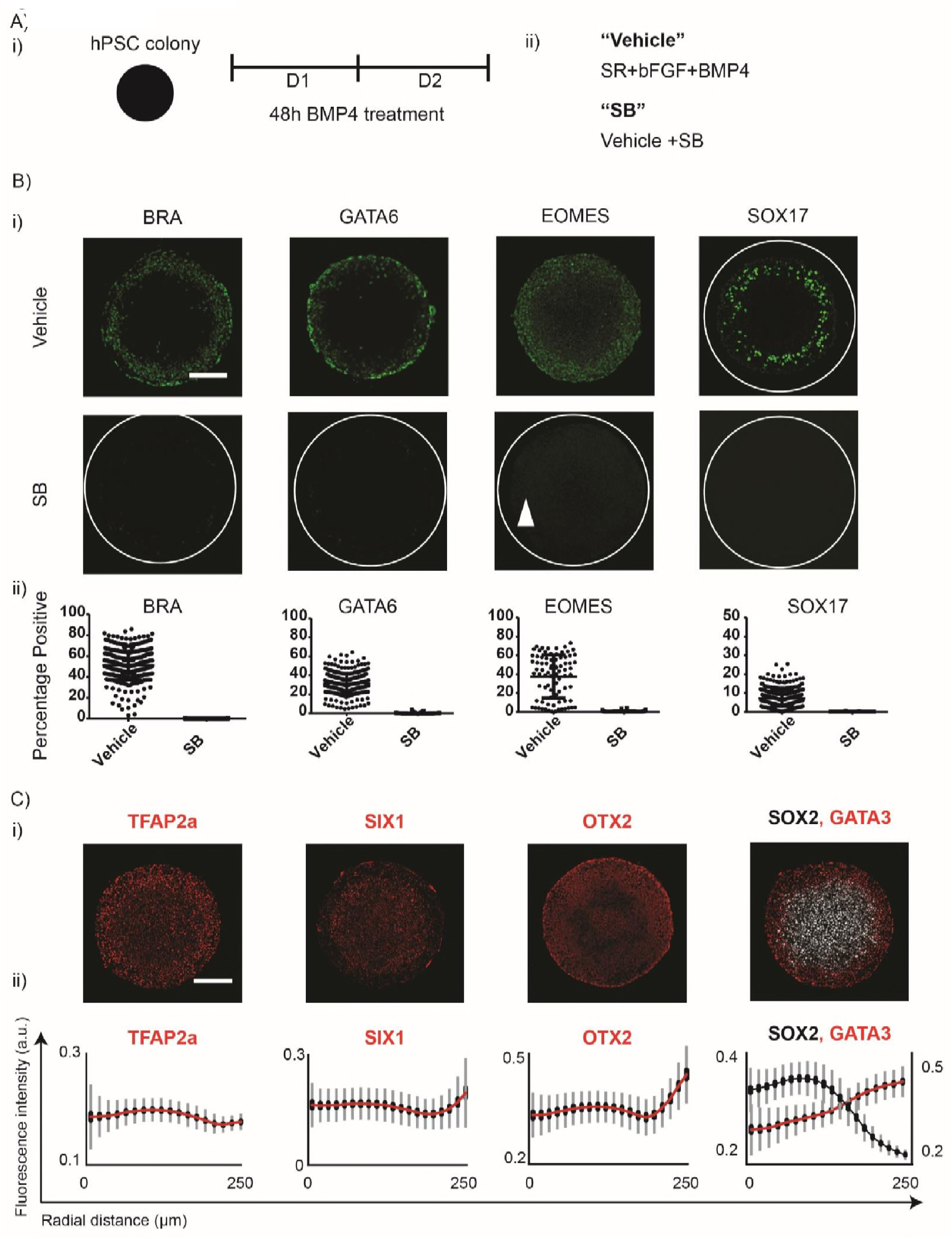
Nodal inhibition during BMP4 treatment of hPSC colonies abrogates peri-gastrulation-associated fates and induces pre-neurulation-associated fates. A) (i-ii) Overview of experimental setup. (i) Geometrically confined hPSC colonies were treated with BMP4 for 48h. (ii) Media tested. ‘**Vehicle**’ indicated SR medium (see Materials and Methods for composition) supplemented with BMP4 and bFGF. ‘**SB**’ indicated vehicle supplemented with μM SB431542. B) (i-ii) Response of gastrulation-associated fate patterning in Vehicle and SB conditions. (i) Representative immunofluorescent images for BRA, GATA6, EOMES, and SOX17 for Vehicle and SB conditions. White triangle represents non-specific background staining for EOMES. Scale bar represents 200μm. (ii) Quantified expression of gastrulation-associated fates. Each data point represents an identified colony. The total number of colonies were (373,248), (325, 317), (81,72), and (506, 329) for BRA, GATA6, EOMES, and SOX17 respectively for (Vehicle and SB treatments). The data are pooled from two experiments except for EOMES, which was performed once. C) (i-ii) Pre-neurulation-like fate patterning observed in the presence of SB. (i) Representative immunofluorescent images of TFAP2A, SIX1, OTX2, and co-stained image of SOX2, and GATA3. Scale bar represents 200μm. (ii) Average radial expression intensity of the pre-neurulation-associated fates represented as line plots. Standard deviation shown in grey, and 95% confidence intervals shown in black.

**Figure S12:**
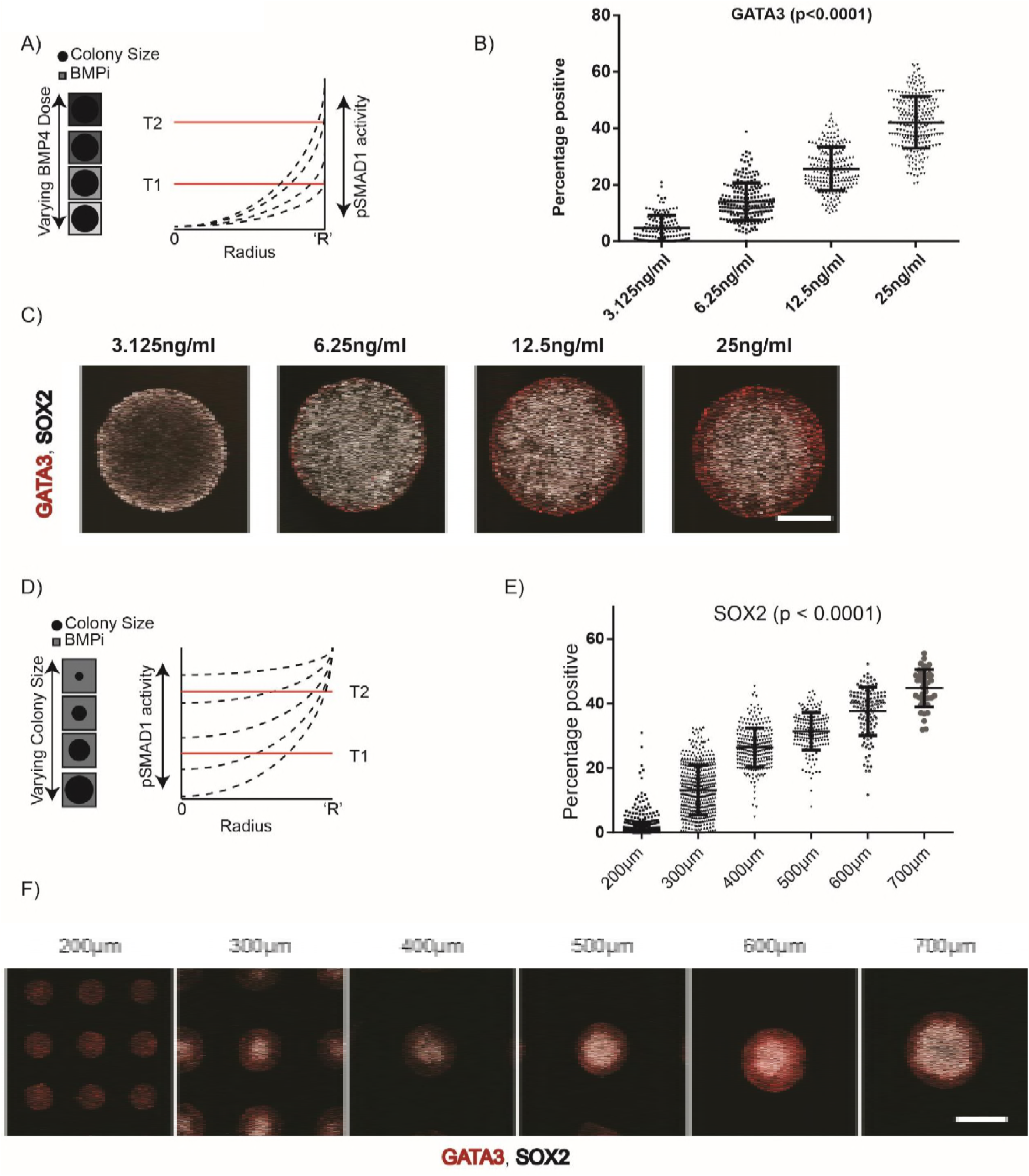
SOX2 and GATA3 expression is consistent with a pSMAD1 dose-dependent fate patterning. A) Overview of experimental setup. Perturbing BMP4 dose in induction medium while maintaining colony size varies pSMAD1 concentration levels at the colony periphery (see **Fig S9A-C** for details). B) Percentage of cells in each identified colony expressing GATA3 when colonies of 500μm in diameter were treated with varying doses of BMP4 (3.125ng/ml, 6.25ng/ml, 12.5ng/ml, and 25ng/ml) in induction medium. Each data point represents an identified colony. The total number of colonies were 131, 208, 215, and 244 for the respective doses. Data pooled from two experiments. Bars represent mean ± s.d. The p-value was calculated using Kruskal-Wallis test. C) Representative immunofluorescent images of colonies stained for SOX2 and GATA3. Scale bar represents 200μm. D) Overview of experimental setup. Perturbing the colony size while maintaining the BMP4 dose constant in the induction medium varies the pSMAD1 levels at the colony center (see **Fig. S9D-E** for details). E) Percentage of cells in each identified colony expressing SOX2 when colonies of varying sizes (200μm, 300μm, 400μm, 500μm, 600μm, and 700μm in diameter) were treated with 25ng/ml of BMP4 in induction medium. Each data point represents an identified colony. The total number of colonies were 932, 439, 256, 175, 122, and 45 for the respective sizes. Data pooled from two experiments. Bars represent mean ± s.d. The p-value was calculated using Kruskal-Wallis test. F) Representative immunofluorescent images of colonies stained for SOX2 and GATA3. Scale bar represents 200μm.

**Figure S13:**
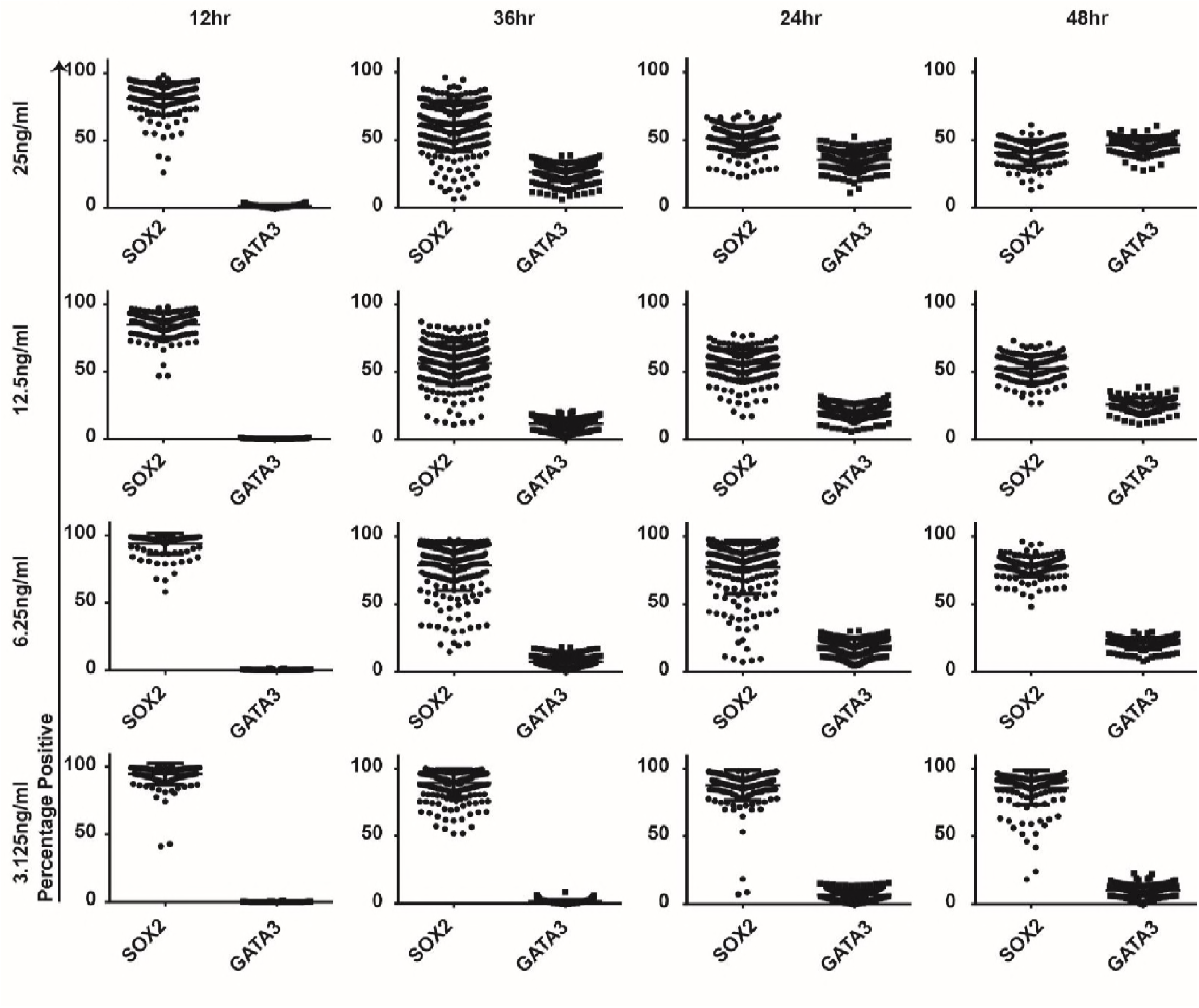
GATA3 expression arises as a function of BMP4 dose and induction time. Percentage of cells expressing SOX2, and GATA in 500μm colonies induced to differentiate at varying concentrations of BMP4 (3.125 ng/ml, 6.25ng/ml, 12.5 ng/ml, and 25 ng/ml) and induction times (12 hours, 24 hours, 36 hours, and 48 hours). Each data point represents an identified colony, and each condition had over 100 colonies. Data pooled from two experiments. Bars represent mean ± s.d.

**Figure S14:**
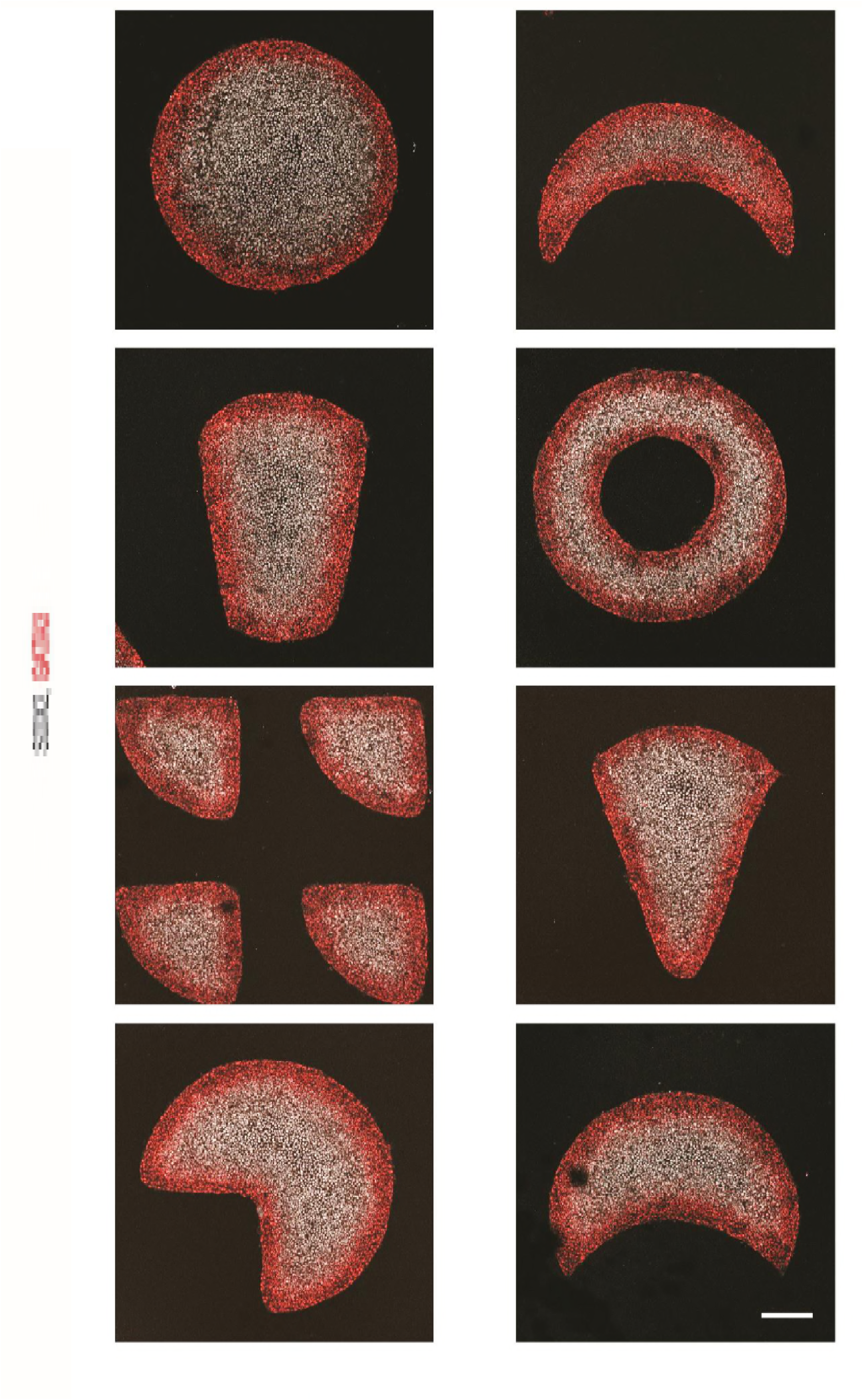
Changing shapes does not affect outside-in spatial patterning. Representative images of various shapes of geometrically-confined hPSC colonies treated with BMP4 and SB in SR medium. Varying colony shapes does not result in any deviation from anticipated fate patterning. The experiment was performed once. Scale bar represents 200μm.

**Figure S15:**
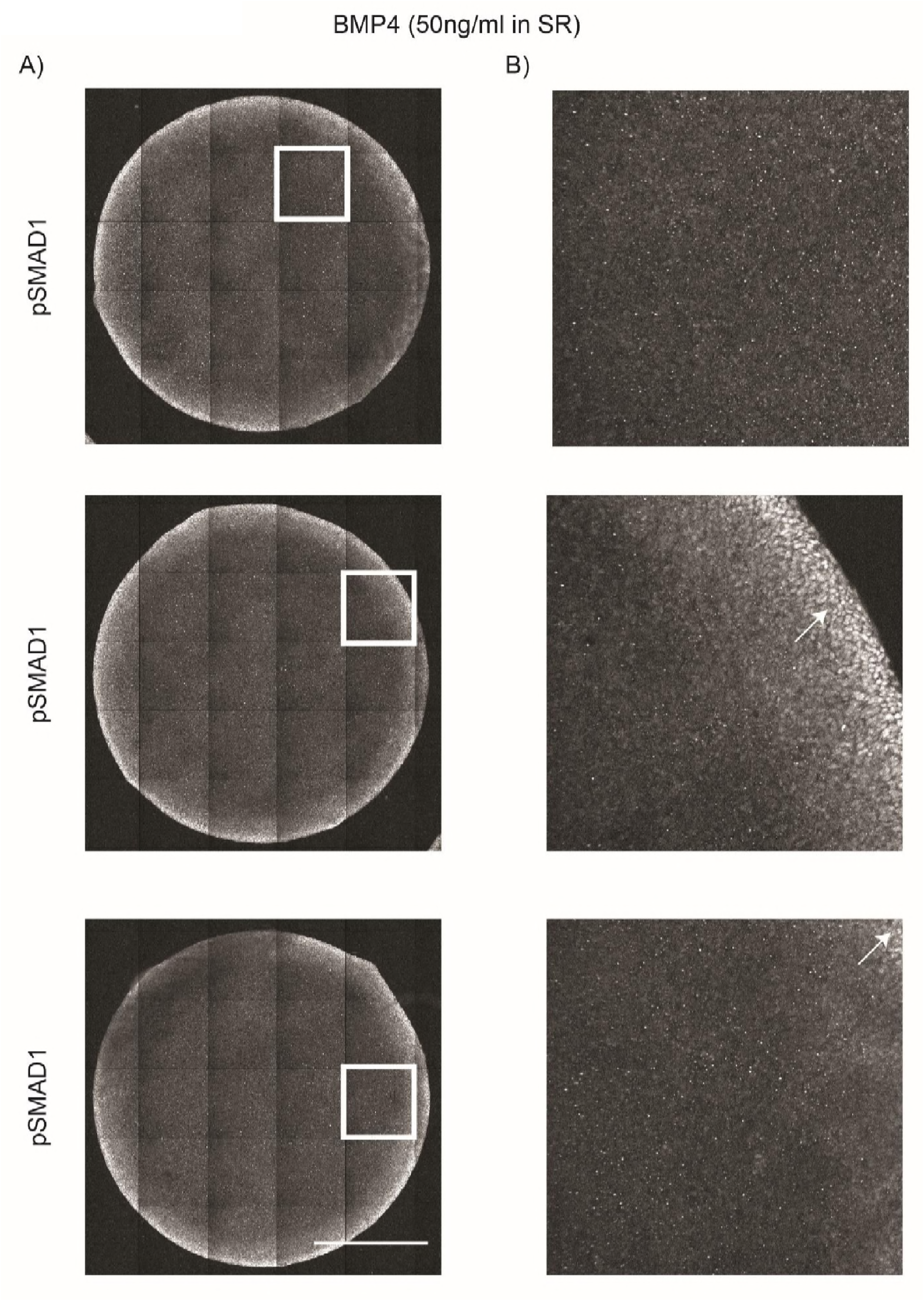
No spatial oscillations of pSMAD1 detected when large geometrically confined hPSC colonies are treated with 50ng/ml BMP4 and SB in SR medium. A-B) No discernable spatial oscillations of pSMAD1 expression detected with geometrically confined hPSC colonies of 3mm diameter were treated with 50ng/ml of BMP4 and SB for 24h in SR medium. A) Stitched images of the entire colony stained for pSMAD1 shown in greyscale for ease of visibility. B) Enlarged fields that are indicated by white squares in A. White arrows indicate regions that contain cells with positive pSMAD1 expression. Scale bar represents 1mm.

**Figure S16:**
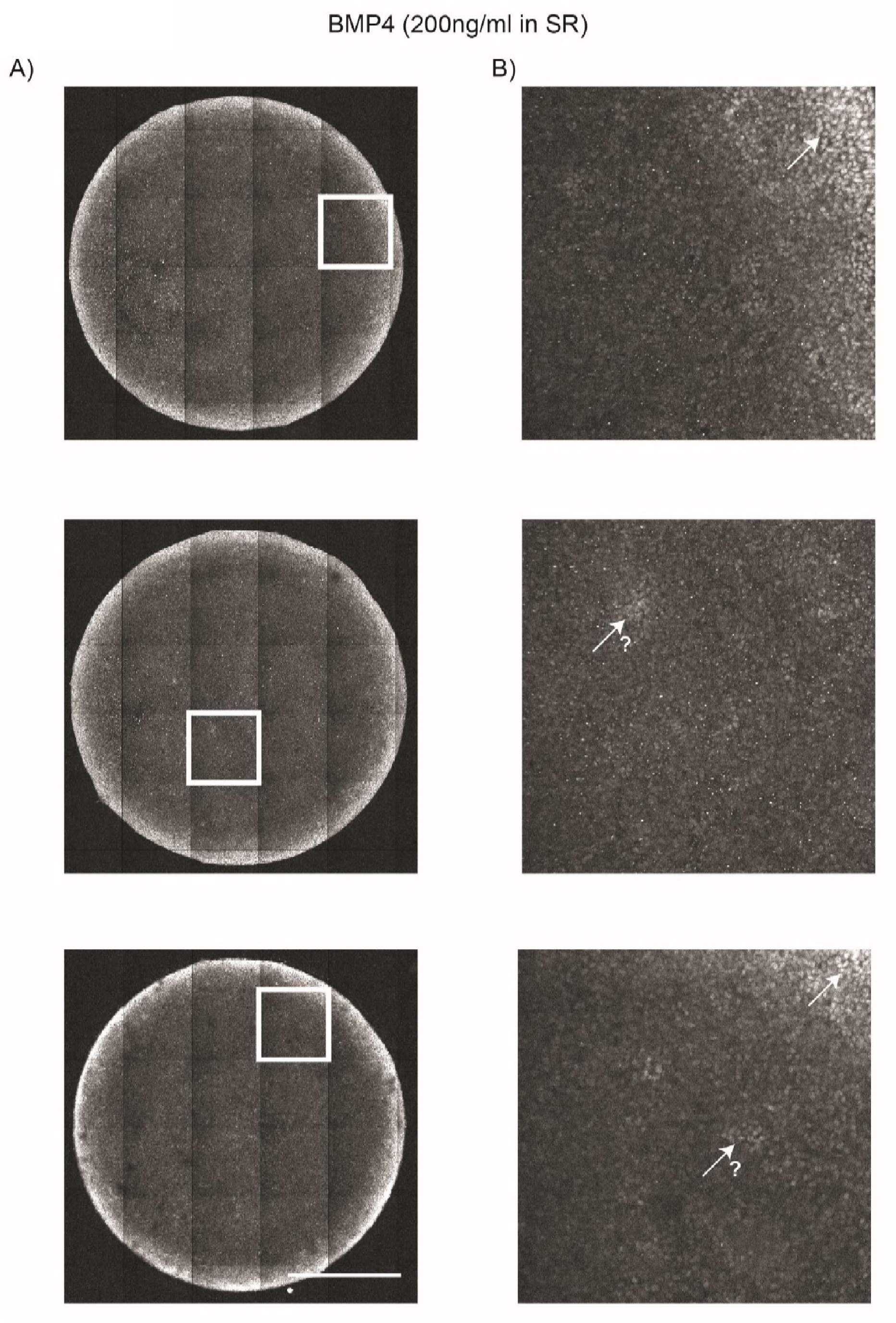
Negligible spatial oscillations of pSMAD1 detected when large geometrically confined hPSC colonies are treated with 200ng/ml BMP4 and SB in SR medium. A-B) Negligible spatial oscillations of pSMAD1 expression detected with geometrically confined hPSC colonies of 3mm diameter were treated with 200ng/ml of BMP4 and SB for 24h in SR medium. A) Stitched images of the entire colony stained for pSMAD1 shown in greyscale for ease of visibility. B) Enlarged fields that are indicated by white squares in A. White arrows at the colony periphery indicate regions that contain cells with positive pSMAD1 expression. White arrows with accompanying question marks indicate regions that possibly show expression of pSMAD1; however, the staining in these regions is inconclusive. Scale bar represents 1 mm.

**Figure S17:**
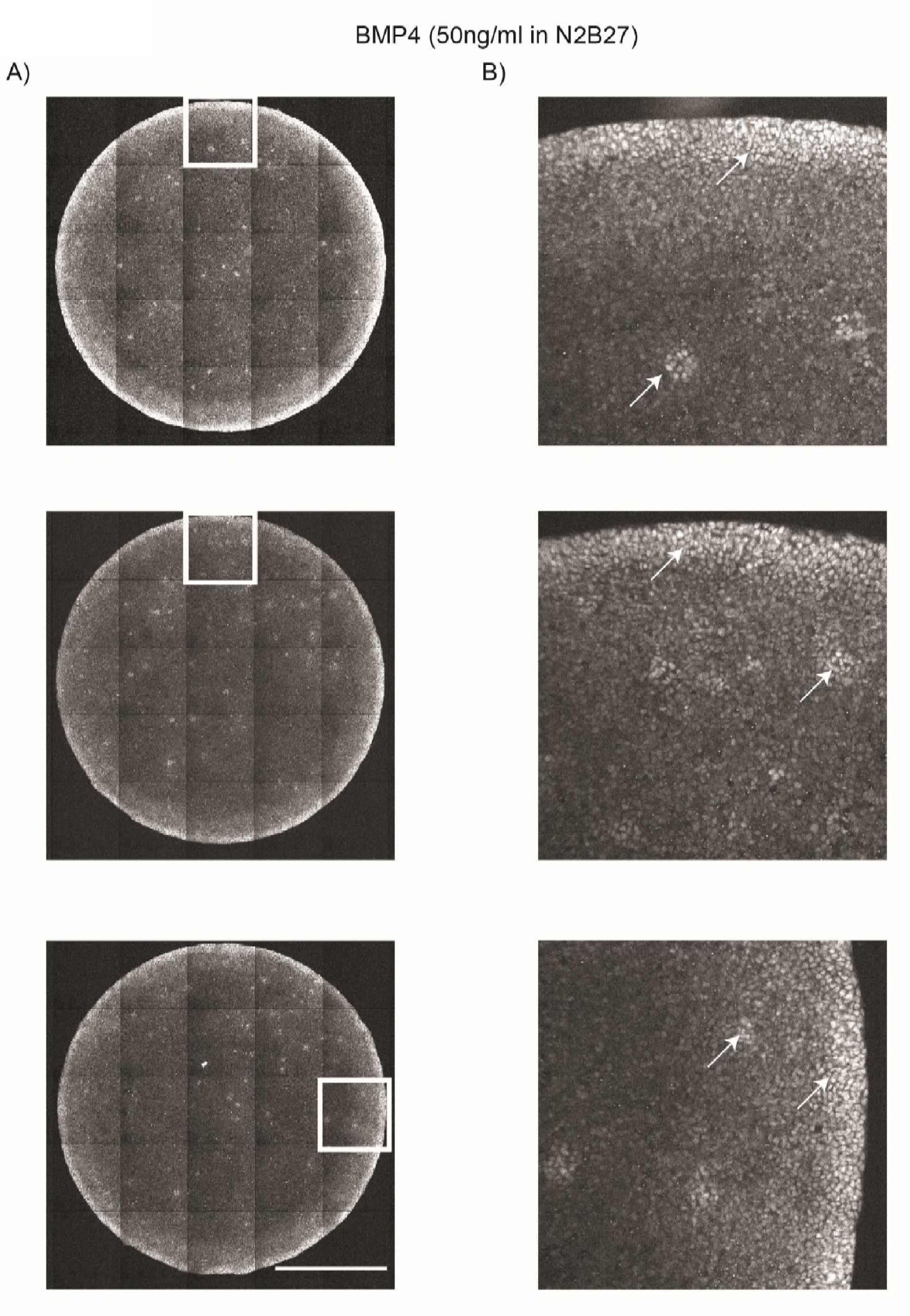
Marginal spatial oscillations of pSMAD1 detected when large geometrically confined hPSC colonies are treated with 50ng/ml BMP4 and SB in N2B27 medium. A-B) Marginal spatial oscillations of pSMAD1 expression detected with geometrically confined hPSC colonies of 3mm diameter were treated with 50ng/ml of BMP4 and SB for 24h in N2B27 medium. A) Stitched images of the entire colony stained for pSMAD1 shown in greyscale for ease of visibility. B) Enlarged fields that are indicated by white squares in A. White arrows indicate regions that contain cells with positive pSMAD1 expression. Scale bar represents 1mm.

**Figure S18:**
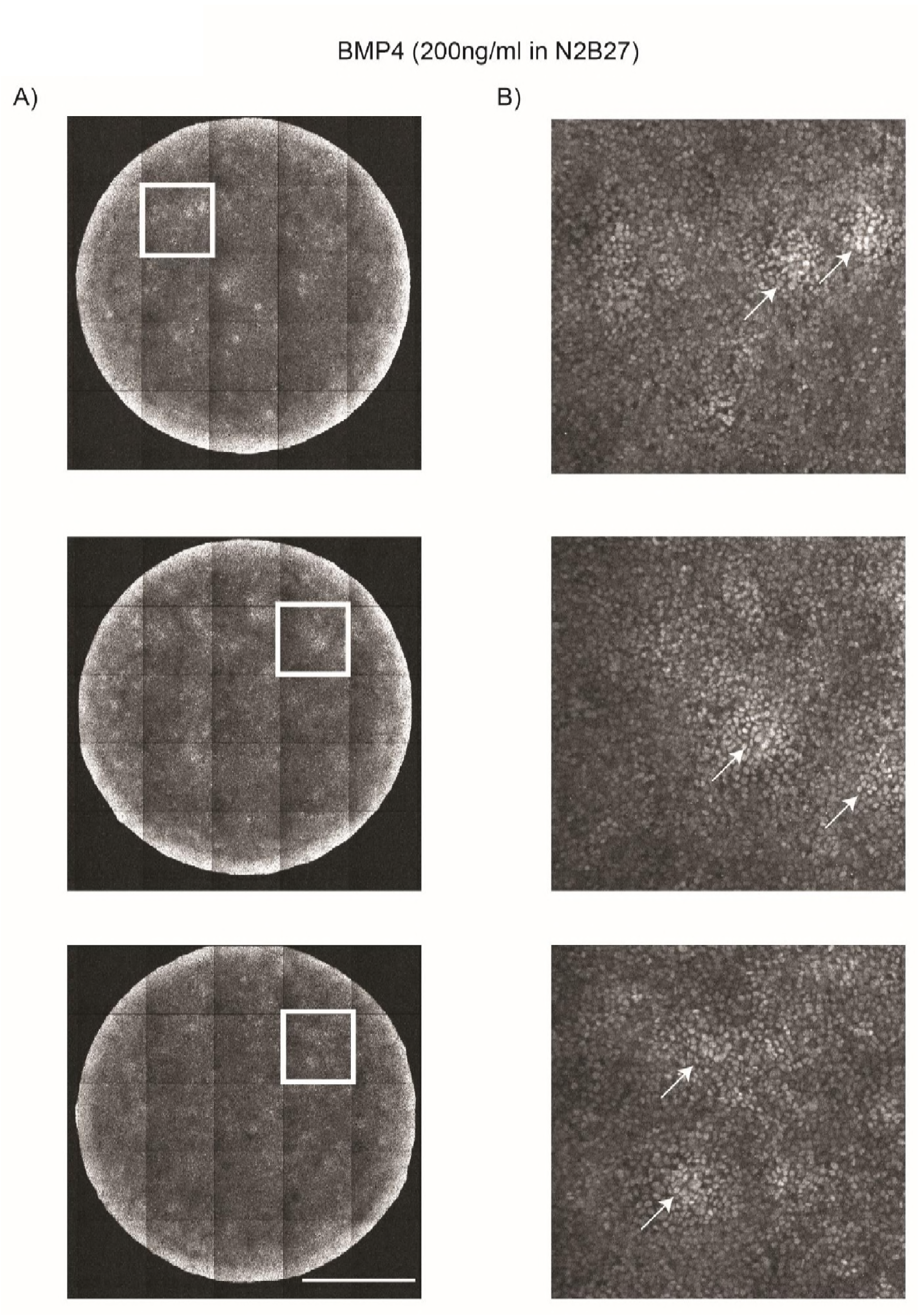
Treatment of large geometrically confined hPSC colonies with 200ng/ml BMP4 and SB in N2B27 medium results in spatial oscillations of pSMAD1. A-B) Spatial oscillations of pSMAD1 expression detected with geometrically confined hPSC colonies of 3mm diameter were treated with 200ng/ml of BMP4 and SB for 24h in N2B27 medium. A) Stitched images of the entire colony stained for pSMAD1 shown in greyscale for ease of visibility. B) Enlarged fields that are indicated by white squares in A. White arrows indicate regions that contain cells with positive pSMAD1 expression. Scale bar represents 1mm.

**Figure S19:**
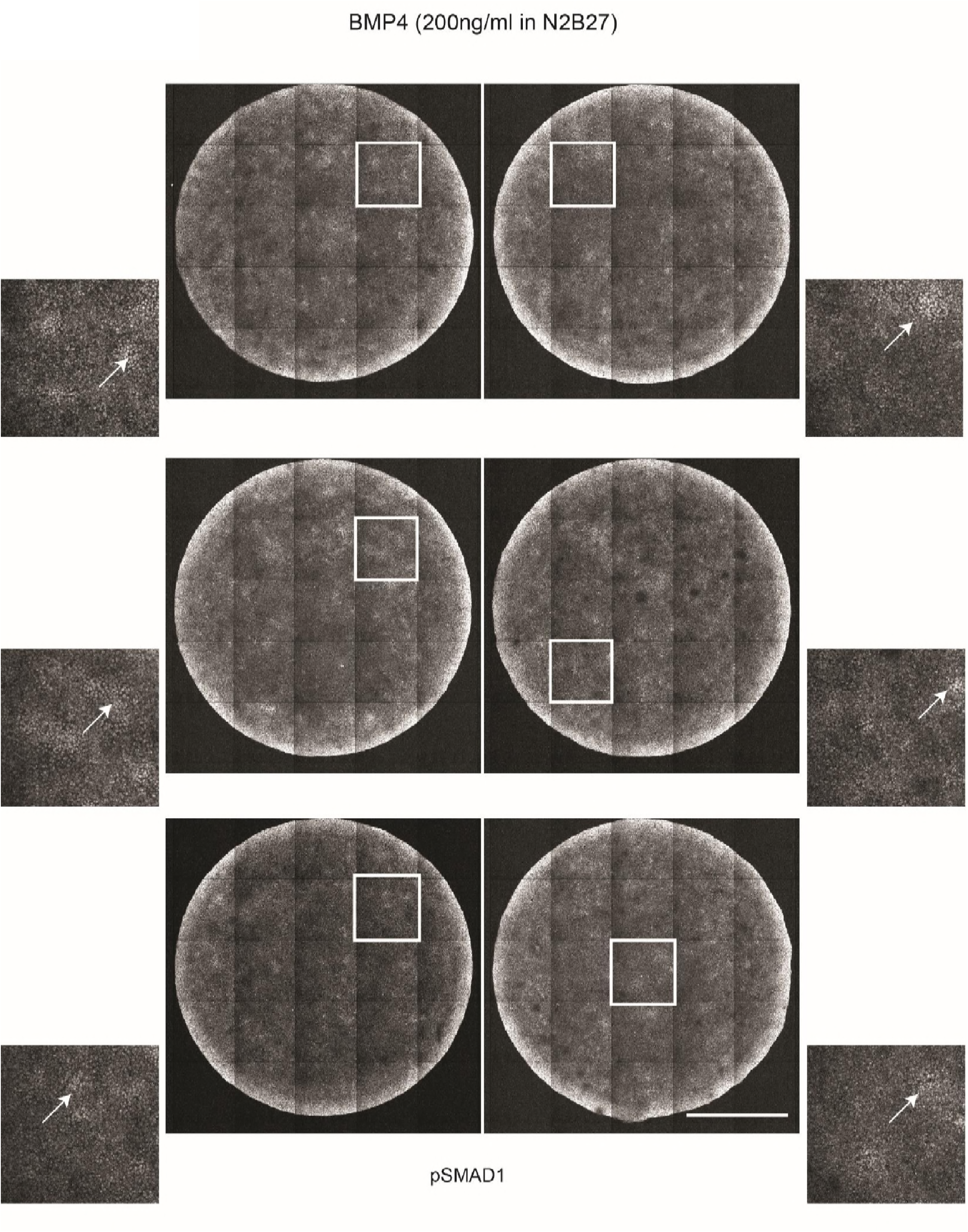
Additional replicates of immunofluorescent images demonstrating oscillatory pSMAD1 expression in the center of large geometrically confined hPSC colonies treated with 200ng/ml BMP4 and SB in N2B27 medium. Additional representative images of 3mm diameter geometrically confined hPSC colonies treated with 200ng/ml BMP4 and SB in N2B27 medium for 24h and stained for pSMAD1. Zoomed-in images of fields contained within white squares shown adjacent to the stitched images. White arrows indicate regions that contain cells with positive pSMAD1 expression. Scale bar represents 1mm.

**Figure S20:**
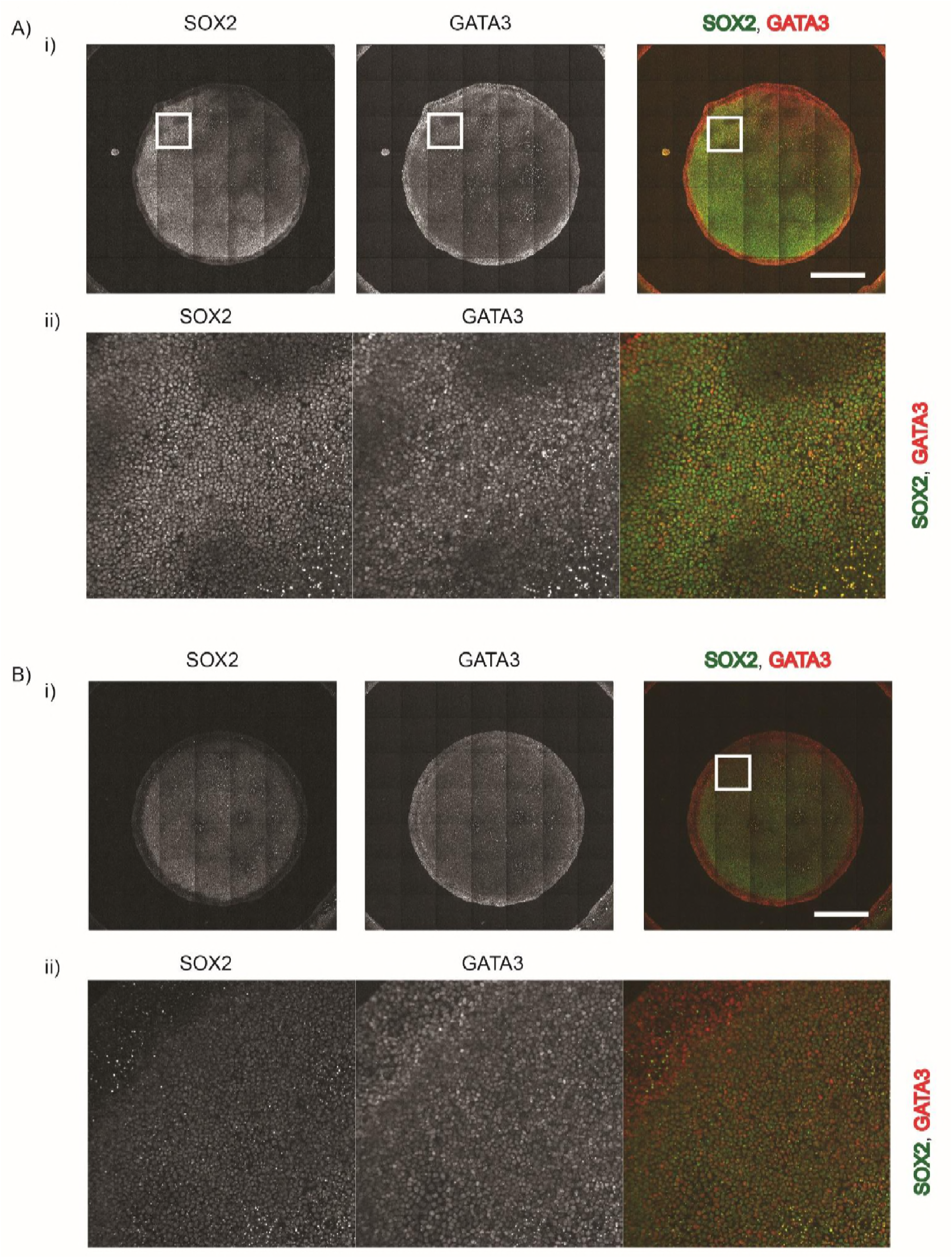
No spatial oscillations of pre-neurulation-like fates detected when large geometrically confined hPSC colonies are treated with 200ng/ml BMP4 and SB in SR medium. A-B) Treatment of geometrically confined-hPSC colonies with 200ng/ml of BMP4 and SB for 48h results in RD-like periodic spatial oscillations of SOX2 and GATA3 expression. i) Representative stitched images of 3mm diameter hPSC colonies differentiated with 200ng/ml of BMP4 for 48h. Scale bar represents 1 mm. ii) Zoomed section outlined by the white square in (i). The experiment was repeated two times.

**Figure S21:**
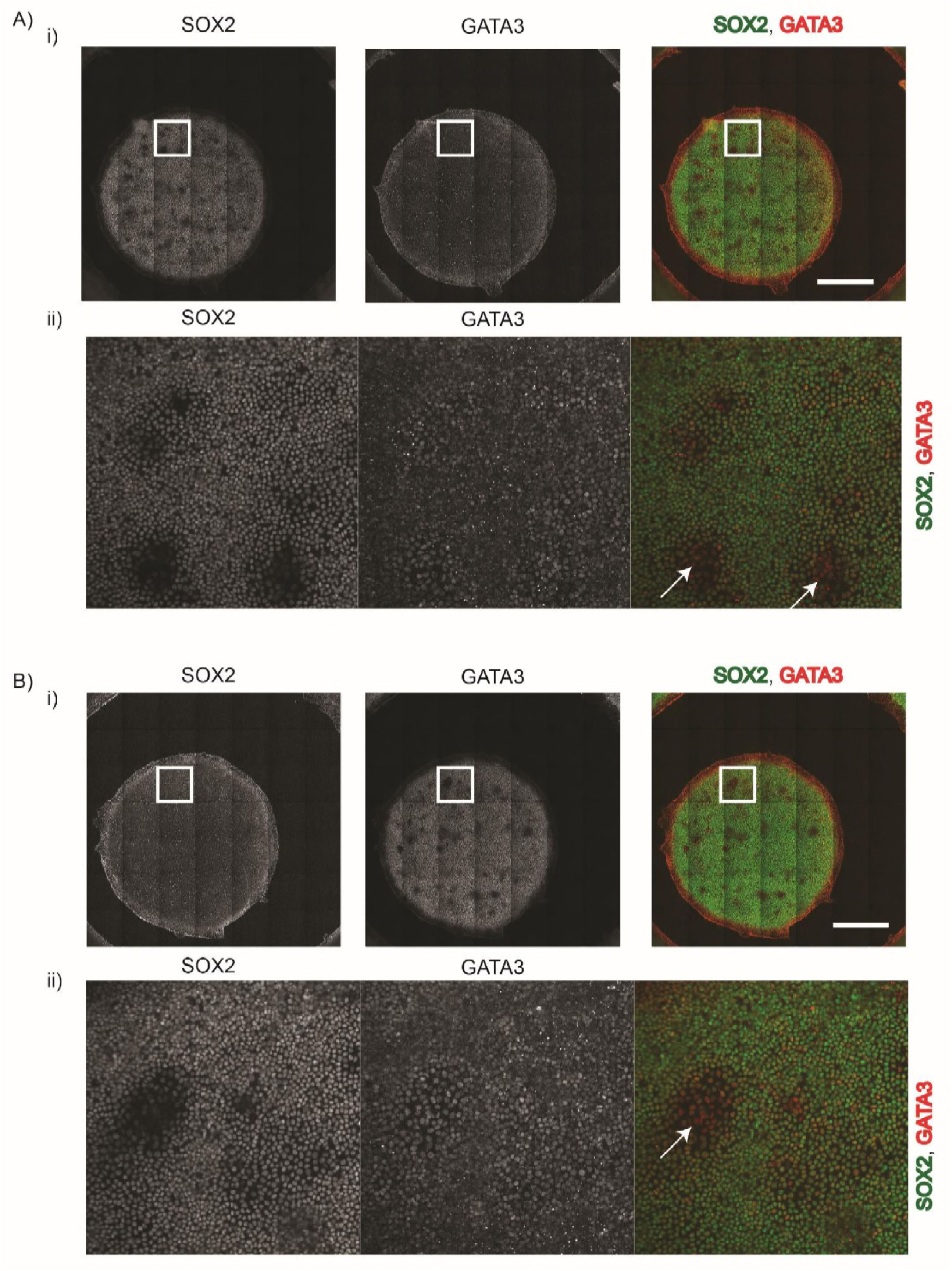
Minor spatial oscillations of pre-neurulation-like fates detected when large geometrically confined hPSC colonies are treated with 50ng/ml BMP4 and SB in N2B27 medium. AB) Treatment of geometrically confined-hPSC colonies with 200ng/ml of BMP4 and SB for 48h results in RD-like periodic spatial oscillations of SOX2 and GATA3 expression. i) Representative stitched images of 3mm diameter hPSC colonies differentiated with 200ng/ml of BMP4 for 48h. Scale bar represents 1mm. ii) Zoomed section outlined by the white square in (i). The experiment was repeated two times.

**Figure S22:**
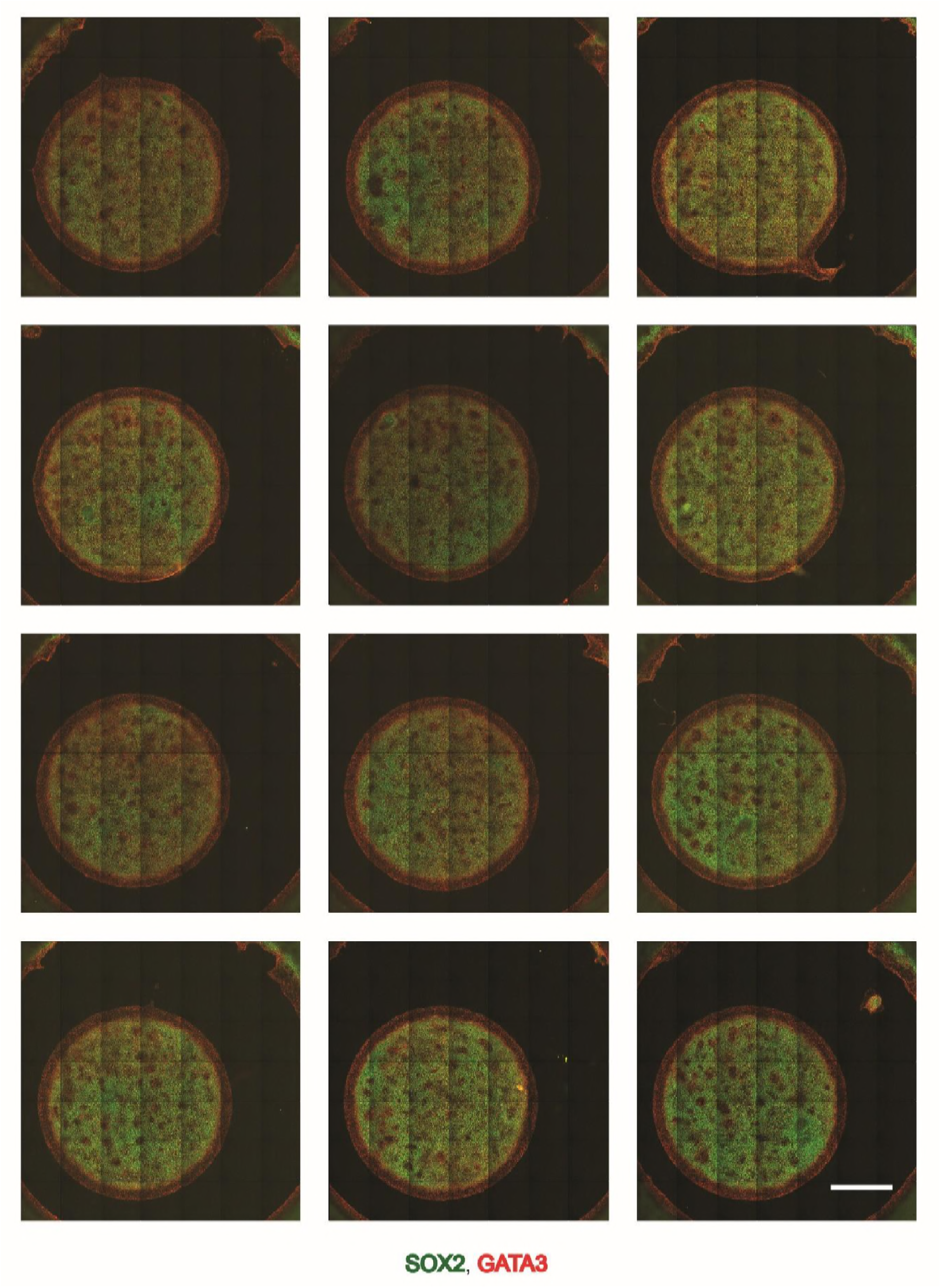
RD-like spatial oscillations of pre-neurulation-like fates detected when large geometrically confined hPSC colonies are treated with 200ng/ml BMP4 and SB in N2B27 medium. Representative immunofluorescent images of geometrically confined hPSC colonies of 3mm diameter stained for SOX2, and GATA3. The colonies were treated with 200ng/ml of BMP4 and SB for 48h. Scale bar represents 1mm.

**Sup. Table 1:**
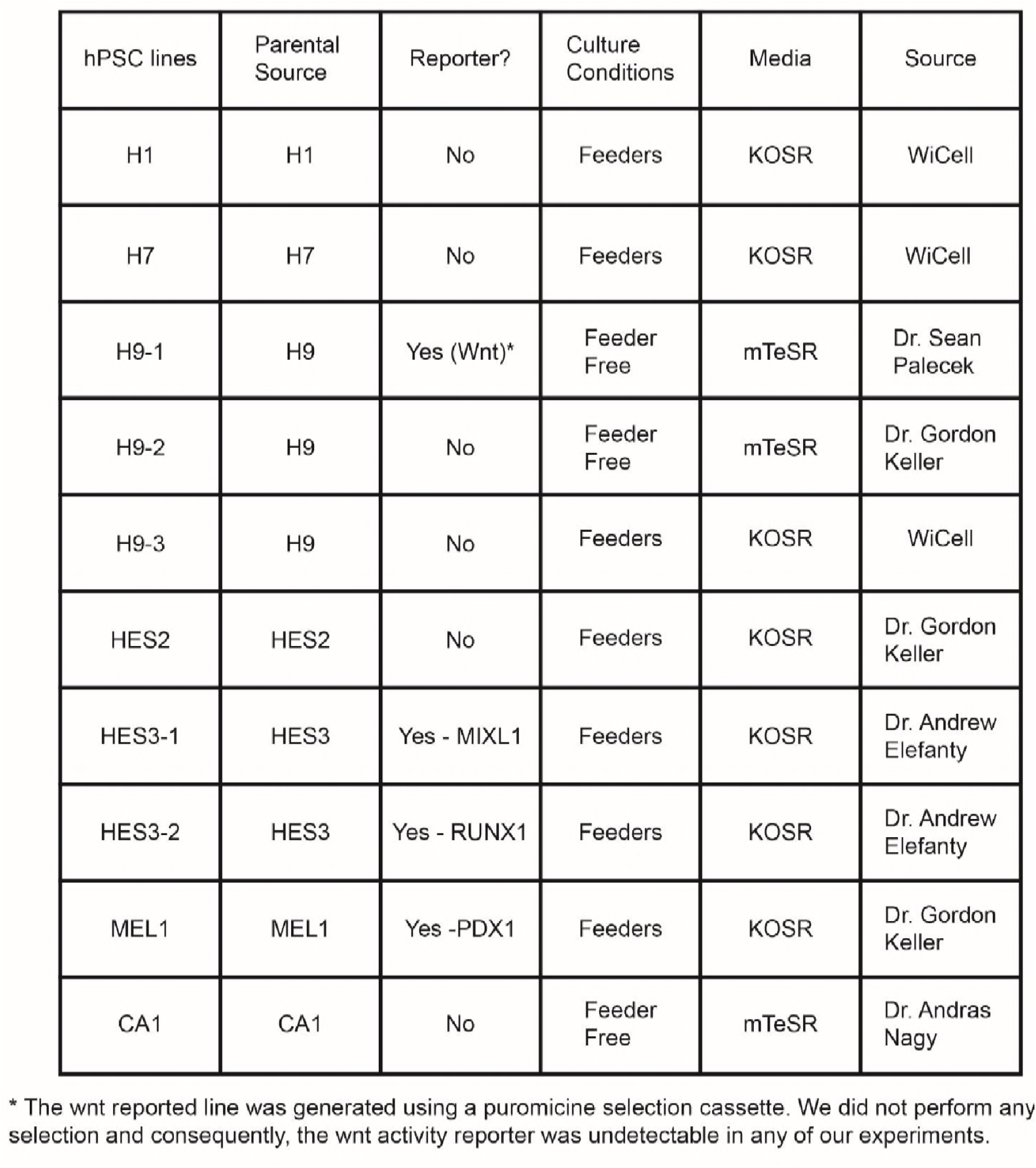
Human PSC lines utilized in this study: Complete list of hPSC lines used for the study; and the culture conditions employed for their maintenance.

**Sup. Table 2:**
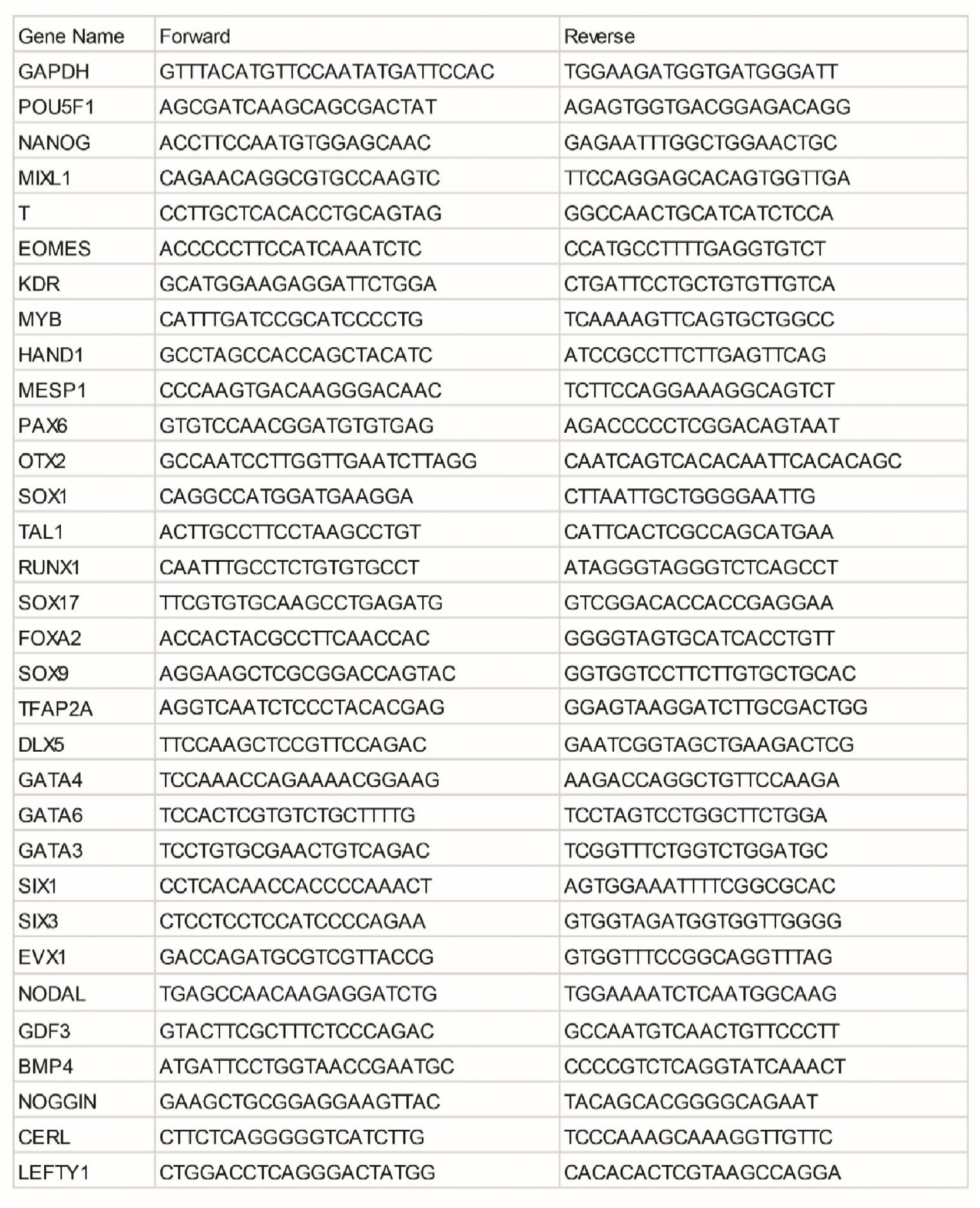
List of primers used in this study

**Sup. Table 3:**
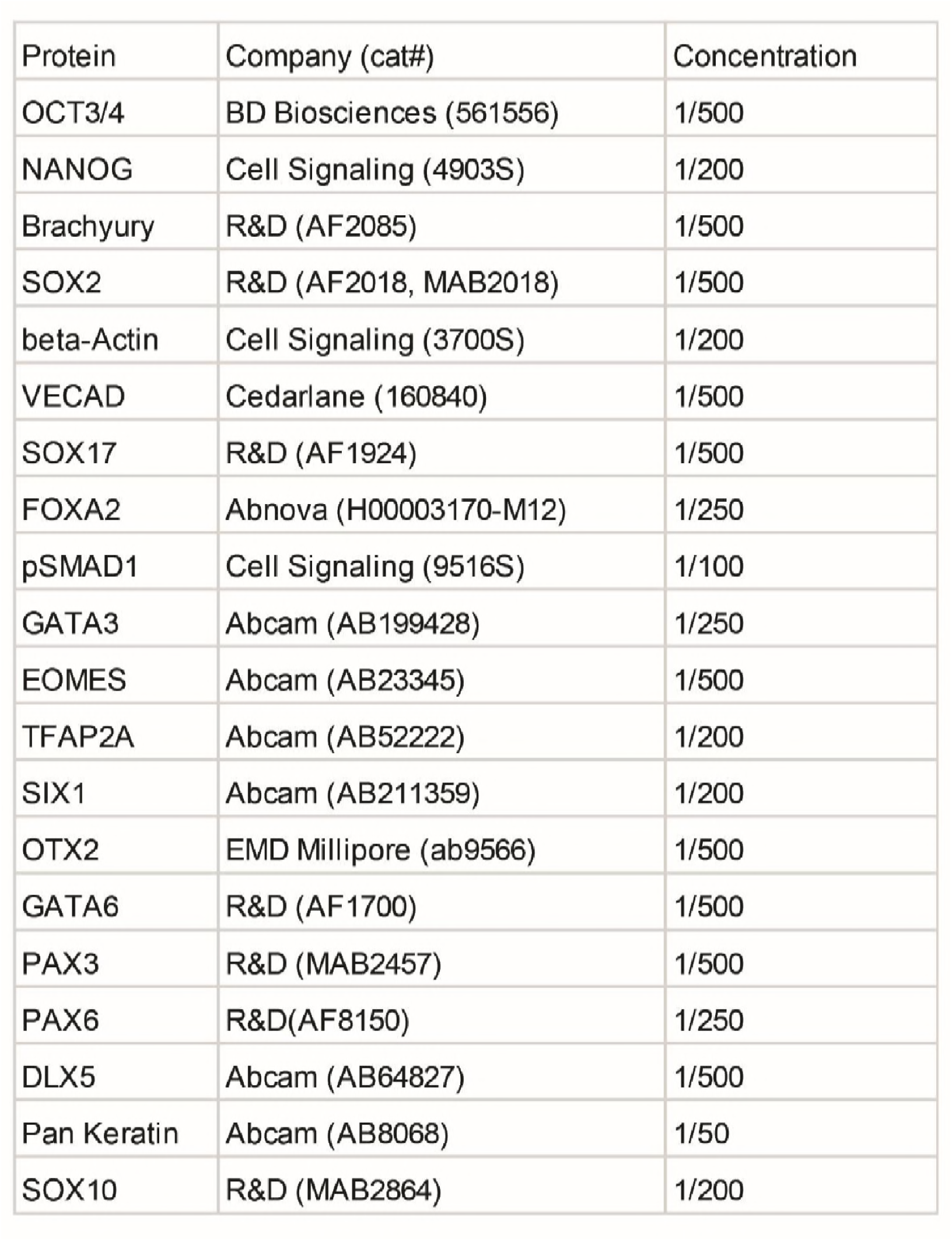
List of antibodies used in this study

## Acknowledgements

We kindly thank Dr. Andras Nagy for providing the CA1 human embryonic stem cell line; Dr. Sean Palecek for providing the 7TGP(H9) line; Dr. Gordon Keller for providing the GKH9(H9), HES2, and the PDX1-GFP(MEL1) lines; and Dr. Andrew Elefanty for providing the RUNX1-GFP(HES3), and the MIXL1-GFP(HES3) lines. We thank Dr. Emanuel Nazareth for providing us with the micro-contact printing stamps. We gratefully acknowledge Dr. Alfonso Martinez Arias, Dr. James Briscoe, Dr. Teresa Rayon Alonso, Dr. Christopher Demers, and Dr. Celine Bauwens for providing insightful feedback to our manuscript. This work was funded by the Canadian Institutes for Health Research and Medicine by Design, a Canada First Research Excellence Program at the University of Toronto. MT received funding from CREATE Materials, Mimetics, and Manufacturing from the Natural Sciences and Engineering Research Council of Canada. DD is a PhD fellow of Research Foundation - Flanders (FWO). FMW gratefully acknowledges funding from the UK Medical Research Council and Wellcome Trust. PWZ is the Canada Research Chair in Stem Cell Bioengineering.

